# Functional Characterization of RSV Clades Associated with Prophylactic Breakthrough Infections in Pediatric Transmission Clusters

**DOI:** 10.64898/2026.05.29.728867

**Authors:** Estefany Rios-Guzman, Ria Almohtadi, Seth H. Borrowman, Tien Doan, Charlie R. Boyle, Dulce S. Garcia, Margarita Rzhetskaya, Lacy M. Simons, Jacob W. Class, Alexa Mendoza, Molly Schnieders, Anna Pawlowski, Sameer J. Patel, Ramon Lorenzo-Redondo, Judd F. Hultquist

## Abstract

The recent development of new respiratory syncytial virus (RSV) prophylactics for the prevention of severe lower respiratory tract infections in infants and older adults promises in lowering disease burden in these vulnerable populations. However, it remains unclear if periodic breakthrough infections in these populations may drive the emergence of resistant isolates or clades and what factors might contribute to these breakthroughs. In this retrospective cohort study, we performed whole-genome sequencing of RSV isolates from infants and adults during the past two RSV seasons (2023-2025) to assess viral and clinical correlates of nirsevimab breakthrough. RSV infections from nirsevimab breakthrough cases were associated with less severe clinical outcomes in the first, but not second, season after administration. While breakthrough isolates did not share any Fusion glycoprotein mutations in predicted antigenic sites, they largely belonged to only a few circulating clades that were responsible for driving temporally distinct pediatric transmission clusters. To determine if these transmission clusters and breakthrough infections were in part driven by differences in the Fusion proteins of these clades, we compared the relative fusogenicity and neutralization susceptibility of Fusion proteins from contemporary circulating clades. Notably, RSV-A clade A.D.3 exhibited modestly reduced susceptibility to nirsevimab neutralization, though it wasn’t associated with any transmission clusters or breakthrough infections. Collectively, these data suggest that clade associations with prophylactic breakthrough are driven by pediatric transmission clusters rather than clade-associated resistance, though continued surveillance will be vital as prophylactic coverage continues to rise.

## INTRODUCTION

Respiratory Syncytial Virus (RSV) is a leading cause of acute respiratory tract infections in infants and poses a significant health risk for older adults and immunocompromised populations.^1^ Nearly all children in the United States are infected with RSV by age two, and the virus is responsible for over 3 million hospitalizations worldwide each year, underscoring its substantial disease burden.^2,3^

To address this global health challenge, pharmaceutical efforts resulted in the recent development and FDA approval of several new prophylactic agents for the prevention of severe disease in pediatric populations. These include the maternal vaccine, Abrysvo (2023), and two monoclonal antibodies (mAbs) nirsevimab (2023) and clesrovimab (2025), all of which target the RSV Fusion (F) glycoprotein.^4,5,6^ The RSV F glycoprotein mediates viral entry by driving membrane fusion and is the primary target of all currently available prophylactic monoclonal antibodies and vaccines.^7,8,9^ While these interventions represent a major advancement in RSV prophylaxis, they also raise concerns about added selective pressure on the virus and the potential emergence of resistance mutations on the F glycoprotein.^10, 11^ Failures in earlier RSV mAb clinical trials highlight the need for ongoing genomic surveillance and functional characterization. In 2017, the mAb suptavumab failed to meet clinical endpoint in phase 3 trial after emergent resistance mutations in RSV-B F (L172Q and S173L) became dominant and drastically reduced protective efficacy.^12^

In recent seasons, collective efforts to expand RSV genomic surveillance networks have yielded new insights into RSV evolutionary trends.^13^ , ^14, 15^ During the COVID-19 pandemic, reduced RSV exposure and altered seasonality due to non-pharmaceutical interventions (NPIs) shifted transmission patterns and RSV genetic diversity.^16,17^ Since then, these same networks have been used to monitor prophylactic efficacy and to assess the risk of emergent resistance.^18,19^ While highly effective at preventing the onset of severe disease, periodic breakthrough infections in pediatric populations have been reported for both nirsevimab and Abrysvo, though these have not been linked to any widespread resistance mutation.^10,20,21,11^ As prophylactic coverage continues to increase, the factors driving these breakthrough infections, and the potential they may drive the emergence of resistant isolates or clades, remains a critical unknown. To address this question, we performed a retrospective cohort study to investigate the incidence, clinical characteristics, and viral genotypes associated with nirsevimab breakthrough infection. Breakthrough infections were associated with mild disease outcomes in the first, but not second, season after administration. While they were not associated with any known or predicted resistance mutation in the F glycoprotein, they frequently occurred in the context of pediatric-driven transmission clusters dominated by a subset of clades. Cell entry and antibody neutralization assays comparing currently circulating contemporary isolates showed slight resistance of RSV-A clade A.D.3 to nirsevimab, though this clade was not associated with any of the breakthrough cases or transmission clusters. Collectively, these data suggest that clade associations with prophylactic breakthrough is driven by pediatric transmission clusters rather than emergent resistance, though continued surveillance and functional phenotyping will be necessary as the viral population continues to evolve in the context of heightened prophylactic rollout.

## RESULTS

### RSV population structures in the US did not globally shift since prophylactic rollout in 2023

To characterize RSV viral population dynamics since the rollout of new prophylactics in 2023, we conducted a retrospective cohort study among adult and pediatric patients who tested positive for RSV via PCR diagnostic tests at Northwestern Medicine (NM) hospital systems and Lurie Children’s Hospital (LCH) in Chicago, IL. From the NM system, we collected 209 RSV-positive residual nasopharyngeal swab diagnostic specimens from June 2023 through February 2025. All of these were processed for whole-genome sequencing using an amplicon-based strategy as previously reported^22^, of which 80 isolates yielded whole-genome sequences. 54 specimens (55%) were from children 18 years of age or younger and 36 (45%) were from adults (**Supplementary Table 1**). A majority (92.5%) of patients experienced an RSV-B infection, predominantly during the 2023-2024 season and a large majority of patient encounters (90%) were outpatient.

From LCH, we collected 1,235 RSV-positive residual diagnostic specimens from November 2024 to April 2025, from which we sub-sampled 591 specimens by age, sex, and epidemiological week for sample processing and sequencing. Of these, 340 isolates yielded whole-genome sequences, a majority of which were RSV-A (85%) (**Supplementary Table 1**). Infants and children less than 2 years of age yielded most of the sequenced isolates (55.6%) with children aged 2 to 18 (44.6%) comprising a majority of the rest. Similar to the clinical outcomes observed for our NM patient encounters, many patients did not require inpatient hospital care at LCH (84.7%). Overall, 800 specimens were processed for sequencing, yielding 420 genomes across both study sites.

All genomes were assigned a clade based on the G-glycoprotein nomenclature in addition to the newly unified classification system that accounts for whole-genome heterogeneity.^23^ All RSV-A (n = 312) and RSV-B (n = 108) samples belonged to G clades GA2.3.5 and GB5.0.5a, respectively. By whole-genome clade designation, RSV-A exhibited expansion of several A.D.1, A.D.3, and A.D.5 subclades across the 2023-2024 and 2024-2025 seasons (**Figure 1A**), while RSV-B showed a strong predominance of clade B.D.E.1 with lower representation of other subclades (**Figure 1B**). While most clades were represented in both the pediatric and adult populations, large expansions of genetically similar isolates in pediatric populations were readily apparent, particularly for clades A.D.1, A.D.5.1, A.D.5.3 and B.D.E.1.2 (**Figure 1A, B**). To visualize this more clearly, RSV isolates from pediatric and adult specimens were stratified on separate phylogenetic trees and amended with NM specimens from prior seasons and again the expansion of highly similar isolates appeared specific to the pediatric populations (**Supplementary Figure 1**). To determine whether the homogeneity in these clades was present in a broader population, we expanded our phylogenetic analysis to include all publicly available RSV whole genome sequences from June 2023 to March 2025 in the United States (US) (**Figure 1C, D**). While our specimens were largely dispersed and reflected the population structure of RSV in the US, some of the more homogenous clusters were composed almost exclusively of specimens from our study.

**Figure 1.**
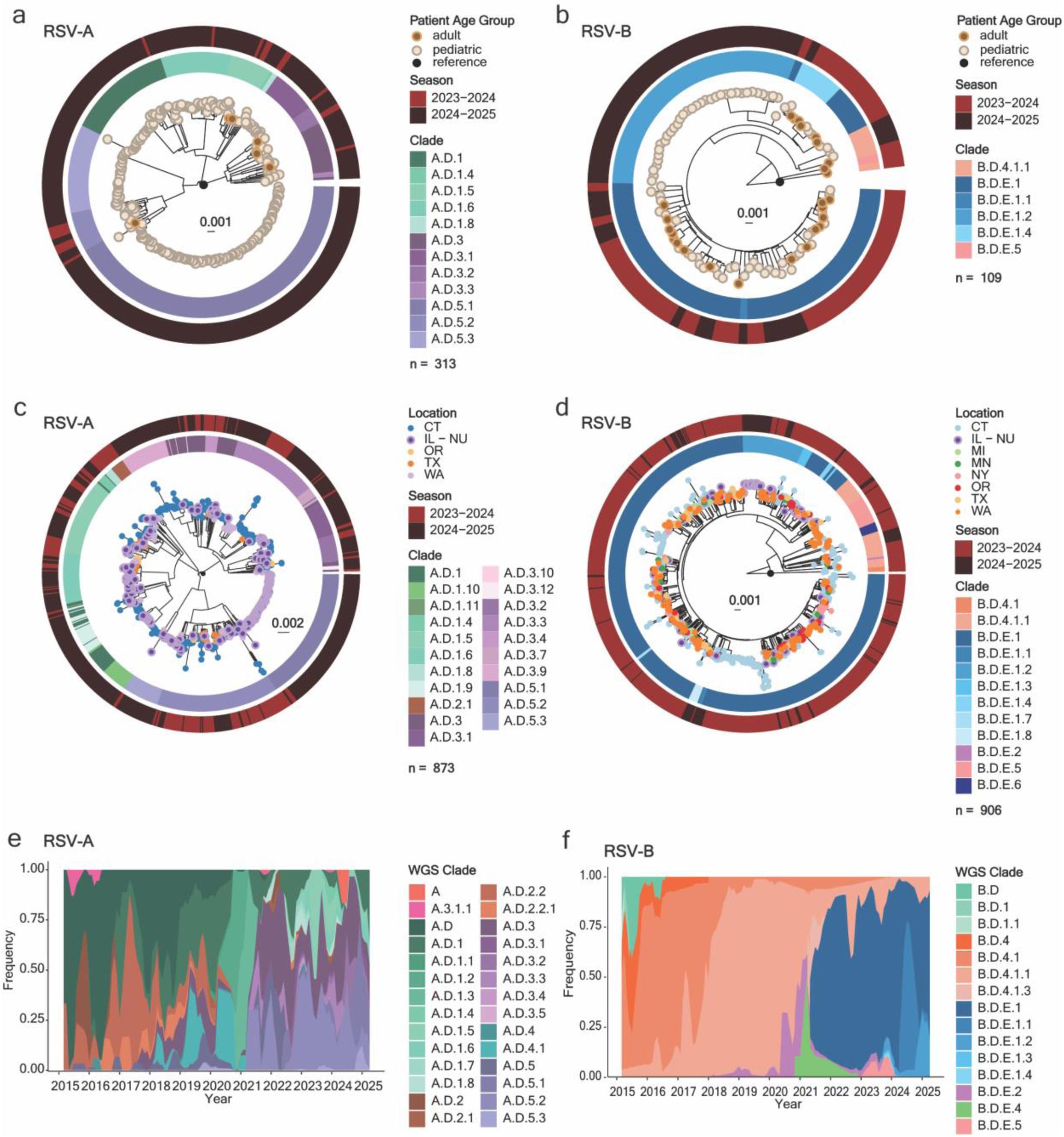
| Phylogenetic analysis of RSV-A and -B whole genome sequences. **a,** Maximum likelihood (ML) phylogenetic analysis of RSV-A genomes from Northwestern Medicine (NM) and Lurie Children’s Hospital (LCH) in Chicago, Illinois, from the 2023 to 2025 RSV seasons (n = 313). **b,** Equivalent ML phylogenetic analyses of RSV-B whole genome sequences from NM and LCH from the 2023 to 2025 RSV season (n = 109). **c,** ML phylogenetic analysis of RSV-A genomes from NM with amended publicly available genomes from the United States (US) collected between the 2023-2025 RSV seasons (n = 873). **d,** ML phylogenetic analysis of RSV-B whole genome sequences from NM and LCH in Chicago, Illinois, with publicly available genomes from the US collected between the 2023-2025 RSV seasons (n = 908). For in-house RSV sequence phylogeny, branch tips are colored by patient age group. For aggregated in-house and US-based RSV sequences, the branch tip is colored by geographical origin of sequence. For all trees, the inner ring is colored by Nextclade v3.21.0 WGS clade designation, and the outer ring indicates the RSV season from which the isolate is from. **e,** Cumulative distribution of RSV-A clade frequency from March 2015 to March 2025 of NM, LCH, and US genomes using a 3-month rolling average. **f,** Cumulative distribution of RSV-B clade frequency from March 2015 to March 2025 of NM, LCH, and US genomes using a 3-month rolling average.

The 2023-2024 season was predominantly driven by RSV-B while the 2024-2025 season was predominantly driven by RSV-A and mirrors the more dramatic swings in subtype observed locally since the COVID-19 pandemic (**Supplementary Figure 2A**). To observe larger-scale trends in RSV clade distribution in the US over the past decade, we visualized the proportional distribution of clades within each subtype as a 3-month rolling average from March 2015 to March 2025. As previously documented, the RSV-A clades A.D.3 and A.D.5 supplanted A.D.1 and diverged into multiple subclades after 2021 (**Figure 1E**). Similarly, RSV-B clade B.D.E.1 superseded B.D.4.1.1 and has since diversified into several subclades (**Figure 1F**). Again, the clade distributions before and during the COVID-19 pandemic appear starkly different from those that follow, though no such selective sweep is apparent in recent years since prophylactic rollout. Notably, the relative percent of pediatric specimens that yielded a whole genome sequence was higher than that for adult specimens, in large part due to the overall higher viral load in these specimens (**Supplementary Figure 2B**). To better visualize this, the distribution of cycle threshold (Ct) values in collected specimens was visualized by age range (**Supplementary Figure 2C**). In general, infants and kids younger than 5 years of age had significantly lower Ct values (higher viral loads) than older age groups (**Supplementary Figure 2D**), though we were unable to control for time since symptom onset was not collected in this study.

Overall, the population structure of RSV-A and RSV-B has seen only minor shifts since the COVID-19 pandemic and the rollout of RSV prophylactics. Similar clade distributions are observed in adult and pediatric populations, but the RSV population structure in pediatric populations is characterized by the expansion of genetically homogenous clusters.

### Temporally distinct RSV transmission clusters are uniquely observed in pediatric populations

Adult under-sampling in RSV genome sequencing has historically obstructed efforts to evaluate larger-scale transmission patterns and genetic differences between age groups. Since our study cohort includes both adult and pediatric patients, we generated a Bayesian time-scaled phylogeny tree modeling RSV transmission events within and between these age groups (**Figure 2A**). Initially, we observed 4 RSV-A and 1 RSV-B pediatric-exclusive transmission clusters, defined by more than 15 sequences with statistical genetic distance cutoffs (TN93 values for RSV-A: 1.1*10^-3^ and RSV-B: 2.99*10^-3^ substitutions per site). Each cluster was uniquely comprised of a singular RSV clade (Cluster 1A: A.D.5.1, Cluster 2A: A.D.5.3, Cluster 3A: A.D.1, Cluster 4A: A.D.3, & Cluster 1B: B.D.E.1.2). Transmission potentials quantified using basic reproductive numbers (R_0_) showed a rapid and acute transmission pattern among RSV-A clusters 2A, 3A, 4A, and RSV-B cluster 1B in the 2024-2025 winter months. Conversely, RSV-A cluster 1A, which is comprised exclusively of the predominating A.D.5.1 clade, exhibited a sustained and cumulative transmission pattern throughout the season, indicating differences in transmission potential among contemporary RSV clades (**Figure 2B**).

**Figure 2.**
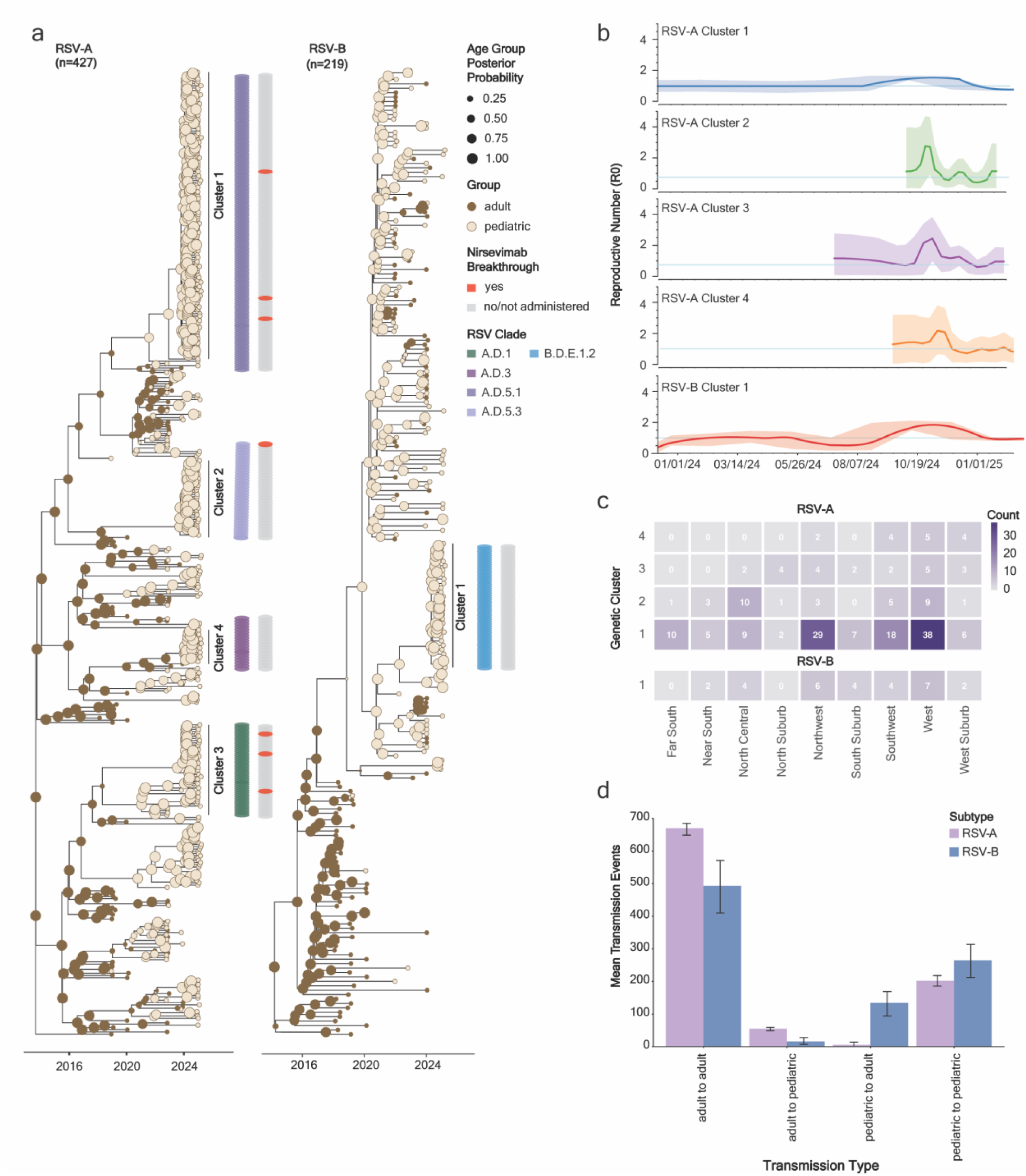
| RSV transmission dynamics between pediatric and adult populations. Bayesian phylogenetic temporal tree of circulating **a,** RSV-A (n=427) and RSV-B (n=219) complete genome sequences sampled in NM and LCH systems as of March 2025. Each tree corresponds to sequences from patient encounters with known age groups depicted by tip color, and the most probable sampled population (pediatric vs. adult) depicted by node colors. The size of the node circle represents the probability of the ancestral origin for the node of either pediatric or adult age groups. Inner bars adjacent to the tree represent Nextstrain clade designations for each sequence within the annotated clusters. Outer bars indicate nirsevimab breakthrough status. **b,** RSV-A and RSV-B cluster reproductive number (R_0_) respective to time. Blue line indicates baseline R_0_=1, and shadings within each plot indicate the 95% highest posterior density (HPD). **c,** Heatmap of sequences geographical sampling in Chicago and adjacent Chicago suburbs. The nine main geographical regions are referred to as Healthy Chicago Zones as designated by the Chicago Department of Public Health. **d,** Paired bar chart of detected transmission events with 95% HPD intervals between and within age groups from Bayesian phylogeny with in-house and publicly available US genomes with age status for RSV-A (n=466) and RSV-B (n=454).

To determine if these transmission clusters were geographically related, zip code data for the home address of each pediatric patient encounter was mapped to one of nine regional geographic areas, known as Healthy Chicago Zones, as defined by the Chicago Department of Public Health.^24^ We observed statistically significant geographical spread for RSV-A (χ^2^ = 56.71, p-value = 6E-04), but not RSV-B (χ^2^ = 14.76, p-value = 0.065), with biased distributions for specific clusters throughout the city. For example, cluster 1A was predominantly represented by pediatric patients residing in the Northwest, Southwest, and West sides of Chicago neighborhoods (**Figure 2C**).

These pediatric-specific transmission clusters suggest a high transmission rate among kids, though it is unclear how that might compare to transmission rates between kids and adults. To first improve the representation of adult sequences across the 2023-2025 RSV seasons within our transmission analysis, we amended our NM and LCH dataset with publicly available RSV genomes from adults sampled by US surveillance networks.^14, 25^ We then calculated transmission events between and within age groups (**Supplementary Figure 3**). We inferred a high mean number of transmission events occurring either from adult-to-adult or pediatric-to-pediatric patients. Pediatric-to-adult and adult-to-pediatric transmission events, however, were over 10-fold lower for both RSV-A and RSV-B (**Figure 2D**). For RSV-A, we observed that transmission rates from adults to children were slightly higher than from children to adults, indicated by more pediatric sequences stemming from ancestral adult nodes and 95% Highest Posterior Density (HPD) estimates. On the contrary, we observed higher child-to-adult transmission in RSV-B. Regardless, predicted transmission events between age groups (child-to-adult and adult-to-child) were much more infrequent as compared to predicted transmission events within each age group (adult-to-adult and child-to-child).

### Prophylactic breakthrough infections are not linked to Fusion protein resistance mutations

Nirsevimab and the maternal vaccine, Abrysvo, have both been shown to protect infants from severe RSV disease outcomes within 5 months of administration or birth, respectively.^26^ However, they do not protect wholly against infection, especially as antibody levels wane in the months after administration. While infants at high risk of severe disease are eligible to get nirsevimab prior to their experiencing their first and second RSV season, most infants entering their second season have no prophylactic options. It remains unclear how protection of infants in their first respiratory virus season may impact the severity of disease in their second.

To better assess the outcomes and viral determinants associated with prophylactic breakthrough infections, we reconciled our medical record data from pediatric patients presenting at LCH with the Illinois Comprehensive Automated Immunization Registry (I-CARE) and primary care records to identify individuals who received nirsevimab or who were born to mothers with Abrysvo immunization. Of our cohort of 340 pediatric patients with viral sequencing data, 156 (45.9%) were ineligible for either prophylactic, 42 (12.4%) were known to have received nirsevimab, 112 (32.9%) were known to have been eligible but did not received nirsevimab, 5 (1.5%) were born to mothers who received Abrysvo, and 23 (6.8%) were unknown (**Supplementary Table 1**). Regionally, nirsevimab coverage in Chicago was approximated to be 11% and 41% in the 2023-2024 and 2024-2025 seasons, respectively.^27^ RSV detection was nearly 2.5 times higher among eligible infants who did not receive nirsevimab than among those who did, showing a strong protective effect. Looking at the time between nirsevimab administration and the RSV encounter, only 7 infants had a true breakthrough infection within 5 months after administration^28^, while 35 occurred in subsequent seasons after nirsevimab protection waned (**Figure 3A**).

**Figure 3.**
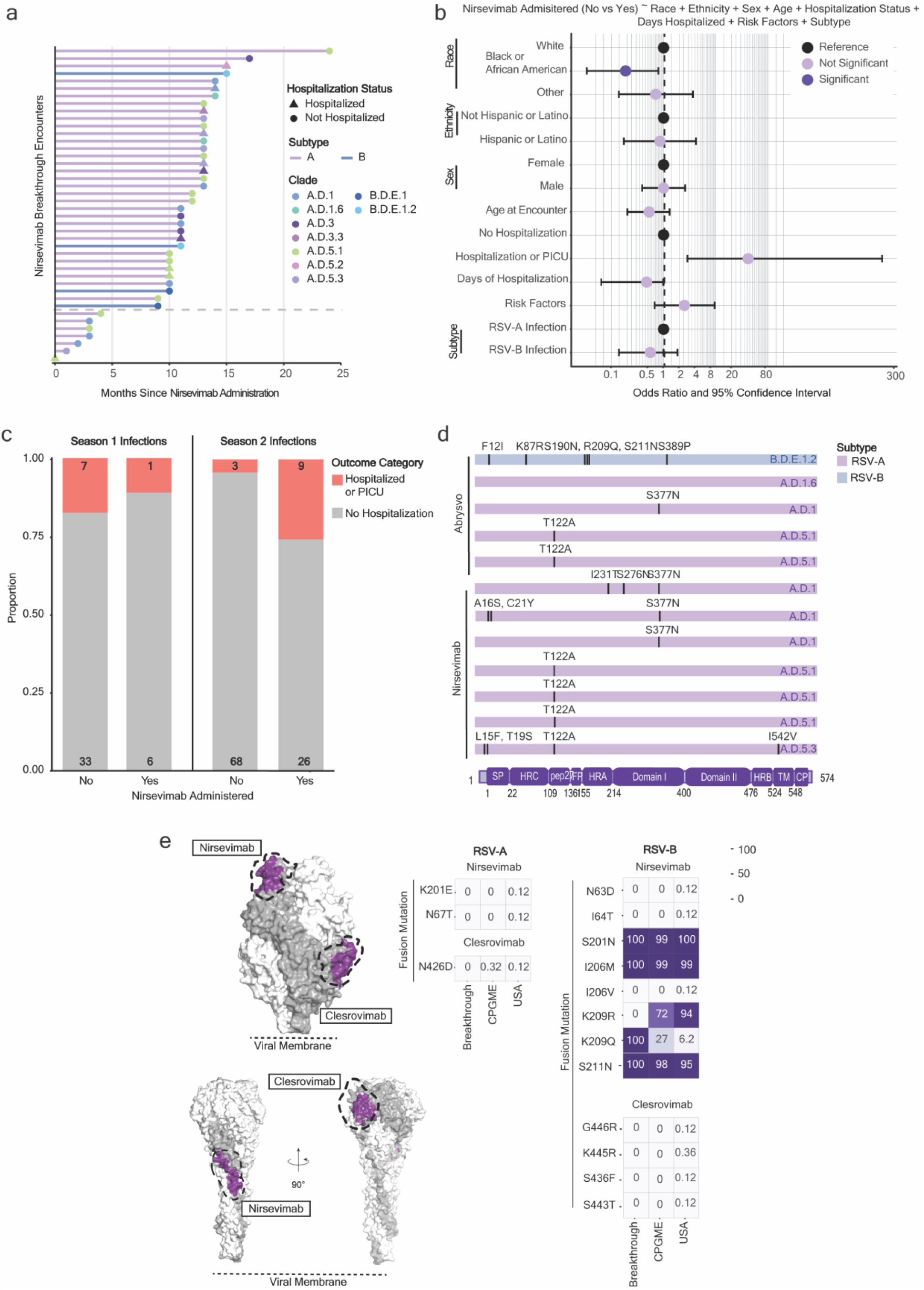
| Clinical and virological assessment of RSV prophylaxis breakthrough Infections. a, Timeframe of patient encounters (n=41) relative to the date of nirsevimab administration **b**, Odds ratio plot with 95% confidence intervals (CI) as calculated by a multivariable logistic regression model with no nirsevimab administered (reference) or nirsevimab administered as the outcome variable for pediatric encounters who experienced an RSV infection during their second RSV season. Significant features (p < 0.05) are highlighted in dark purple (Black or African American; 0.033), insignificant features (p >= 0.05) are displayed in light purple, and reference categories for categorical variables are shown in black (refer to p values in **Supplementary** Figure 4B). **c** Percentage and absolute count of pediatric cohort with an RSV infection in their second season with either hospitalization or PICU admission (orange) or non-hospitalization outcome (grey). **d,** Mutational mapping of sequences obtained from nirsevimab (n=5) or Abrysvo (n-6) breakthrough infections using for RSV-A (Reference NCBI: PP109421) and RSV-B (Reference NCBI: OP975389). **e,** Heatmap displaying mutational frequencies at positions within the nirsevimab, and clesrovimab binding sites within our breakthrough encounters, in-house sequencing, and USA-sampled RSV genomes for RSV-A and RSV-B.

While most breakthrough infections did not require hospitalization (63.6%), the rate of severe outcomes (36.4%) was higher than observed in the overall cohort (15.1%) (**Supplementary Figure 4a**). 3 (60%) out of 5 documented Absyrvo breakthrough infections and 1 (15%) out of 7 documented nirsevimab breakthrough infection required hospitalization or PICU admission. For infants that received nirsevimab and reported an RSV infection in the second season, 9 (25%) required hospitalization. To determine whether there is an association between nirsevimab administration and patient outcome during their second RSV season, we subsampled our cohort to include only infants who were eligible for nirsevimab in the 2023-2024 season (*i.e.*, born between March 1, 2023 and March 31, 2024) and who presented with RSV infection in the 2024-2025 season. We then used a multivariable logistic regression to model nirsevimab administration (*i.e.*, whether a patient received nirsevimab or not in the 2023-2024 season) while controlling for sex, race, ethnicity, age at encounter, hospitalization status, duration of hospitalization, sum of risk factors associated with RSV-infection (prematurity, pulmonary, cardiovascular, immunocompromised, or other chronic conditions), and RSV subtype (**Supplementary Figure 4b**).

The only significant demographic feature was Black or African American race (p-value = 0.033, odds ratio = 0.19), which was significantly associated with a lower likelihood of having been administered Nirsevimab (**Figure 3B**, **Supplementary Figure 4b**). Although it did not reach statistical significance, hospitalization or PICU admission in the second season (i.e. 2024-2025 RSV season) trended towards an association with prior nirsevimab administration (p-value = 0.077, odds ratio 41.7), which was consistent with a notably increased proportion of hospitalization in the nirsevimab group (4.2% hospitalization rate with nirsevimab versus 26% without) (**Figure 3C**). The observed hospitalization rates in our subsampled cohort are suggestive of shift in clinical burden among individuals offered protection via nirsevimab from the first season into the second.

In rare cases, breakthrough infections have been linked to known resistance mutations in the antigenic regions of the viral Fusion protein, but these mutations have generally come at a fitness cost and are not widespread in the viral population.^10^ To determine if any of the nirsevimab or Abrysvo breakthrough infections were associated with any known or novel resistance mutations in the Fusion protein, we mapped the consensus mutations in each sequence relative to either the RSV-A A.D clade reference (PP109421, n = 11) or the RSV-B B.D clade reference (OP975389, n = 1) (**Figure 3D**). While several mutations in RSV-A Fusion were identified, most were associated with the parental clade, and none were in the nirsevimab antigenic site Ø. There were clade specific biases, however, with nearly all breakthrough infections associated with clade A.D.5.1 (3/7 nirsevimab; 2/5 Abrsyvo) or clade A.D.1 (3/7 nirsevimab; 1/5 Abrysvo). Notably, all 7 nirsevimab breakthrough infections were associated with one of the pediatric transmission clusters (1A, 2A, or 3A, **Figure 2A**). In contrast, the sole RSV-B breakthrough to Abrysvo maternal vaccination did have some variation in antigenic site Ø (R209Q and S211N), though both are associated with clade B.D.E.1.2.

To better understand the prevalence of mAb resistance mutations in our cohort and in recent U.S. sequences (2023-2025), we extracted mutational frequency data for all residues in the nirsevimab or clesrovimab antigenic sites (**Figure 3E**, left). There was little to no variation reported in the nirsevimab antigenic site Ø among RSV-A sequences (**Figure 3E**, right). In RSV-B, 3 mutations are approaching fixation (S201N, I206M, S211N) while there is variation at position 209. These mutations, both combinatorially or singularly, have previously shown either no or modest impact in neutralization potency of nirsevimab.^20^ There was little to no variation in the clesrovimab binding site for RSV-A, except for a rare N426D (0.12%) mutation, which has been shown to confer resistance to a monoclonal antibody precursor that also targets antigenic site IV ^29^ (**Figure 3E**). Similarly, we observe only rare mutations in the clesrovimab binding site among RSV-B sequences, namely G446R (0.12%), K445R (0.36%), S436F (0.12%), and S443T (0.12%). Although there is no documented evaluation of these specific amino acid substitutions, mutations at sites 443, 445, 446 in Fusion are known to occur *in vitro* during serial passaging and confer reduced susceptibility to clesrovimab.^30^

### RSV-A F proteins from contemporary clades confer similar fusogenicity and cell entry activity

Nirsevimab breakthrough isolates were associated with a few specific clades found in pediatric transmission clusters but otherwise shared no mutations in the Fusion (F) protein that would suggest a common escape mutation. Relative to commonly used reference strains, contemporary circulating clades have several F protein mutations that may impact viral entry and/or neutralization. The top 5 circulating RSV-A clades have 16 shared mutations relative to the reference strain A2 with additional clade defining mutations in the signal peptide (SP), heptad-repeat C (HRC), and 27-mer fragment (pep27) domains (**Figure 4A**, left). The top 5 circulating RSV-B clades have 8 shared mutations relative to the reference strain B1 with additional clade defining mutations in the SP, heptad-repeat A (HRA), and Domian I regions (**Figure 4A**, right).

**Figure 4.**
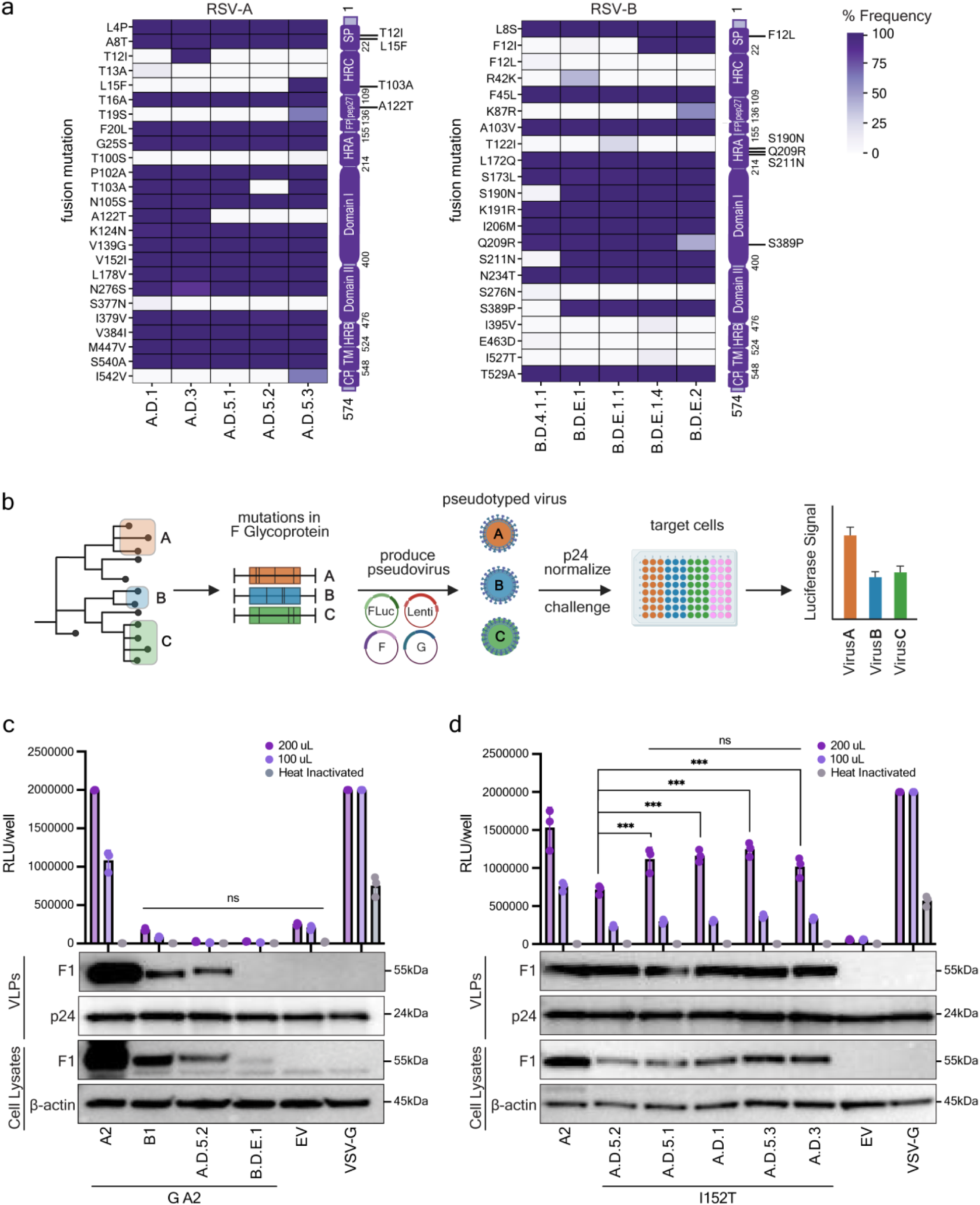
| Identification and characterization of viral entry in contemporary RSV clades. **a**, Heatmap displaying mutational frequencies of RSV-A (left) and RSV-B (right) top 5 circulating strains in reference to NCBI Accession WWC50954.1for A and NP_056863.1for B, respectively. A schematic of F domains with clade-defining mutations shown on the right. **b,** Schema of the RSV pseudotyping assay. Lentiviral pseudoviruses bearing RSV F and RSV G from the indicated viral strains were produced in HEK293T cells and were quantified and normalized via p24 ELISA before infection of TIM-1–expressing HEK293T target cells. Viral entry was quantified by luciferase activity at 72 hours post-infection **c,** Lentiviral pseudoviruses bearing full-length [A2, B1, A.D.5.2, B.D.E.1] F and G glycoprotein from RSV A2 with 31 AA deletion were generated to assess a measurable readout to determine viral entry efficiency. **d,** Lentiviral pseudoviruses bearing truncated [A2, A.D.5.2, A.D.5.1, A.D.1, A.D.5.3, A.D.3] F with each contemporary clade bearing mutant (I152T) were generated to measure viral entry differences between clades. **c-d,** Viral entry was quantified by luciferase activity at 72 hours post-infection. Cell lysates (CL) and purified virus-like particles (VLPs) normalized to p24 were analyzed by immunoblotting for each RSV F. p24 was used as a VLP loading control, and β-actin as a cellular loading control. Data points represent the mean ± s.d. from three independent biological replicates. Statistics were calculated using two-way ANOVA followed by Tukey’s multiple comparisons test. Pointed brackets indicate statistical comparisons between two specific conditions. Flat horizontal lines denote comparisons across all conditions. ns, not significant; p < 0.05 (\**); p < 0.01 (*\*\*); p < 0.001 (***); p < 0.0001 (****).

To assess the phenotypic impact of these mutations on cell entry and mAb neutralization, we leveraged a previously described lentiviral pseudotyping system.^31^ Briefly, luciferase reporter viral-like particles (VLPs) were generated in HEK293T cells with incorporation of G glycoprotein and different clade-specific F glycoproteins (**Figure 4B**). Virus stocks were normalized to lentiviral p24 prior to challenge of TIM1-expression HEK293T cells and luciferase readout at 72 hours post-challenge. All challenges were performed in technical triplicate at two doses to show titratability alongside heat inactivated and no F protein (empty vector) controls. VSV-G–pseudotyped VLPs were included as positive controls. Human codon optimized F overexpression constructs were generated for the top 5 circulating lineages of RSV-A and RSV-B (**Figure 4A**) by the addition of the relevant clade-defining mutations to a base construct containing the most recent common ancestor (**Supplementary Table 4**).

As proof-of-concept, the F proteins from lab-adapted RSV-A A2 and RSV-B B1 were compared with those from two representative contemporary clades, A.D.5.2 and B.D.E.1, respectively. Each F protein was paired with an N-terminally truncated G glycoprotein^31^ from lab-adapted RSV-A2 to ensure that any differences in viral entry were due solely to changes in F. As expected, VSV-G conferred efficient viral entry with signal at the assay maximum at both challenge volumes (**Figure 4C**). RSV-A A2 F likewise conferred titratable and efficient viral entry. RSV-B B1 and both contemporary strains, however, did not yield a signal that was statistically different from the empty vector. This largely correlated with F expression in producer cell lysates (CLs) and incorporation into VLPs with RSV-A A2 showing much higher levels of F relative to RSV-B B1 and both contemporary isolates (**Figure 4C**).

In an attempt to rescue entry efficiency, we first truncated four amino acids from the cytoplasmic tail of F, as prior studies have shown this region modulates fusion activity and viral assembly.^32,33,34^ The full-length (FL) or truncated (Tr) versions of each F glycoprotein were then paired with either the lab-adapted RSV-A A2 or corresponding contemporary G glycoprotein, each harboring a 31 amino acid deletion. The FL and Tr versions of RSV-A A2 F protein showed comparable expression, incorporation, and cell entry (**Supplementary Figure 5A**). The Tr version of the RSV-A A.D.5.2 F protein showed slightly improved expression and incorporation compared to the FL protein, but it was insufficient to facilitate detectable viral entry, regardless of the co - expressed G glycoprotein (**Supplementary Figure 5A**). For RSV-B, truncation of the RSV-B B1 F protein was sufficient to dramatically increase F expression, incorporation, and cell entry (**Supplementary Figure 5B**). However, while again the Tr version of the RSV-B B.D.E.1 F protein showed slightly improved incorporation compared to the FL protein, it was still insufficient to facilitate detectable viral entry, regardless of the co-expressed G glycoprotein (**Supplementary Figure 5B**). This was not specific to these contemporary clades, as truncated versions of the other circulating RSV-A (A.D.5.1, A.D.1, A.D.5.3, and A.D.3) and RSV-B (B.D.E.1.1, B.D.4.1.1, B.D.E.2, and B.D.E.1.4) F proteins likewise failed to yield detectable viral entry activity in this assay (**Supplementary Figure 5C-D**).

We next sought to test if we could rescue the expression and activity of the RSV-A A.D.5.2 F protein by reversion of one of the 13 amino acid differences it has with RSV-A A2 F. Each substitution was introduced individually into the truncated RSV-A A.D.5.2 F protein construct and assessed for entry activity as before (**Supplementary Figure 5E**). Among all mutations tested, only one mutation in the fusion peptide (I152T) was found to enhance protein expression, incorporation, and viral entry activity. While a second mutation (N515H) also significantly increased viral entry activity relative to the wild-type A.D.5.2 construct, it was not to the same extent as I152T and did not notably alter F expression or incorporation (**Supplementary Figure 5E**). Note that we also sought to rescue the expression and activity of the RSV-B B.D.E.1 F protein by reversion of one of the 13 amino acid differences it has with RSV-B B1 F, but none of the tested mutations restored activity above background (**Supplementary Figure 5F**).

Finally, we engineered the I152T rescue mutation into the truncated F expression constructs for the top circulating RSV-A clades (A.D.5.2, A.D.5.1, A.D.1, A.D.5.3, and A.D.3) and compared their entry activity head-to-head (**Figure 4D**). All five F constructs yielded similar F expression and incorporation, with slightly lower entry efficiency compared to the RSV-A A2 control. Between clades, A.D.5.2 had slightly lower activity at higher dose challenges, though these differences were not apparent at lower doses (**Figure 4D**).

To confirm these results, we used these same constructs for an orthogonal split-reporter cell-cell fusion assay that evaluates fusion independently of lentiviral particle incorporation. Briefly, HEK393T cells expressing each RSV-A fusion protein and T7 polymerase were co-cultured with HEK293T cells containing a T7-driven luciferase reporter construct with cell-cell fusion monitored by luciferase activity 48 hours post co-culture^35^ (**Supplementary Figure 6A**) Co-transfection of T7 polymerase and the luciferase reporter resulted in maximum luciferase signal while the empty vector control gave minimal background. While all lower than the RSV-A A2 F protein, there were no significant differences in fusion activity between the 5 contemporary RSV-A clades (**Supplementary Figure 6B**). The addition of the I152T rescue mutation enhanced steady-state levels of F protein and increased the fusion activity of each construct, but again there were no significant differences in fusion activity between the 5 contemporary RSV-A clades (**Supplementary Figure 6B**). The RSV-B B1 F protein did show fusion activity in this assay, but again all RSV-B F proteins from contemporary clades failed to show signal above the empty vector control (**Supplementary Figure 6C**). Overall, this suggests that the F proteins from all 5 contemporary RSV-A clades have similar entry efficiencies.

### A.D.3 Fusion is slightly less susceptible to nirsevimab neutralization, though no clade is strongly resistant

A previous study reported comparable protective effects of nirsevimab across all circulating RSV clades during the 2023–2024 respiratory season.^10^ To extend these findings for the 2024–2025 season, we adapted our pseudotyping assay to quantify the neutralization potency of FDA-approved mAb therapeutics (palivizumab, clesrovimab, and nirsevimab) across the 5 most prevalent circulating RSV-A clades. Over an 8-point dose curve, palivizumab and clesrovimab both demonstrated comparable neutralization potency across all tested clades with no significant differences in neutralization IC50 values **(Figure 5A-B).** As previously reported, mutations N262D and G446E conferred resistance to neutralization by palivizumab and clesrovimab, respectively (PMID: 22184728, PMID: 31515478) **(Figure 5A-B)**.

**Figure 5.**
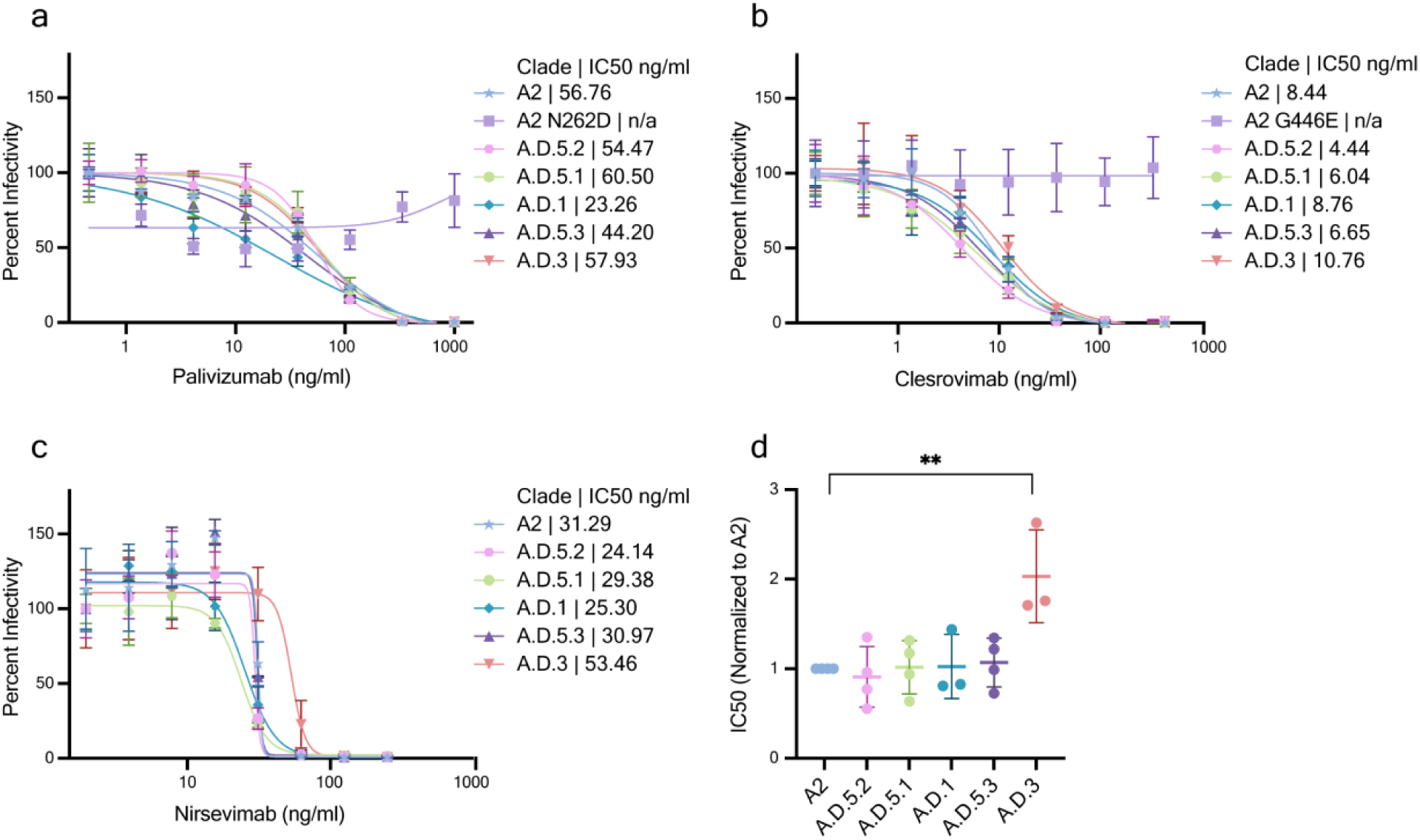
**| Evaluation of mAbs neutralization efficacy across RSV-A contemporary clades. a-c**, Dose–response neutralization curves for RSV-A pseudoviruses bearing the truncated F protein from contemporary clades [A2, A.D.5.2, A.D.5.1, A.D.1, A.D.5.3, A.D.3] and drug-resistant controls F [N262D and A2 G446E] treated with serial dilutions of **a,** palivizumab, **b,** clesrovimab or **c,** nirsevimab. Dose response curves were quantified by luciferase activity at 72 hours post-infection and plotted as percent infectivity relative to untreated controls. Data represent mean ± s.d. of technical triplicates within a single experiment. **d,** Nirsevimab neutralization IC_50_ across contemporary RSV-A clades. Half-maximal inhibitory concentration (IC₅₀) values for each virus–drug combination were calculated by nonlinear regression using a four-parameter logistic model. **\***A data point was excluded due to technical error in A.D.3 during data acquisition. Data is plotted as IC_50_ relative to RSV-A A2 neutralization IC_50_ across four independent replicates (mean +/- s.d.). Statistics were calculated using one-way ANOVA followed by Tukey’s multiple comparisons test. Pointed brackets indicate statistical comparisons between two specific conditions. ns, not significant; p < 0.05 (*); p < 0.01 (**); p < 0.001 (***); p < 0.0001 (****).

Over a similar 8-point dose curve, nirsevimab likewise demonstrated comparable neutralization potency across most of the tested clades, though A.D.3 had a notably higher neutralization IC50 value **(Figure 5C)**. Four independent replicates of this experiment confirmed that A.D.3 displayed a roughly 2-fold increase in neutralization IC50 value compared to lab-adapted RSV-A A2 and other contemporary RSV-A clades **(Figure 5D)**. RSV-A A.D.3 F protein has only a single unique substitution mutation (T12I) within the signal peptide domain. Although other clades contain mutations within this domain, this specific substitution is exclusive to A.D.3 and is the only mutational difference between the A.D.1 and A.D.3 F proteins. Notably, however, none of the first season breakthrough infections in this study were with RSV-A A.D.3, suggesting that this change in IC50 is not sufficient to cause loss of clinical efficacy. Collectively, these data suggest that RSV prophylactic breakthrough is driven by high exposure in the context of pediatric transmission clusters rather than emergent resistance in the Fusion protein of a given clade.

## DISCUSSION

In this dual-center, retrospective cohort study, we performed whole-genome sequencing of RSV isolates from infants and adults during the past two RSV seasons (2023-2025) to assess the epidemiological and phenotypic changes in contemporary RSV viral populations in the U.S. Leveraging our genomic analyses, we also aimed to identify viral and clinical correlates of nirsevimab breakthrough infection. We noted the presence of several temporally and geographically constrained pediatric transmission clusters within our cohort, which were associated with high levels of child-to-child transmission. While breakthrough isolates did not share any F glycoprotein mutations, they all occurred in the context of one of these transmission clusters. The F proteins from the clades associated with these clusters did not differ in relative fusogenicity or nirsevimab neutralization susceptibility. Fortunately, these breakthrough infections were infrequent and resulted in only mild disease, though infections in the second season after administration trended more severe. Collectively, these data suggest that clade associations with prophylactic breakthrough are driven by pediatric transmission clusters rather than clade-associated resistance and suggest the need for continued monitoring in kids in their second season after nirsevimab administration due to a potential shift in clinical burden.

We demonstrate that, as RSV restored pre-COVID-19 pandemic seasonality patterns, RSV-A continued to diversify iteratively with multiple clades co-circulating within a single season. In contrast, RSV-B B.D.E.1 clade and genetically similar subclades have persisted in recent seasons. Upon evaluating regional, pediatric-specific transmission clusters, we observed differences in transmission dynamics across several RSV-A and -B contemporary clades, along with the detection of nirsevimab breakthrough cases within these clusters. Clinical evaluations of our patient encounters at NM and LCH indicated overall mild manifestations with evidence of a clinical burden shift in pediatric cohorts with administered nirsevimab experiencing an RSV infection the following season. When evaluating several F mutations identified outside of mAb epitopes using an optimized, novel pseudotyping platform, we found that the majority of currently circulating RSV-A clades exhibited similar cell entry capabilities and were potently neutralized by palivizumab, clesrovimab, and nirsevimab.

Our phylogenetic analyses demonstrated that as RSV restored pre-COVID-19 pandemic seasonality patterns, RSV-A continued to diverge with multiple co-circulating clades while RSV-B has been dominated by a more limited set of genetically similar clades. These data recapitulate broader RSV genomic trends in the U.S. Several multicenter studies reported a similar predominance of A.D.3, A.D.5, and B.D.E.1 subclades in the US as of late 2023, with some studies also reporting regional-specific clades, such as A.D.1.6 in Maryland.^36,25^ Nonetheless, clade distribution patterns suggest RSV-A evolutionary dynamics, both before and after the 2022 resurgence, is markedly distinct from that of RSV-B, with multiple RSV-A subclades co-circulating in any given season compared to a more homogenous RSV-B population structure. These data also make clear the effect of RSV population bottlenecking event during the COVID-19 pandemic with dramatic shifts in population structures before and after the implementation of NPIs to prevent the spread of SARS-CoV-2 .^16^

Since infant and adult patients were included in this study cohort, we were uniquely positioned to address genomic gaps in population transmission patterns across age groups. Adult-to-adult and child-to-child transmission rates were inferred to be higher than transmission rates between age groups. We also found the presence of pediatric-specific transmission clusters within our cohort, most of which were both geographically and temporally related. The high rates of child-to-child transmission align with several previous studies on family transmission dynamics and in childcare settings. ^37,38,39^ Higher viral loads in children under 2 years of age may drive the rapid expansion of these populations and may facilitate nirsevimab breakthrough infections. The rate of transmission between adults was likewise higher than observed transmission rates between age groups. Regardless, there was little evidence in this cohort of transmission clusters among adults. This could be due to lower viral loads in this population, a decreased frequency of gathering of susceptible adults, and/or a bias in sampling. More studies that conduct sequencing of RSV isolates from both adult and pediatric population will be needed to extend these results.

As of mid 2025, cumulative RSV-associated hospitalizations have decreased among infants aged 0-7 months, but has slightly increased in older children as compared to pre-COVID-19 rates. ^40,41^ , Although one study has documented no clinical outcome difference in hospitalized cohorts with or without nirsevimab^42^, the shift in hospitalization rates influenced by RSV prophylaxis has not been fully interrogated. Our clinical regression analysis likewise suggests a modest shift in increased hospitalization in infants who had nirsevimab administered in their first respiratory season and encountered an RSV infection in their second season. As the susceptible age for severe clinical outcome from RSV infection appears to shift onto older infants due to new prophylaxis, the consequences of shift in clinical burden should be closely monitored. Similarly, our regression analysis findings of racial disparities in nirsevimab administration can inform future efforts to provide equitable protection and prophylaxis rollout across RSV-susceptible populations. Because adult populations have been understudied compared to pediatric counterparts, merging datasets to analyze transmission patterns can help inform more intentional public health efforts to allocate resources and design initiatives to mitigate the RSV burden. For example, as infants aged 0–6 months become immunized, reductions in RSV-associated hospitalizations and onward transmission can indirectly protect older age groups^43^, underscoring the importance of understanding the broader RSV ecosystem.

Our efforts to integrate epidemiological and molecular approaches stemmed from the ongoing disruption of RSV’s virological niche, driven by turbulent transmission dynamics and selective pressures from RSV preventatives. The optimization of molecular tools in this study, probing for the phenotypic consequences of RSV diversity, will contribute to future efforts to monitor shifts in both viral behavior and mAb escape. Several mutations, albeit rare in the global population^44^, have already been well documented to confer resistance to nirsevimab. Although clesrovimab targets F’s highly conserved antigenic site IV, which reduces concerns for emerging escape mutants^45^, its ongoing rollout contributes to the ongoing external selective pressures, which were previously nonexistent in RSV. These population-level measures, compounded by the continuous antigenic pressure from the immune response to detect and suppress RSV during infection contribute to the viral adaptations with unknown phenotypic consequences. Tangentially, concerted efforts that have predominantly focused on the clinical aspects of F fusion have created large gaps in fundamental molecular knowledge regarding the role of genetic diversity in F fusogenicity and pathogenicity. We identified that RSV clade A.D.3 containing the T12I mutation shows a slight, but not drastic, reduction in neutralization by Nirsevimab. Although this mutant is found in the signal peptide of the F ORF, mutations localized to other domains in F (such as L305I) have demonstrated a probable role in altering neutralization epitopes.^46^ This is aligned with studies in other viruses where changes in the signal peptide can drive changes in antibody neutralization due to altered intracellular trafficking and glycosylation patterns.^47^ As such, F protein mutations outside of the antigenic sites should be closely monitored and studied. Ongoing efforts to develop tools to assess fundamental traits such as F stability^48^, fusogenicity^46^, and expression^49^ can also strengthen future, timely responses to emerging F diversity.

Several limitations in the study design and in publicly available data should be taken into consideration when interpreting our results. The absence of retrospective diagnostic LCH specimens and paired clinical data before the 2023 RSV season hindered our ability to study long-term shifts in RSV behavior. Similarly, our incidental catchment of adult populations decreased significantly as universal respiratory testing practices within hospital systems shifted, both within and outside our hospital sites.^50^ As previously described, our molecular work was largely confined to RSV-A, as our construct modifications could not restore fusogenic activity in RSV-B constructs outside lab-adapted strains, a common obstacle in RSV molecular work.^51^

Overall, this integrative approach characterizes contemporary RSV viral patterns that warrant close monitoring as RSV preventive measures roll out globally. Our goals were to (1) define genomic trends after restored seasonality, (2) characterize clinical and transmission dynamics across age groups, and (3) assess viral entry and antibody evasion capabilities of the most prevalent circulating RSV clades. Our findings indicate that breakthrough infections are primarily correlated with predominant contemporary clades rather than fusion resistance, supporting the continued clinical effectiveness of mAb prophylaxis. By integrating genomic, phylogenetic, and functional data, our approach enables the identification of emerging lineages and directly informs prophylactic use, patient risk stratification, and outbreak control.

## METHODS

### Ethics approval

Collection of RSV-positive residual diagnostic nasopharyngeal (NP) specimens for viral whole genome sequencing was approved by Northwestern University Institutional Review Board through #STU00212260 and Lurie Children’s Hospital IRB through 2025-7302. Access to clinical and demographic data of RSV-positive patient encounters for Northwestern Medicine and LCH was granted through IRBs #STU00206850 and 2025-7507, respectively. The corresponding metadata reported in the study are stored on a Research Electronic Data Capture (REDCap) server in compliance with HIPAA regulations. Data were de-identified and linked to sequencing data for analysis using non-descriptive, unique identifiers.

### Public Data Extraction

Publicly available whole-genome sequences were extracted from the NIH NCBI Nucleotide Database. Accession number and metadata information are available at: (http://github.com/erg6437/RSV-Evolution).

*<u>Northwestern Medicine and Lurie Children’s Hospital Data Extraction and Specimen Collection</u>* Demographic and clinical metadata from patients encounters with RSV infections were extracted from NM and LCH’s enterprise data warehouse and manually reviewed from the electronic health record. Biobanks of RSV diagnostic specimens were formed under the Center for Pathogen Genomics and Microbial Evolution (CPGME) at Northwestern University. The residual diagnostic nasopharyngeal swabs from patients with a confirmed positive RSV infection in the Northwestern Medicine healthcare system were collected from December 2017, through March 2026. Similarly, the residual specimen collected by Lurie Children’s Hospital was collected from November 2024 to March 2025.

### Viral RNA Extraction and Quantification

NP specimens stored in viral transport media were utilized for viral RNA extraction using the QIAamp Viral RNA Minikit (Qiagen, cat. no. 52906) and the QIAcube HT Kit (Qiagen, cat. no. 51331). In addition, NP specimens were utilized for total nucleic acid extraction using the MagMAX™ Viral/Pathogen Nucleic Acid Isolation Kit (Applied Biosystems, cat. No. A48310). Viral load quantification and RSV subtype identification were obtained by performing quantitative reverse transcription and PCR (qRT-PCR) using primer and probe set from the World Health Organization Strategy for Global RSV Surveillance Project based on the Influenza Platform.^52^ Primers for RSV polymerase [L] (Forward: 5’-AATACAGCCAAATCTAACCAACTTTACA-3’ and Reverse: 5’-GCCAAGGAAGCATGCAATAAA-3’) were used to target both RSV-A and RSV-B, whereas probes (RSV-A: 5’-TGCTATTGTGCACTAAAG-3’ & RSV-B: 5’-CACTATTCCTTACTAAAGATGTC-3’) distinguish between subtypes. Specimens with either an RSV-A L or RSV-B L probe cycle threshold (Ct) value less than or equal to 28 were considered of sufficient virtual load for subsequent sample preparation for whole genome sequencing.

### cDNA Synthesis and Viral Genome Amplification

cDNA synthesis was performed using SuperScript IV First-Strand Synthesis Kit (Invitrogen, cat. no. 18091050) using RSV-specific primers.^22^ Direct amplification of RSV genome cDNA was performed in separate PCR reactions to generate ∼4000 base pair fragments that span the genome. PCR amplification of the generated cDNA was performed using the Phusion Hot Start Flex system (New England Biolabs, cat. no. M0536L). Genomic amplification was confirmed via agarose gel electrophoresis, followed by reaction pooling and reaction cleanup using MAGwise Paramagnetic Beads (seqWell, cat. no. ABIN7271581). After verification of genome amplification, the four amplicon sets were pooled before sequencing library preparation.

### Sequencing Library Preparation and Whole Genome Sequencing

Two separate approaches to library preparation for sequencing on the Illumina MiSeq platform were used in this study: seqWell’s plexWell and expressPlex. For the seqwell protocol, up to 384 ng of DNA was used for the total Viral genome sequencing reads were utilized to generate consensus sequences using the HAplotype and PHylodynamics pipeline (HAPHPIPE) for viral assembly, population genetics, and phylodynamics.^53^ This pipeline consists of the following: (1) trimming sequencing reads to remove both adapters and low-quality sequences using Trimmomatic v1.0 and (2) three-step assembly refinement. The generated consensus reads were then annotated using Viral Genome ORF Reader (VIGOR) v. 4.0.^54^

### Phylogenetic analyses

Generated consensus sequences and ORFs were aligned using MAFFT v7.490, followed by inspection and removal of poorly aligned positions using the Gblocks^55^ feature in Seaview v. 5.0.4.^56^ IQ-Tree’s v. 2.2.0 ModelFinder function^57^ was used for the selection of the nucleotide substitution model best fitted for each alignment and subsequent inference of Maximum Likelihood (ML) phylogeny. Assessment of tree topology was done ultrafast bootstrap (UFboot) and Shimodaira–Hasegawa approximate likelihood-ratio test (SH-aLRT) each with 1000 replications. TN93 genetic distances to determine transmission clusters was performed by using HIV-TRACE and setting a maximum pairwise distance of less than 0.0011 and 0.00299 substitutions per site for RSV-A and RSV-B, respectively.^58^ ORF analysis was done by extracting cds information using VIGOR v4.1.20200702. which was then used for subsequent phylogenetic and selection analysis.

### RSV Temporal Dynamics

Samples from NM/LCH were combined with contemporaneous GISAID for which sample date and participant age metadata were available. Viral transmission dynamics were estimated under a Bayesian serial birth-death skyline framework in BEAST 2.7, using the BDSKY v1.5.1 package and an optimized relaxed molecular clock model. ^59,60,61, 62,63,64,65,66^ The age of the sample donor, categorized as pediatric or adult, was included as a discrete trait. Model convergence was evaluated by confirming effective sample size (ESS) values ≥200 for all estimated parameters.^67^ Expected numbers of transitions within and between age groups were extracted using BEAST’s Babel package. All data processing and visualization were performed in R v4.4.3 using the ggtree package.

### Cells

Human embryonic kidney (HEK)293T, and the HEK293T cells expressing TIM-1 (provided by Dr. Cassandra Simonich at the Fred Hutchinson Cancer Center) were maintained in a humidified cell culture incubator at 37 °C with 5% CO₂. Cells were cultured in 1x Dulbecco’s Modified Eagle Medium (DMEM) supplemented with 10% heat-inactivated fetal bovine serum and 1% penicillin/streptomycin.

### Pseudovirus Stock Preparation

HEK293T cells are seeded at a density of 1.5 × 10⁶ cells in 10 cm tissue culture plates and incubated for 24 hours at 37°C with 5% CO₂ before transfection. The transfection mixture contains 11.69 μg of plasmid DNA. The luciferase reporter construct is included at 5.8 μg, followed by three lentiviral packaging plasmids encoding gag-pol, tat, and rev, each at 1.26 μg. RSV surface protein expression plasmids are added at 1.17 μg for RSV-F and 0.58 μg for RSV-G, with the specific F and G expression plasmids differing based on the subtype being tested in the assay. For control conditions, RSV surface protein plasmids are replaced with either VSV-G or an empty vector. The plasmid mixture is combined with 300 μL of serum-free DMEM and 30 μL of PolyJet transfection reagent, according to the manufacturer’s instructions, and then applied to HEK293T cells.

After 48 hours of incubation, the supernatant containing pseudovirus particles is harvested and filtered through a 0.45 μm pore-size filter to remove cellular debris. Viral particle concentration is quantified using a p24 ELISA (Cat# ab218268), and the resulting values are used to normalize and dilute the pseudovirus stocks in supplemented DMEM.

For the infection assay, 293T-TIM1 cells are seeded at 40,000 cells per well in 200 μL of supplemented DMEM in 96-well white Costar plates (Product #3917) one day before infection. Normalized pseudovirus preparations are then overlaid onto the plated cells in volumes of either 200 μL or 100 μL. Heat-inactivated virus controls are prepared by incubating the pseudovirus at 60°C for 30 minutes before overlaying it onto the challenge cells. Plates are incubated for 72 hours at 37°C with 5% CO₂.

Following incubation, viral entry efficiency is assessed by measuring luciferase activity using the Promega Luciferase Assay System (Ref. No. E2620) on an Omega plate reader. Relative luminescence units (RLU) are recorded, and technical replicates are included within each experiment.

### p24 ELISA for Viral Titer Quantification

After harvesting with a 0.45-um-pore-size filter, aliquots of pseudoviruses were diluted with the kit-provided sample diluent (NS), starting with a 1:20 dilution and continuing with a 1:5 dilution series. Viral stock concentrations were quantified relative to p24 reference standards (300 to 4.69 pg/mL) provided in the kit, using an internal standard curve. P24 values were measured on an Omega plate reader, and the OD was recorded at 450 nm.

### Pseudovirus Neutralization

To prepare for the neutralization assay, the same methods described in the Pseudovirus Stock Preparation section are followed, with one modification: before overlaying the virus onto the challenge cells, the virus and monoclonal antibody (mAb) are incubated together at 37°C with 5% CO₂ for 1 hour. A setup plate is prepared with serial dilutions of the respective mAb, using either a 1:3 or 1:2 dilution scheme to generate an eight-point dose-response curve. After the incubation period, the virus–drug mixture is overlaid onto the challenge cells and incubated for 72 hours at 37°C with 5% CO₂. Viral entry is then quantified using the same luciferase-based readout (described in the Pseudovirus Stock Preparation section).

### Immunoblotting

HEK293T and 293 TIM-1 cells transfected with pseudovirus and fusion-related plasmids were used to prepare protein lysates. Briefly, cells were washed using Dulbecco’s Phosphate-Buffered Saline (DPBS) and resuspended in 300 μL of reducing sample buffer (RSB) (Tris-HCl, 50% glycerol, 10% SDS,-mercaptoethanol, and 1% bromophenol blue). Following sucrose-gradient centrifugation, viral pellets were resuspended in RSB and normalized based on p24 levels, using a final volume of 50 μL corresponding to the lowest p24 sample. Cell and viral lysates were heated to 98 °C for 20 minutes and then immediately frozen at -20 °C for storage. After running lysates on pre-cast 4-20% Criterion Tris-HCl gels at 90 V for 30 minutes, followed by 150 V for 50 minutes, a wet transfer onto a polyvinylidene fluoride (PVDF) membrane was performed at 90 V for 2 hours. Afterwards, the membrane containing the transferred protein was blocked with 5% milk, 0.1% Tween 20 diluted in PBS for 1 hour, followed by primary antibody incubation overnight at 4 °C. After PBS washing before and after incubation with conjugated IgG horseradish peroxidase (HRP) secondary antibodies at room temperature for 1 hour, proteins were visualized using enhanced chemiluminescence (ECL) Western Blotting Substrate on an iBright blot scanner. Blots were subsequently incubated at room temperature for 30 minutes with antibody stripping solution before reprobing.

### Plasmids

Plasmids expressing HSV glycoproteins (gB, gD, gH, and gL) and nectin-1 (pBG380) were generously provided by Qing Fan at Northwestern University. Plasmids required for pseudovirus production were generously provided by Dr. Cassandra Simonich at the Fred Hutchinson Cancer Center. The system uses a third-generation lentiviral packaging system comprising three helper expression plasmids: 26_HDM-Hgpm2 (Gag-Pol), 27_HDM-tat1b (Tat), and 28_pRC-CMV-Rev1b (Rev). A lentiviral backbone plasmid was also provided (catalog NR-52516). This backbone uses a CMV promoter to express luciferase, followed by an IRES and ZsGreen.

Human codon-optimized full-length fusion ORFs A.D.5.2 and B.D.E.1 were designed and inserted into pcDNA4/TO empty vector via Gibson Assembly. Truncated F constructs were amplified using primers and re-inserted into the pcDNA4 vector.

Fusion point and rescue mutations for all contemporary and lab strain RSV-A and RSV-B clades were generated using site-directed mutagenesis (SDM). The PCR reactions were subsequently transformed into chemically competent *E. coli* (Takara Bio, #636763), expanded, and isolated for mutation verification via Sanger sequencing. All Gibson assembly and SDM constructs used in this study were confirmed by Sanger sequencing with the primers listed in **(Supplementary Table 4)**.

### RSV Fusion Assays

HEK293T cells were used as effector and target populations in a split reporter–based cell–cell fusion assay. HEK293T target cells were transfected with a T7 promoter–driven luciferase reporter plasmid. HEK293T effector cells were transiently transfected with plasmids encoding viral envelope glycoproteins together with T7 polymerase to enable reporter expression upon membrane fusion.

For transfection, HEK293T cells were seeded at 200,000 cells per well in 6-well plates. Effector cells were transfected with 0.4 µg T7 polymerase and either 0.55 µg RSV F glycoprotein or 1.1 µg total plasmid encoding HSV envelope glycoproteins (gB, gD, gH, and gL). Target cells were transfected with 0.4 µg luciferase driven by a T7 promoter together with 1.1 µg empty vector. Twenty-four hours post-transfection, effector cells were gently harvested by mechanical perturbation, counted, and mixed with target cells at a 1:1 ratio. Mixed cell populations were seeded into 96-well Costar plates (Product #3917) and co-cultured to allow membrane fusion. Fusion activity was quantified 48 hours after co-culture by measuring luciferase activity using the Promega Luciferase Assay System (Ref. No. E2620) on an Omega plate reader (BMG LabTech).

## ACKNOWLEDGEMENTS

The authors are grateful for the assistance of Dr. Cecilia Thompson (Lurie Children’s Hospital) and Drs. Timothy Blanke, Michael Malczynski, Chao Qi, and Kendall Kling (Northwestern Memorial Hospital) in specimen acquisition. The authors would like to thank Dr. Qing Fan (Northwestern University) for providing the HSV and reporter plasmids necessary for the fusion assay, Dr. Cassandra Simonich (Fred Hutchinson Cancer Center) for providing the TIM-1 cells and plasmids necessary for the pseudotyping assay, Dr. Heba Mostafa and his team (Johns Hopkins School of Medicine), and Dr. Adam Lauring and his team (University of Michigan) for providing sequencing metadata for transmission analysis. This research was supported in part through the computational resources and staff contributions provided by the Genomics Compute Cluster, which is jointly supported by the Feinberg School of Medicine, the Center for Genetic Medicine, and Feinberg’s Department of Biochemistry and Molecular Genetics, the Office of the Provost, the Office for Research, and Northwestern Information Technology. The Genomics Compute Cluster is part of Quest, Northwestern University’s high-performance computing facility, to advance genomics research. We further gratefully acknowledge all data contributors, i.e., the Authors and their Originating laboratories responsible for obtaining the specimens, and their submitting laboratories for generating the genetic sequence and metadata and sharing via the GISAID Initiative, on which this research is based. All schematics were created using BioRender.com.

## DATA AND CODE AVAILABILITY STATEMENT

Code, tools, and parameter settings required to reproduce these results, including our genome assembly pipeline and within-host variant analysis, is available on GitHub at: http://github.com/erg6437/RSV-Evolution. All viral whole genome consensus sequences have been uploaded to NCBI (accession numbers provided in **Supplemental Table 1**). The individualized clinical data reported in this study cannot be deposited in a public repository due to IRB constraints, but population-level metrics are provided in **Table 1**.

## AUTHOR CONTRIBUTIONS

**Estefany Rios-Guzman**: Conceptualization, Methodology, Software, Validation, Formal Analysis, Investigation, Resources, Data Curation, Writing – Original Draft, Writing – Review and Editing, Visualization; **Ria Almohtadi**: Conceptualization, Methodology, Validation, Formal Analysis, Investigation, Resources, Writing – Original Draft, Writing – Review and Editing, Visualization; **Seth H. Borrowman**: Methodology, Software, Validation, Formal Analysis, Investigation, Data Curation, Writing – Review and Editing, Visualization; **Tien Doan**: Resources, Data Curation; **Charles R. Boyle**: Investigation, Data Curation, Writing – Review and Editing; **Dulce Garcia**: Resources, Data Curation; **Jacob W. Class**: Investigation; **Lacy M. Simons**: Data Curation, Supervision, Project Administration; **Margarita Rzhetskaya**: Investigation, Resources; **Alexa Mendoza**: Software, Investigation, Data Curation; **Molly Schnieders**: Resources, Writing – Review and Editing; **Anna Pawlowski**: Resources; **Sameer J. Patel**: Methodology, Software, Resources, Data Curation, Writing – Review and Editing, Supervision, Project Administration; **Ramon Lorenzo-Redondo**: Conceptualization, Methodology, Software, Validation, Data Curation, Writing – Review and Editing, Supervision, Project Administration; & **Judd F. Hultquist**: Conceptualization, Data Curation, Writing – Original Draft, Writing – Review and Editing, Supervision, Project Administration, Funding Acquisition.

---

**Supplementary Figure 1.**
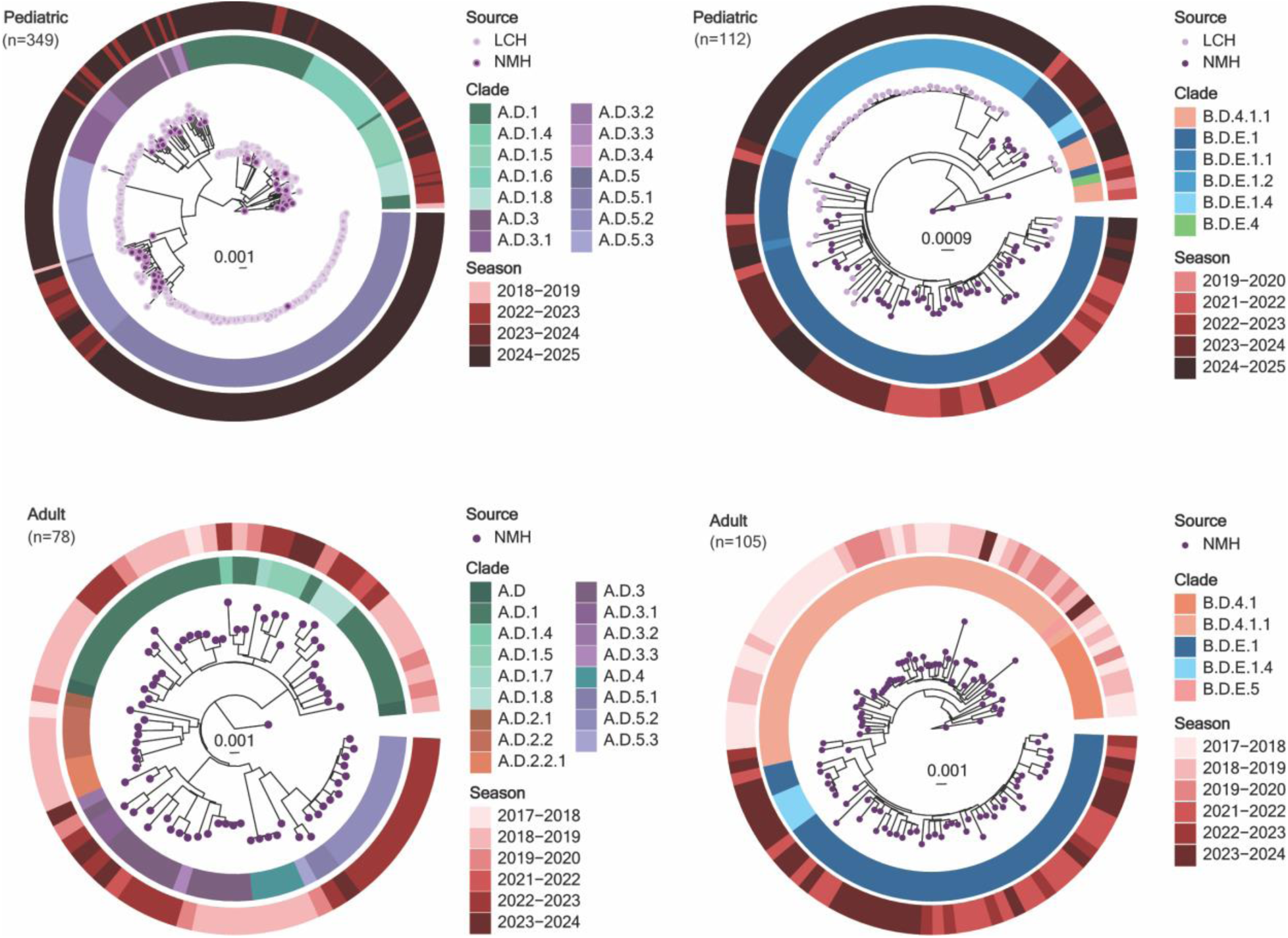
| Historical Sequencing Efforts for adult and pediatric RSV sequences. ML phylogenetic analysis of RSV genomes from NM and LCH in Chicago, Illinois from the 2017 to 2025 RSV seasons (n=644). For all trees, branch tips are colored by patient encounter age group, the inner ring is colored by Nextclade v3.21.0 WGS clade designation, and the outer ring is colored by RSV season in which the isolate is sampled from.

**Supplementary Figure 2.**
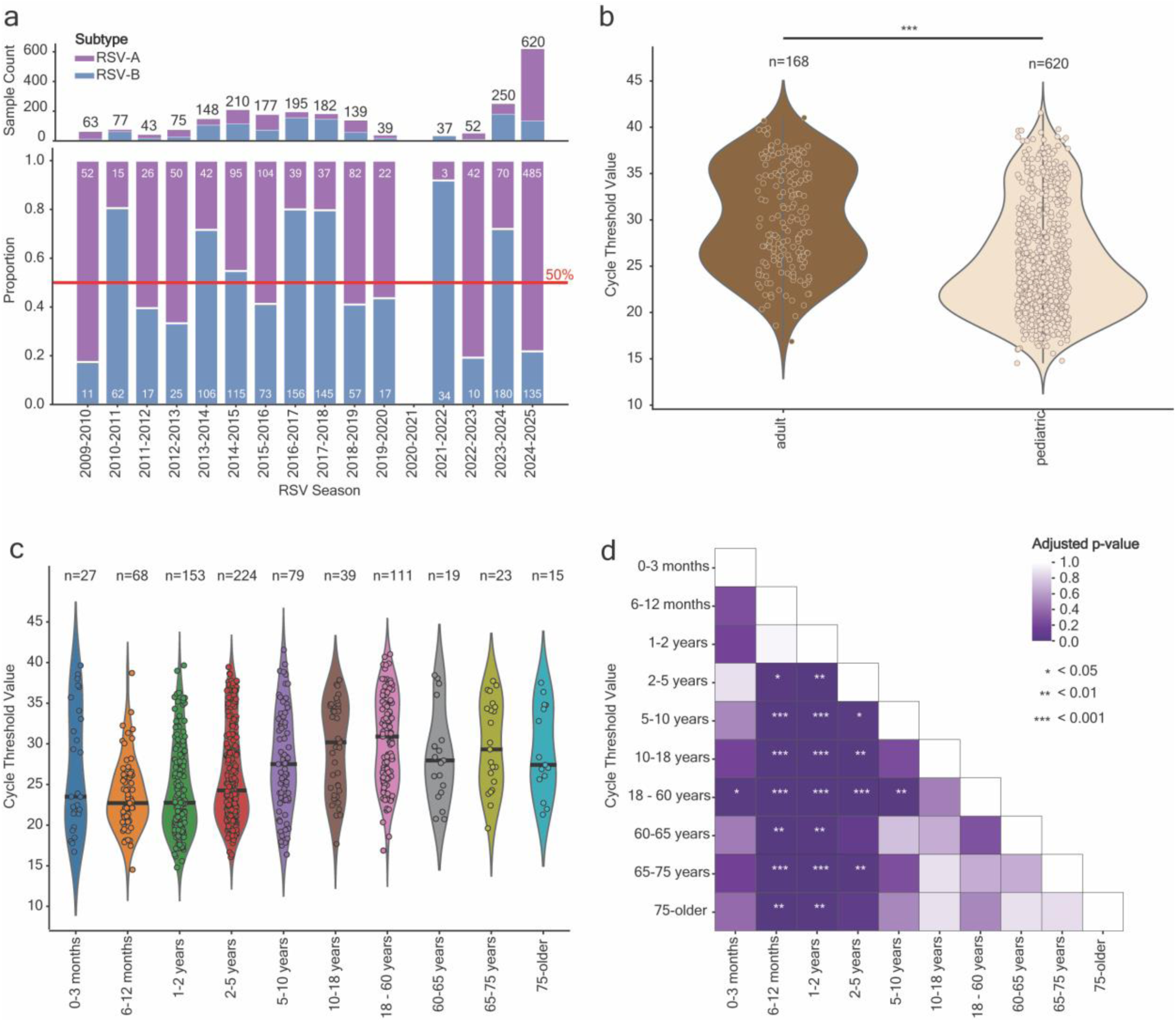
| RSV subtype distribution and viral load characteristics in adult and pediatric populations. **a,** Case count (top) and distribution (bottom) of RSV subtype from patient encounters in NM and LCH from April 2009 to March 2025 among 1,992 specimens with typing information [RSV-A in purple (n=828), RSV-B in blue (n=1164)]. **b,** Violin plot with scatterplot overlay of the Ct distribution among adult (brown, n=168) and pediatric (beige, n=620) RSV diagnostic specimens with available typing data. c, Violin plot with scatterplot overlay of the Ct distributions among patient encounters with available Ct data (n=788) stratified by patient age. **d,** pairwise comparison heatmap of Benjamini–Hochberg adjusted p values. Significant p values denoted by asterisks (* indicates 0.05, ** indicates <0.01, and *** indicates <0.001)

**Supplementary Figure 3.**
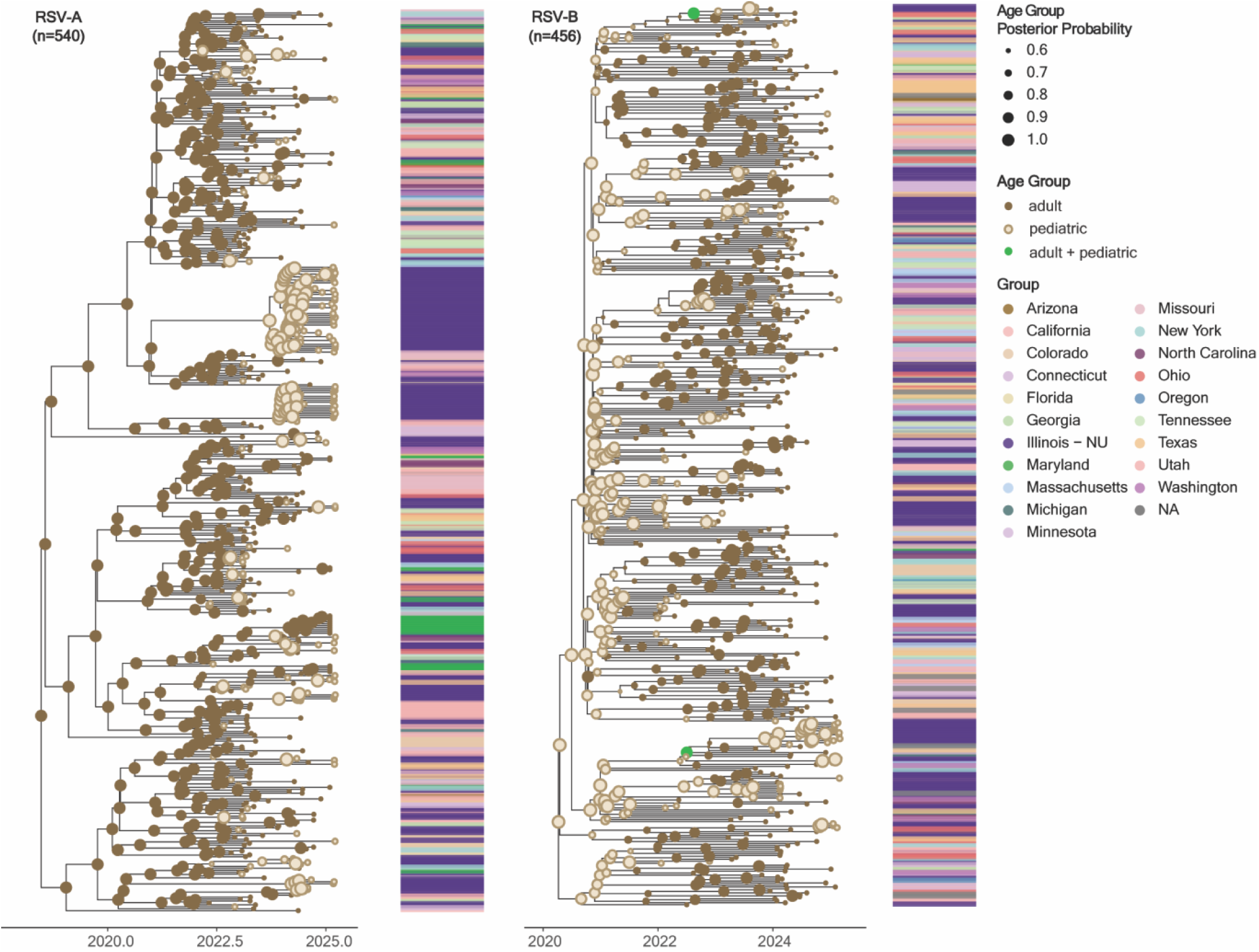
| Temporal Bayesian analysis of in-house and US-based RSV-A and RSV-B sequences by patient age group. Bayesian phylogenetic temporal tree of circulating RSV-A (n=540) and RSV-B (n=456) genomes sampled in-house and from publicly available US genomes with known age group status. For both trees, branch tips are colored by patient age group status, and node color colored by the most probable sampled population (pediatric vs. adult). The size of the node circle represents the probability of the ancestral origin for the node of either pediatric or adult age groups. The bar adjacent to each tree is colored by sampling state.

**Supplementary Figure 4.**
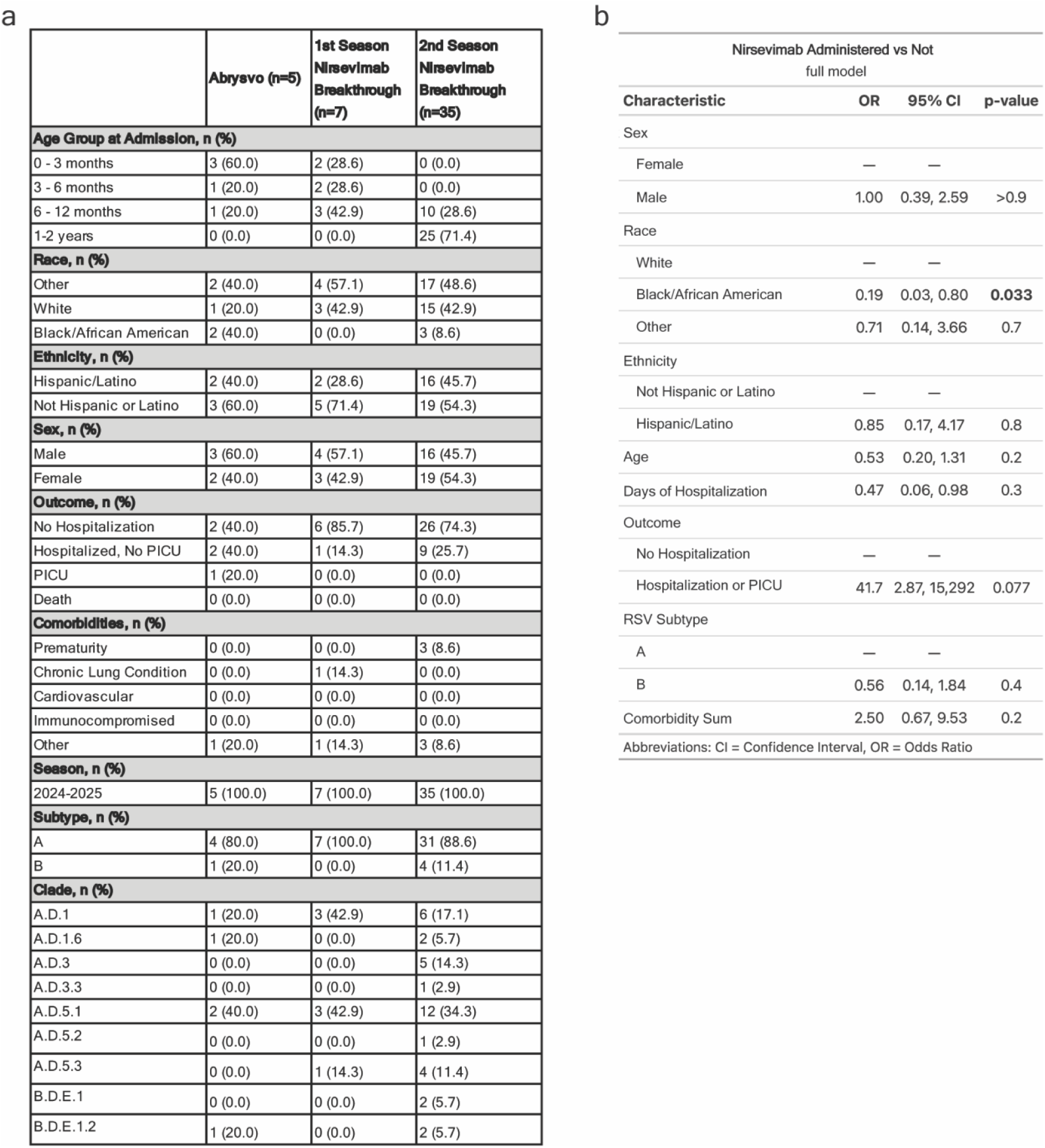
| Demographics of LCH patients with administered RSV prophylaxis and modeling patient outcome by nirsevimab administration. **a,** Demographic and clinical features of pediatric encounters with nirsevimab or Abrysvo-related breakthrough RSV infections. **b,** Demographic and virologic variables incorporated in a multivariable logistic regression to model nirsevimab administration among pediatric patients exposed to infection in their second RSV season (n = 116).

**Supplementary Figure 5.**
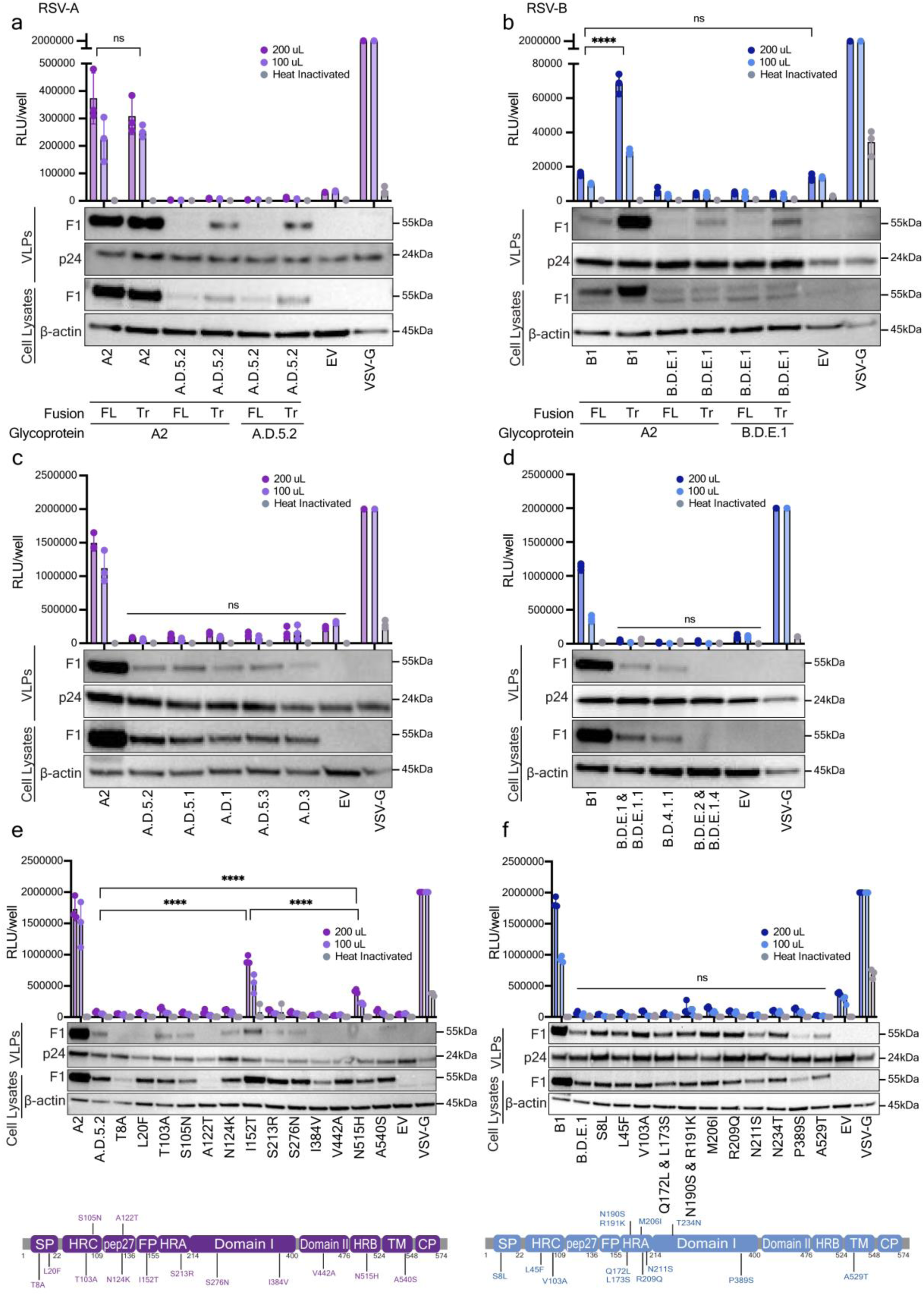
| Optimization of contemporary RSV clades in pseudotyping system. **a,** Lentiviral pseudoviruses bearing full-length (FL) or truncated (Tr) [A2, A.D.5.2] F and [G glycoprotein from RSV A2 with 31 AA deletion or G glycoprotein from A.D.5.2 31 AA deletion] to recover viral entry activity **b,** Lentiviral pseudoviruses bearing full-length (FL) or truncated (Tr) [B1, B.D.E.1] F and [G glycoprotein from RSV A2 with 31 AA deletion or G glycoprotein from B.D.E.1 31 AA deletion] to recover viral entry activity **c,** Truncated (Tr) pseudoviruses constructs based on A.D.5.2 backbone were engineered to contain individual A2 -specific point mutations using site-directed mutagenesis (SDM). **d**, Truncated (Tr) pseudoviruses constructs based on B.D.E1 backbone were engineered to contain individual B1 -specific point mutations using SDM. **e,** Lentiviral pseudoviruses bearing truncated [A2, A.D.5.2, A.D.5.1, A.D.1, A.D.5.3, A.D.3] F to confirm no activity in any contemporary clades. **f,** HEK293T effector cells were transiently transfected with plasmids encoding truncated [A2, A.D.5.2, A.D.5.1, A.D.1, A.D.5.3, A.D.3] F with and without mutation (I152T), as well as a T7 polymerase. Target cells were transfected with T7 promoter-driven luciferase reporter effector HEK293T cells were mixed at a 1:1 ratio with target HEK293T cells and co-cultured fusion-dependent luciferase activity was measured 48 HPI. **a-d**, all pseudoviruses were quantified and normalized to p24 levels before challenging. Viral entry was quantified by luciferase activity at 72 hours post-infection. Cell lysates (CL) and purified virus-like particles (VLPs) normalized to p24 were analyzed by immunoblotting for each RSV F. p24 was used as a VLP loading control, and β-actin as a cellular loading control. Data points represent the mean ± s.d. from three independent biological replicates. Statistics were calculated using two-way ANOVA followed by Tukey’s multiple comparisons test. Pointed brackets indicate statistical comparisons between two specific conditions. Flat horizontal lines denote comparisons across all conditions. ns, not significant; p < 0.05 (\**); p < 0.01 (*\*\*); p < 0.001 (***); p < 0.0001 (****).

**Supplementary Figure 6.**
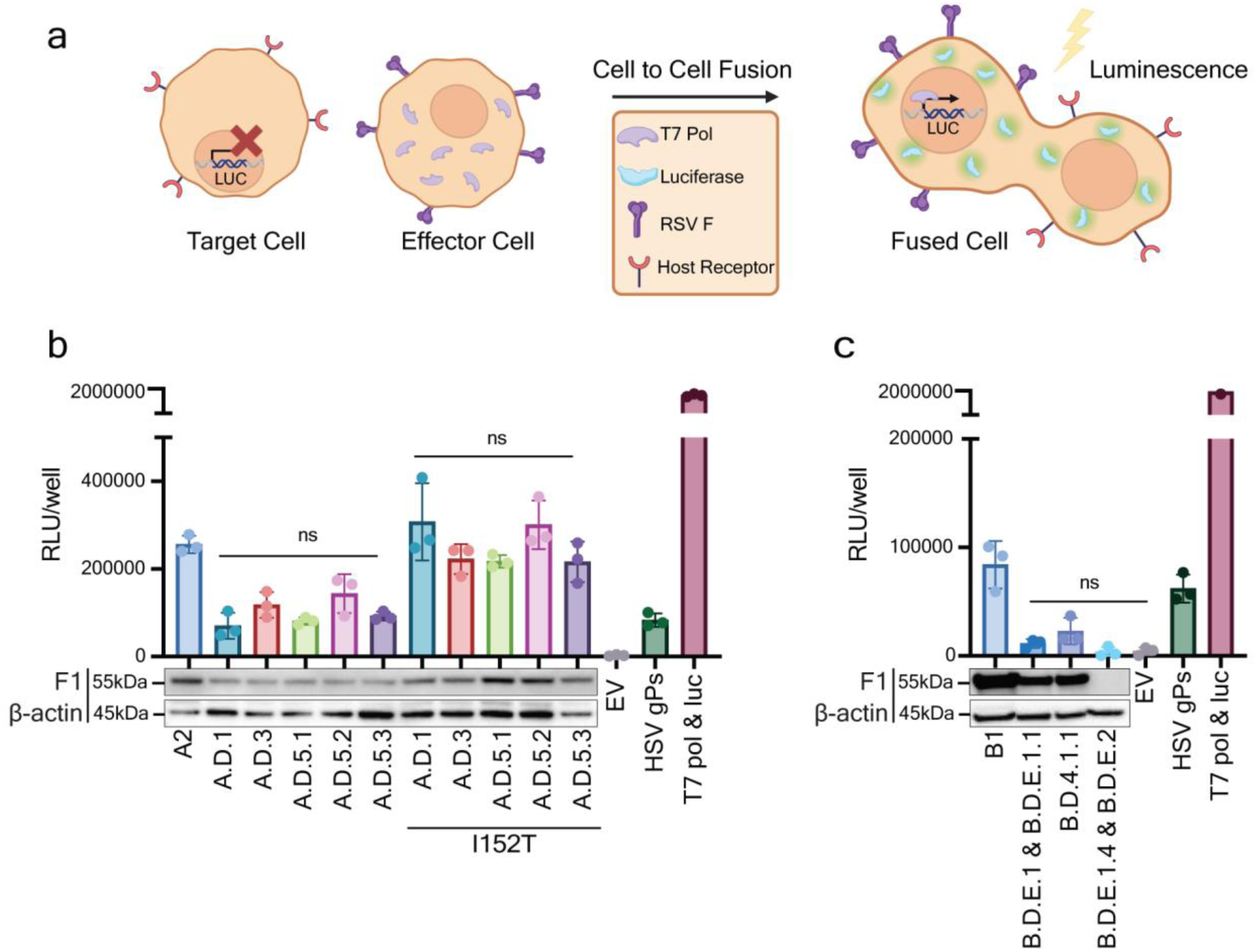
| Characterization of contemporary RSV F activity through Fusion assay. **a**, A schema of the RSV Fusion Assay. HEK293T target cells were transfected with T7 promoter-driven luciferase reporter. HEK293T effector cells were transiently transfected with plasmids encoding viral F glycoproteins together as well as T7 polymerase. After 24-hour transfection, HEK293T effector cells were mixed at a 1:1 ratio with HEK293T target cells and co-cultured. Fusion was quantified via luminescence 48-hour post-infection (HPI). **b**, Effector cells were transiently transfected with plasmids encoding truncated [A2, A.D.5.2, A.D.5.1, A.D.1, A.D.5.3, A.D.3] F with and without mutation (I152T), as well as T7 polymerase. Target cells were transfected with T7 promoter-driven luciferase reporter. Target and effector cells were mixed at a 1:1 ratio and co-cultured. Fusion-dependent luciferase activity was measured 48 HPI. **c**, Effector cells were transiently transfected with plasmids encoding truncated [B1, B.D.E.1 & B.D.E.1.1, B.D.4.1.1, B.D.E.1.4 & B.D.E.2] F, as well as a T7 polymerase. Target cells were transfected with T7 promoter-driven luciferase reporter. Target and effector cells were mixed at a 1:1 ratio and co-cultured. Fusion-dependent luciferase activity was measured 48 HPI. **b-c**, Cell lysates (CL) were analyzed by immunoblotting for each RSV F. β-actin was used as a cellular loading control. Data points represent the mean ± s.d. from three independent biological replicates. Statistics were calculated using one-way ANOVA followed by Tukey’s multiple comparisons test. Flat horizontal lines denote comparisons across all conditions. ns, not significant; p < 0.05 (*); p < 0.01 (**); p < 0.001 (***); p < 0.0001 (****).

**Supplementary Table 1.**
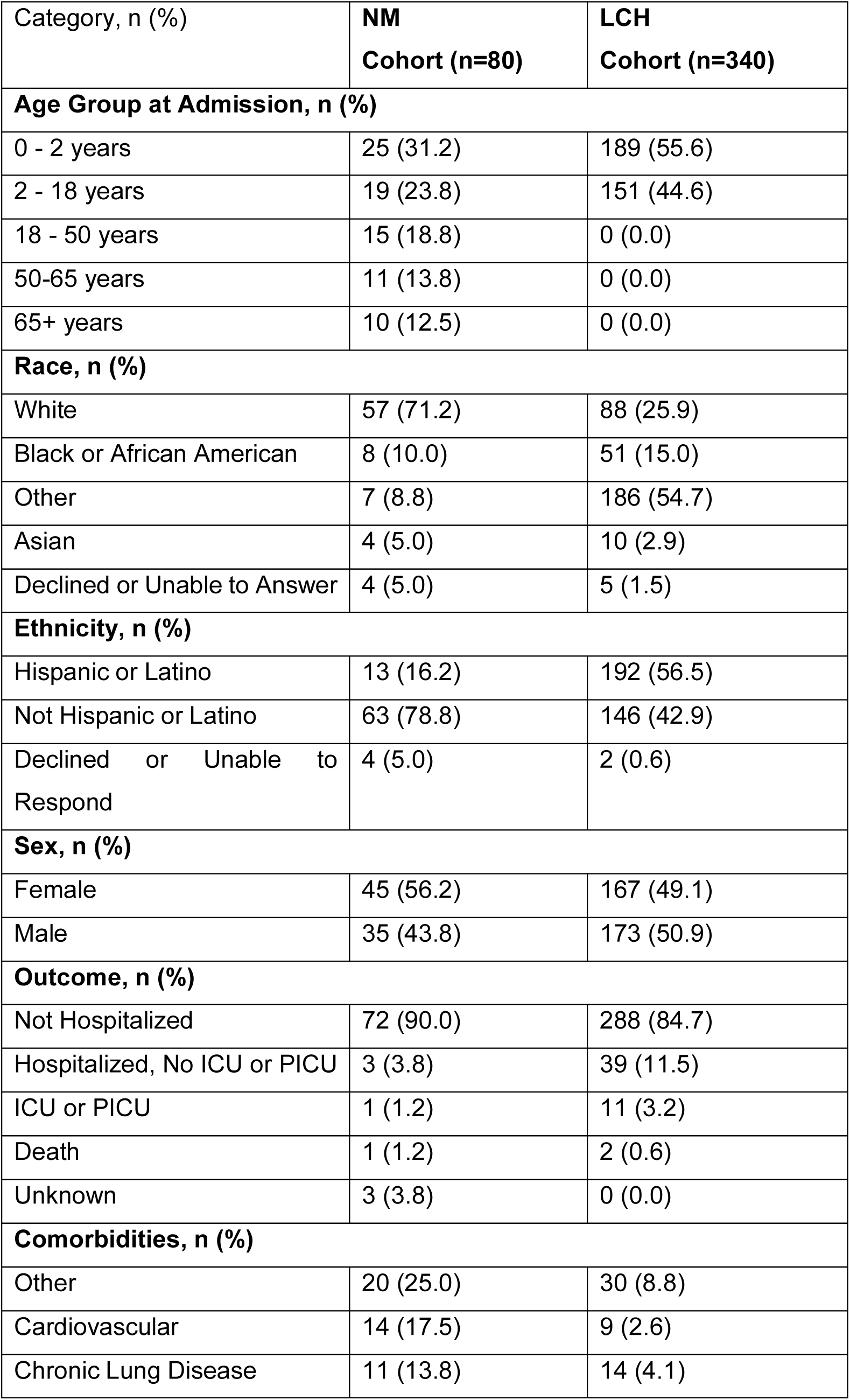
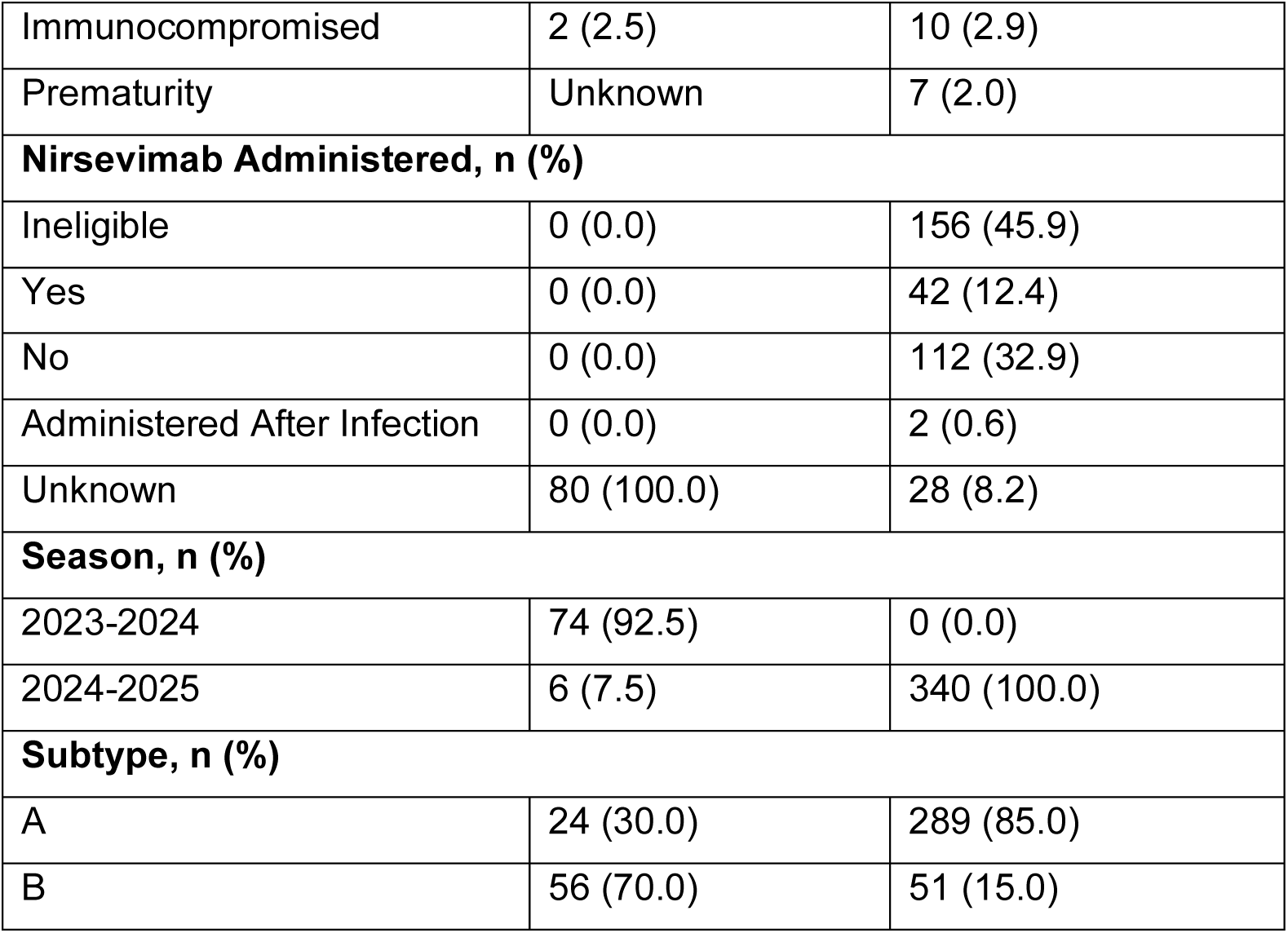
| 2023-2025 RSV cohort demographics. Demographic and clinical features of RSV-positive patients with paired RSV whole genome sequences from NM hospital systems (n=80, left) and LCH (n=338, right). * Nirsevimab eligibility includes infants under 8 months of age encountering their first RSV season between October 1^st^ and March 31^st^ of each year as recommended by the American Academy of Pediatrics.

**Supplementary Table 2.**
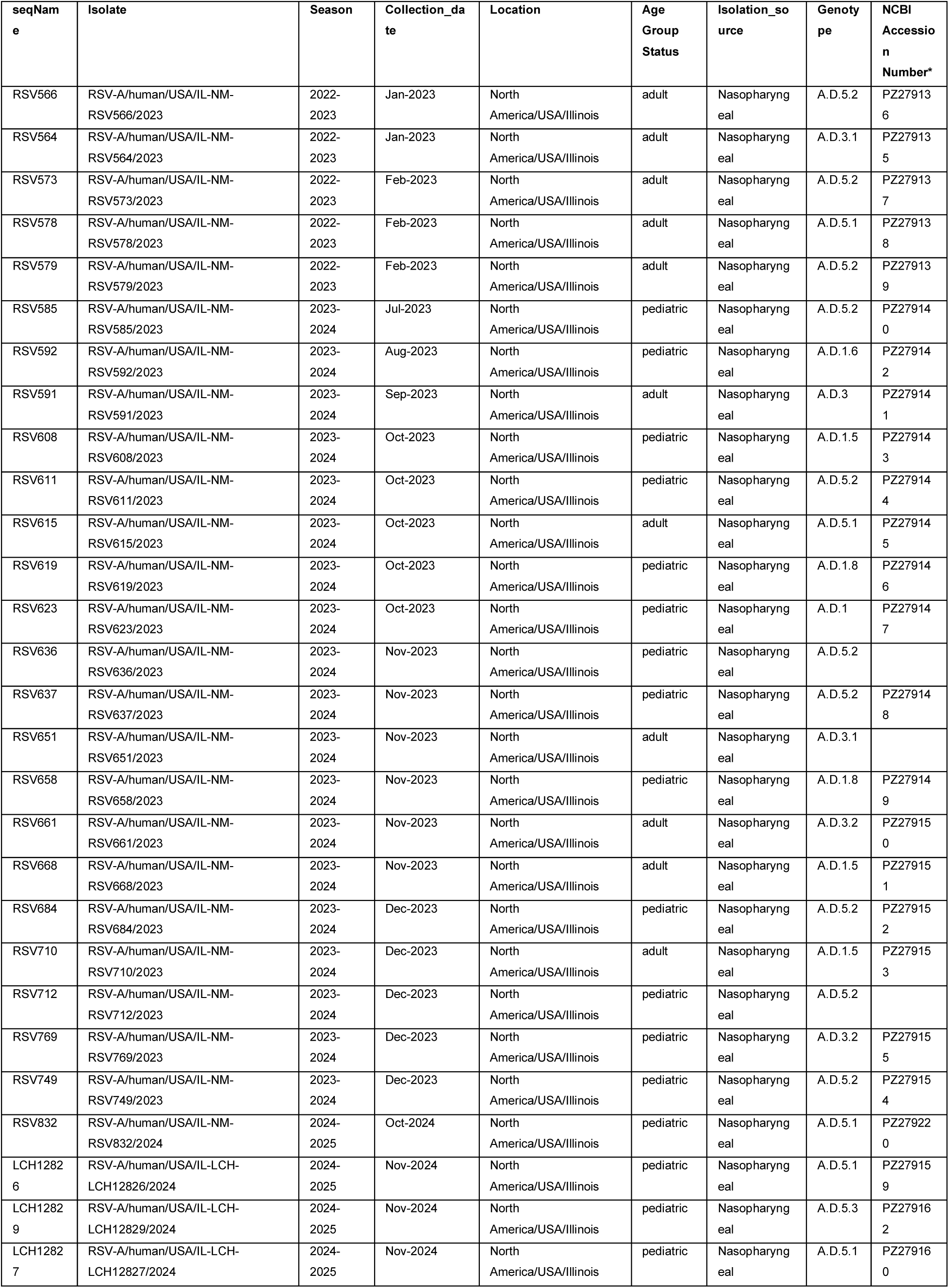
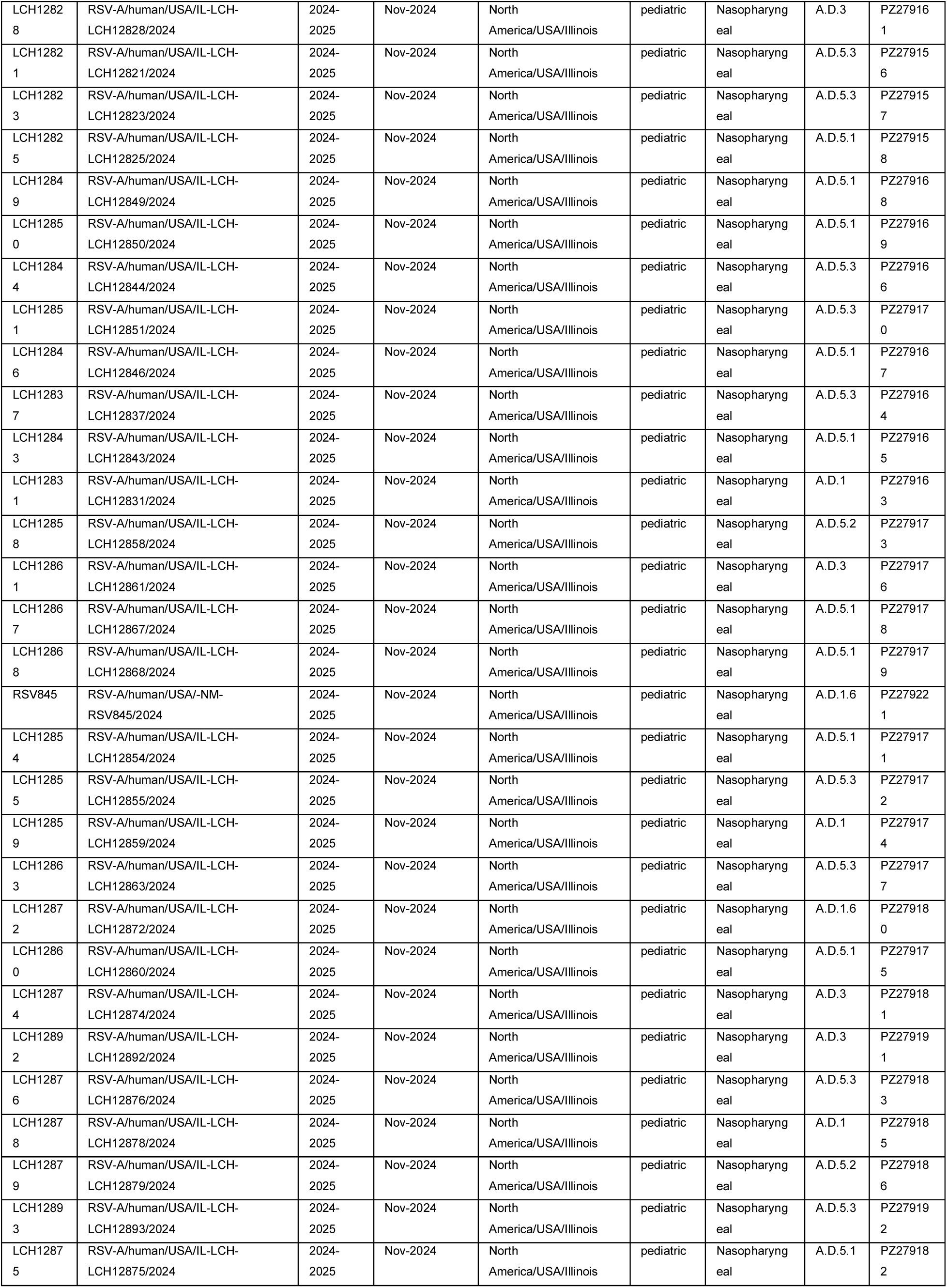
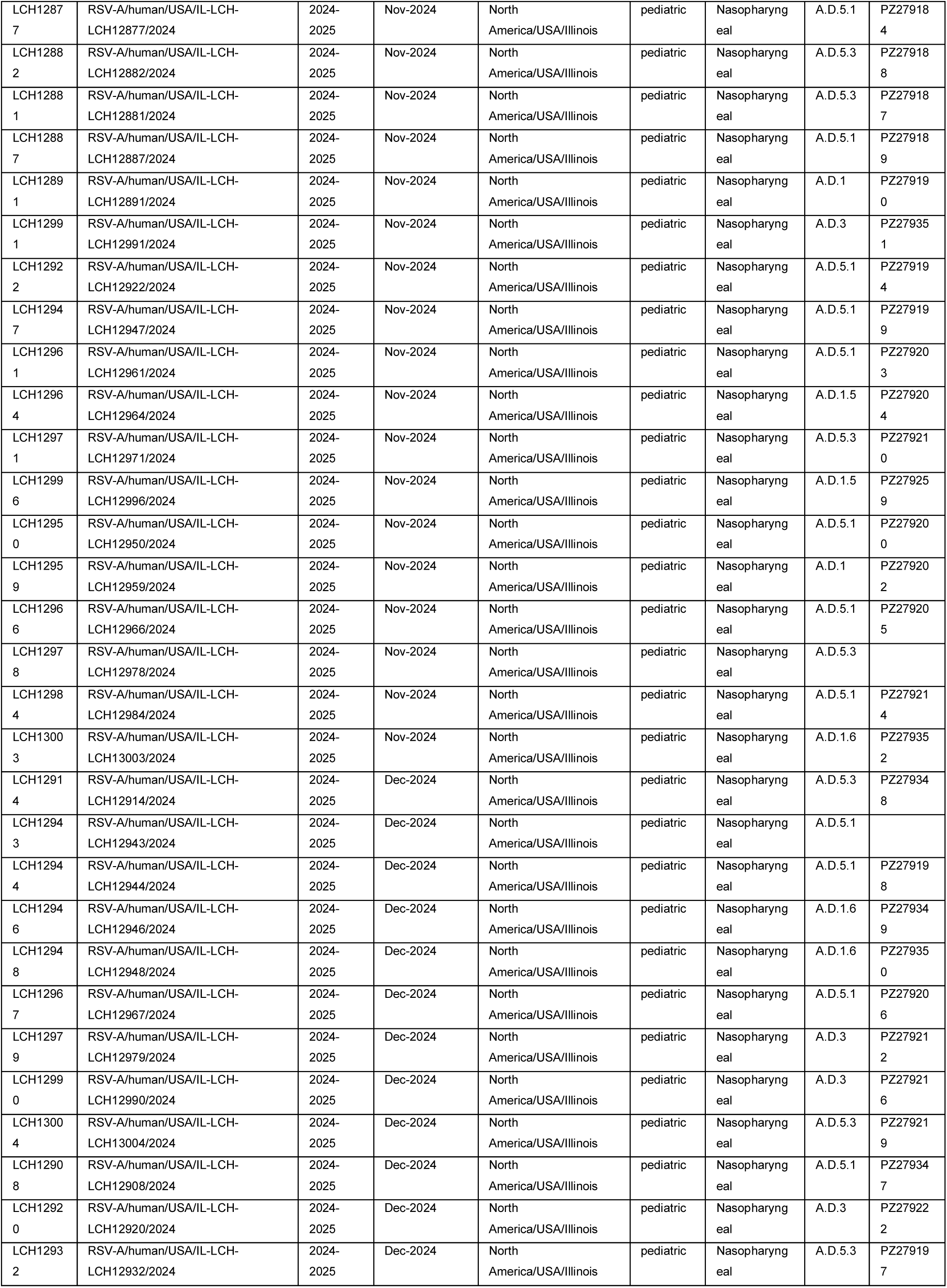
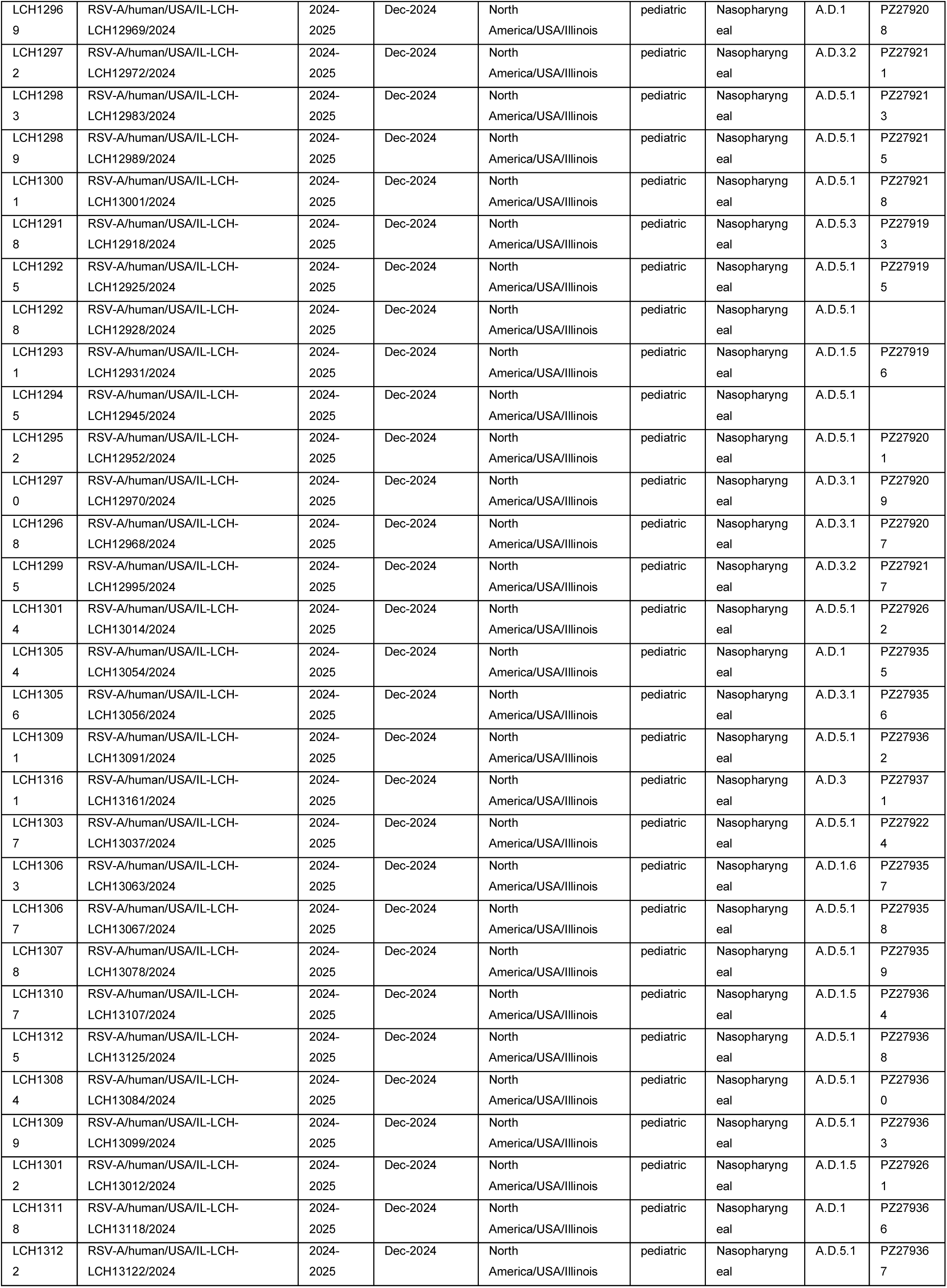
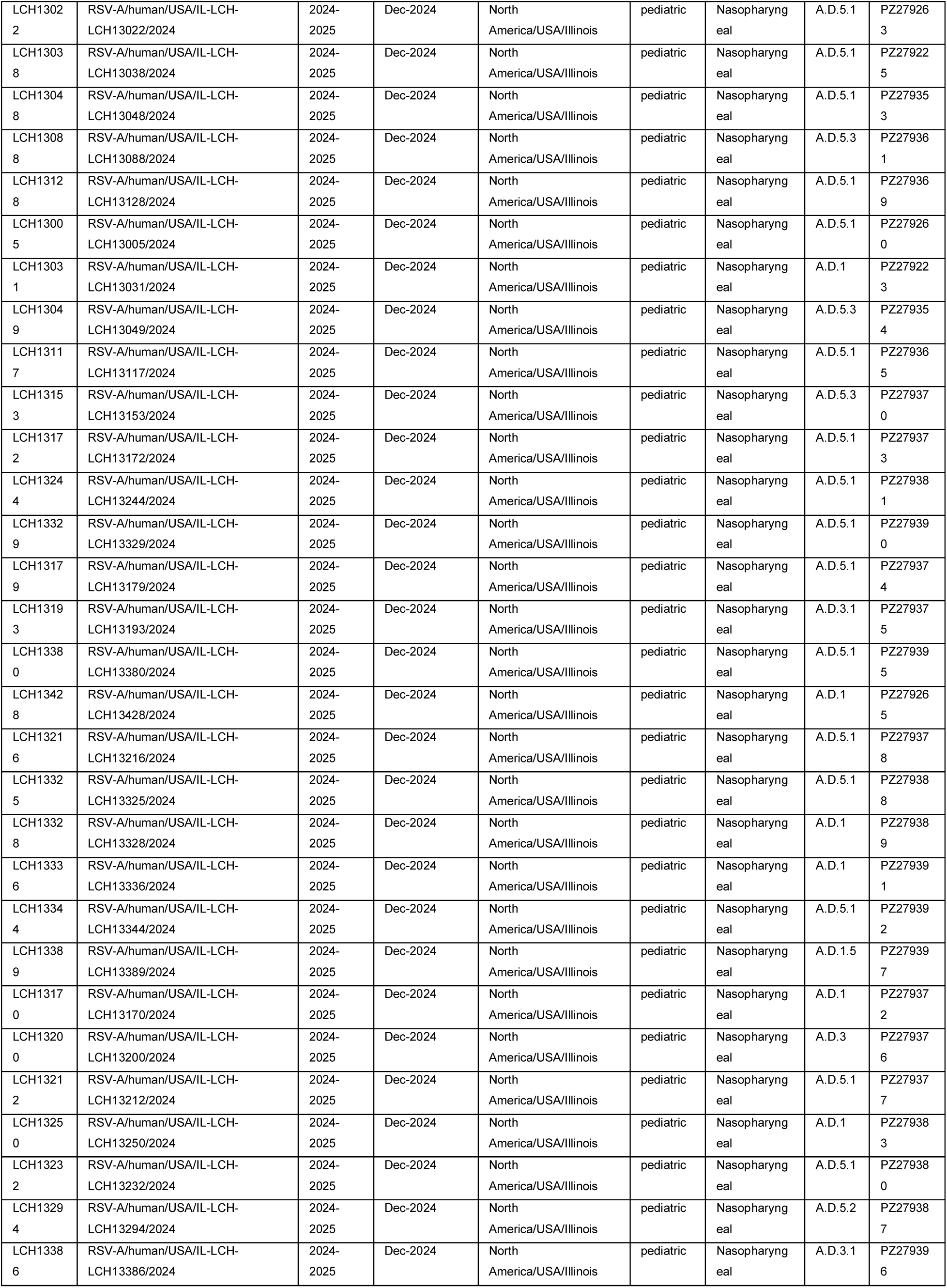
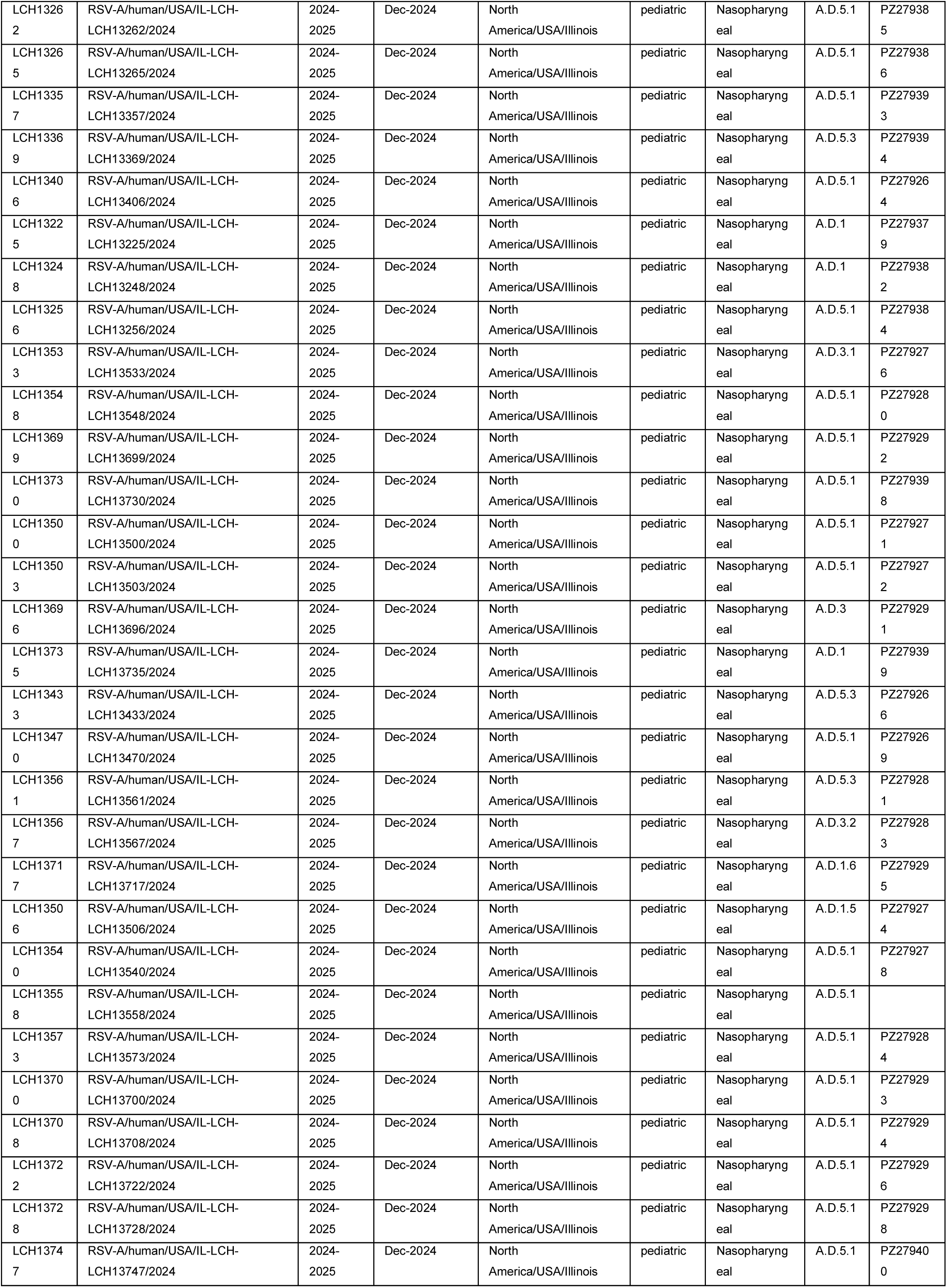
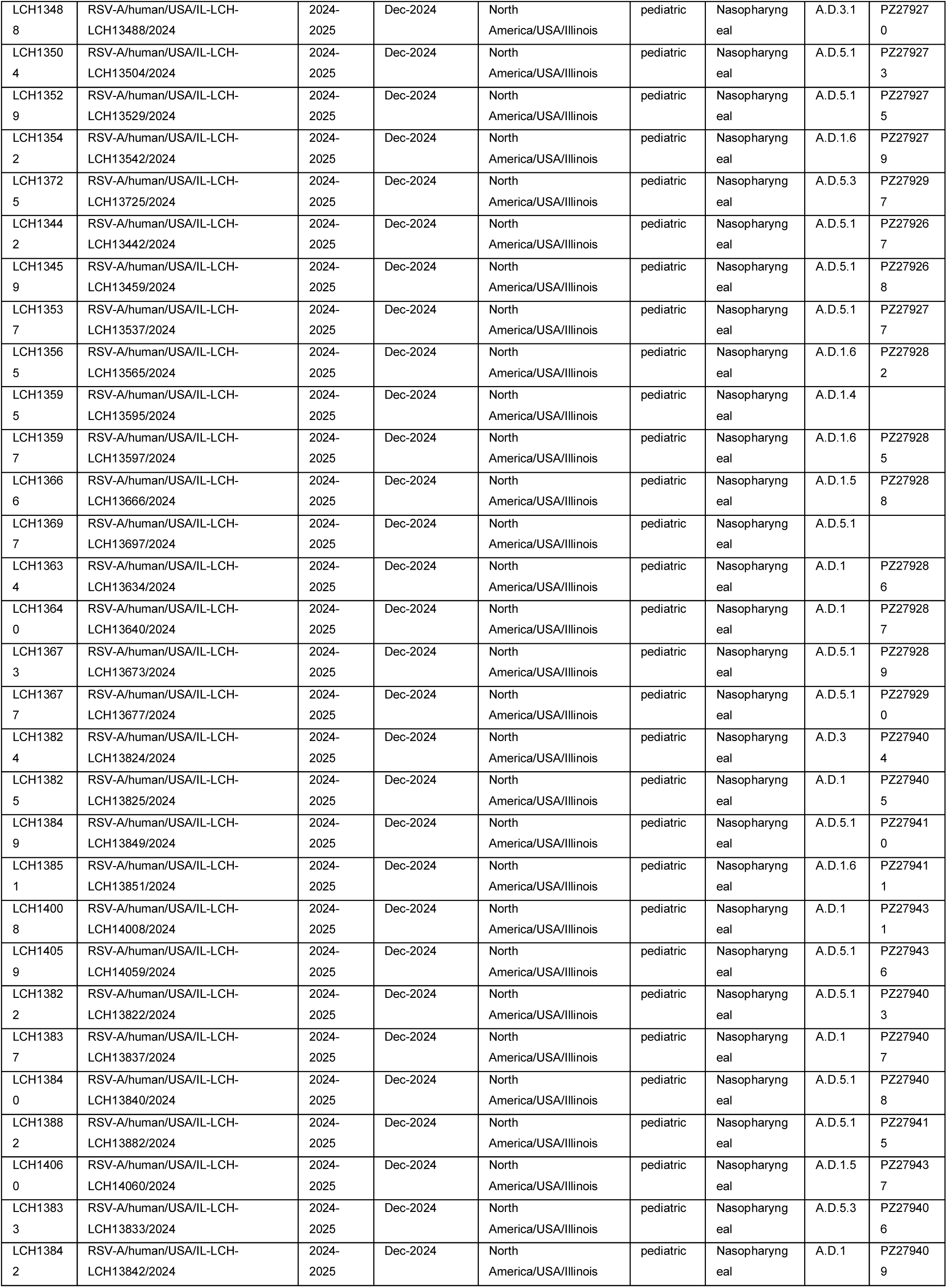
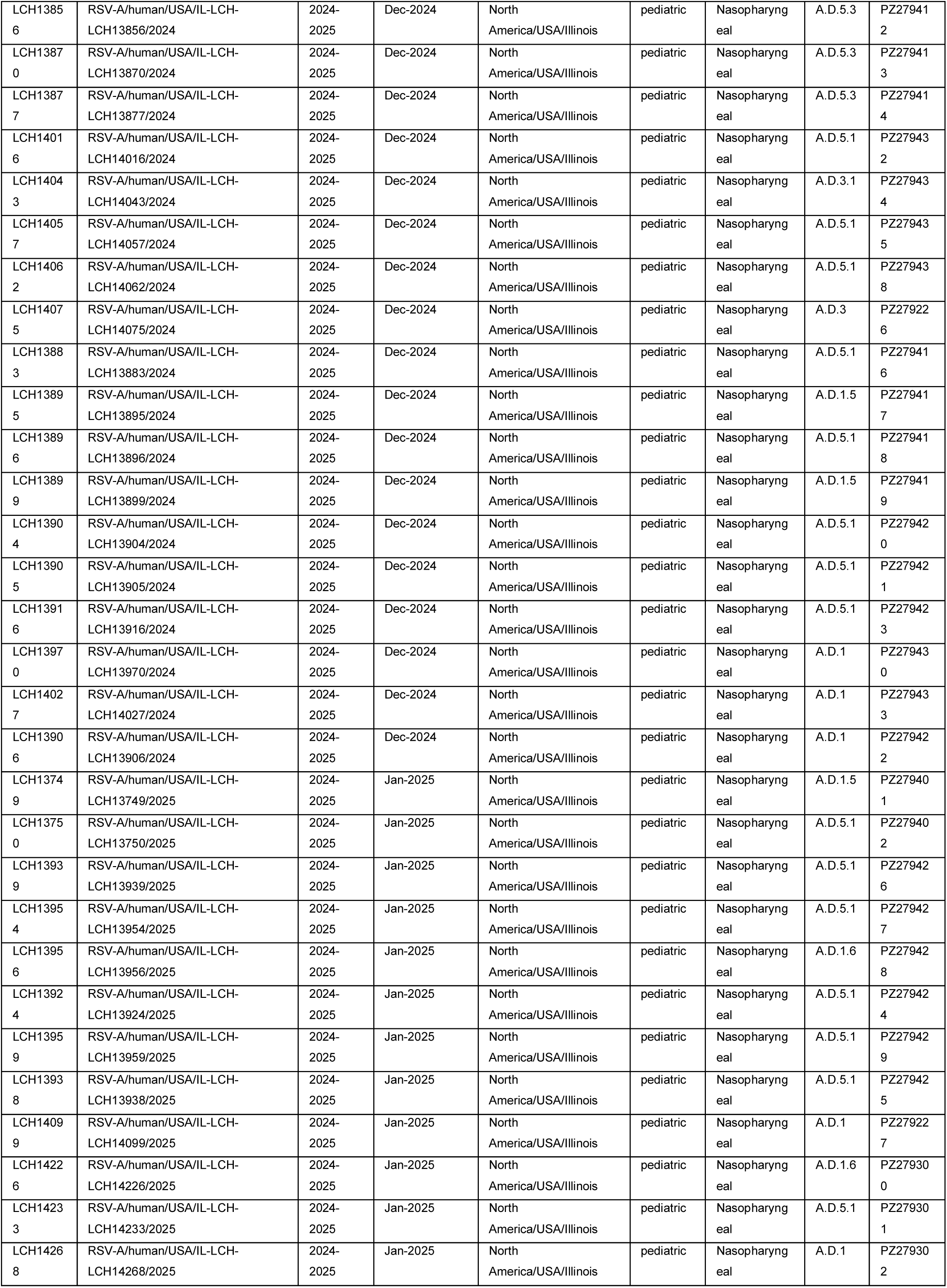
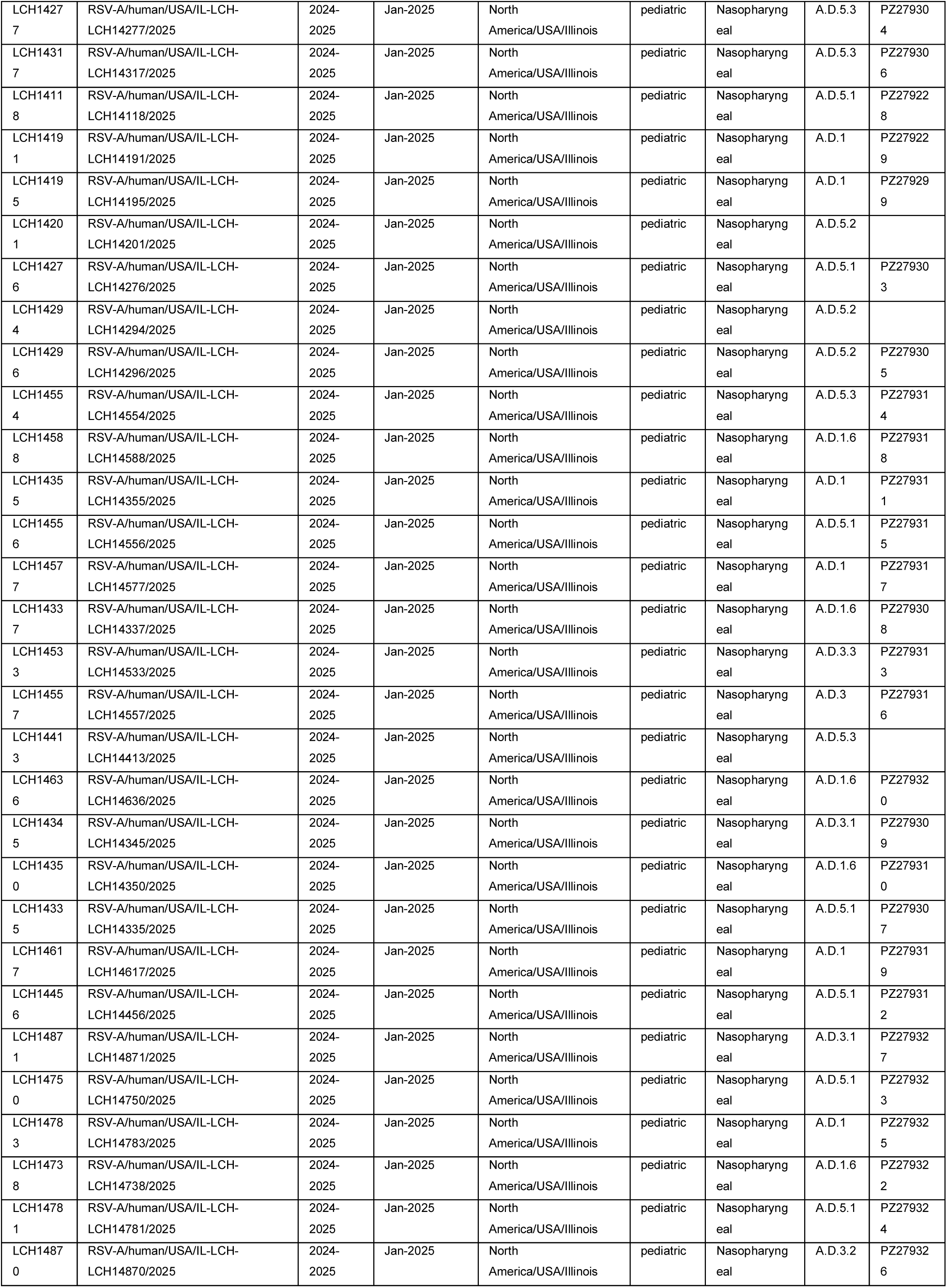
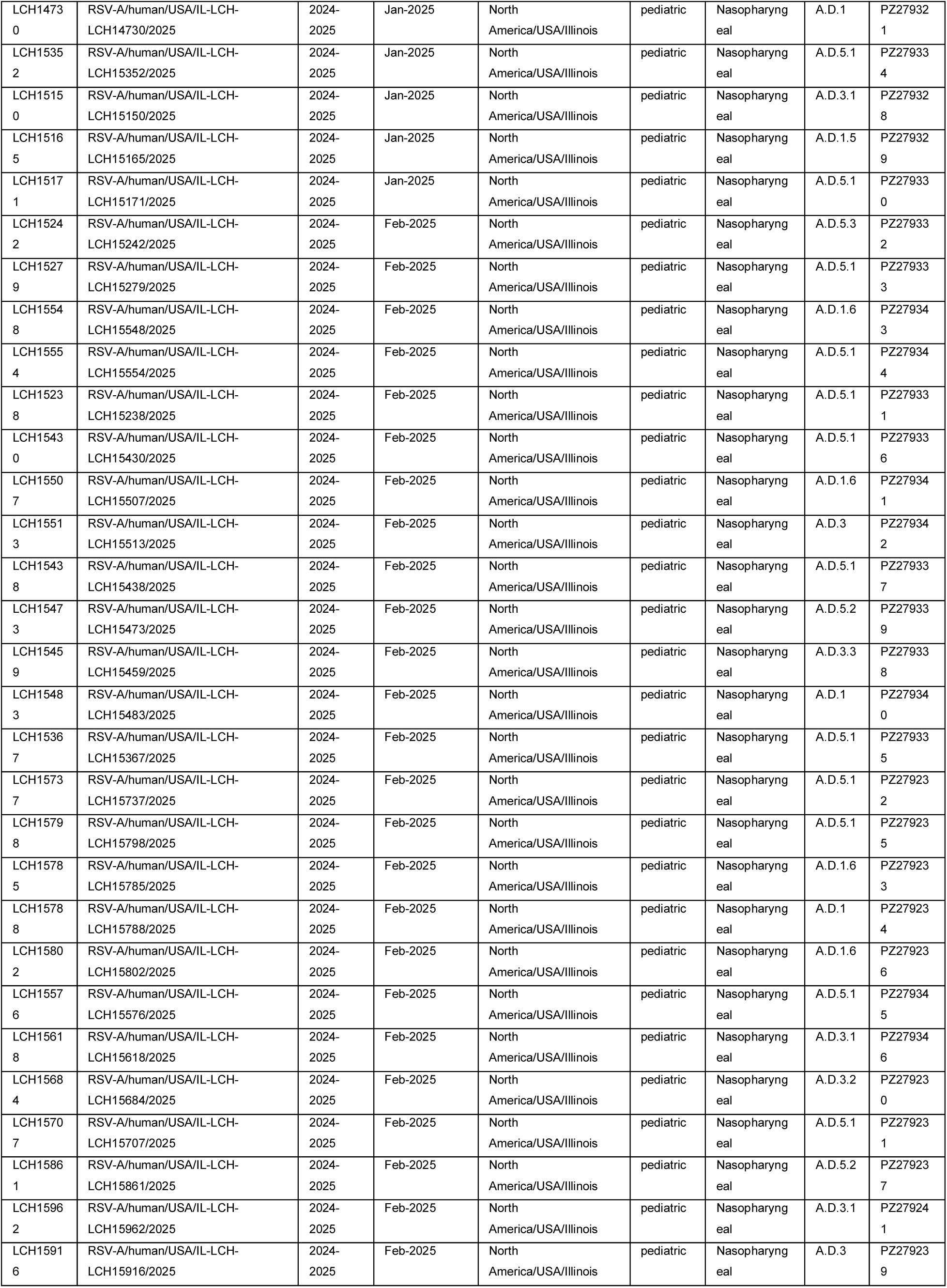
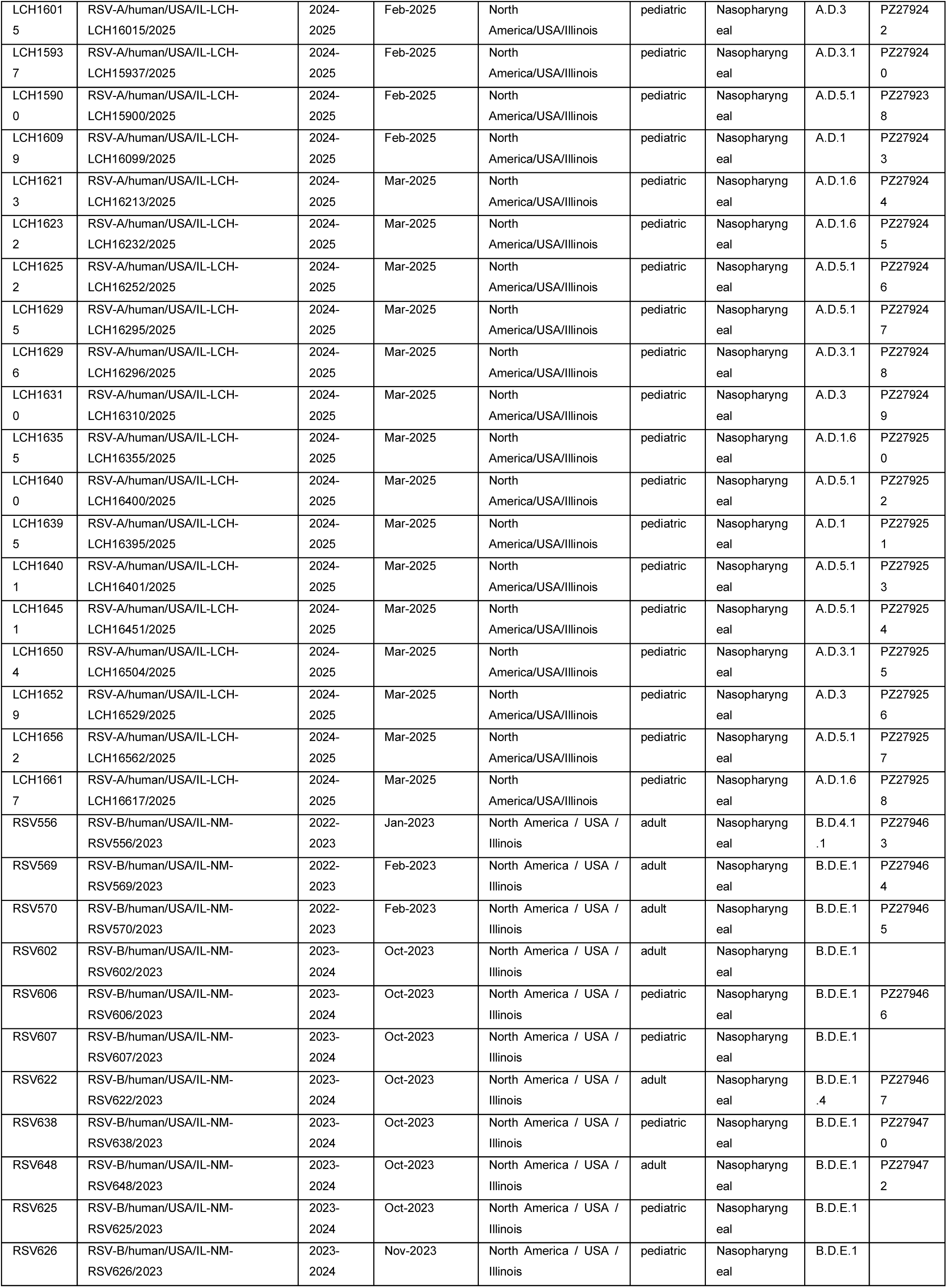
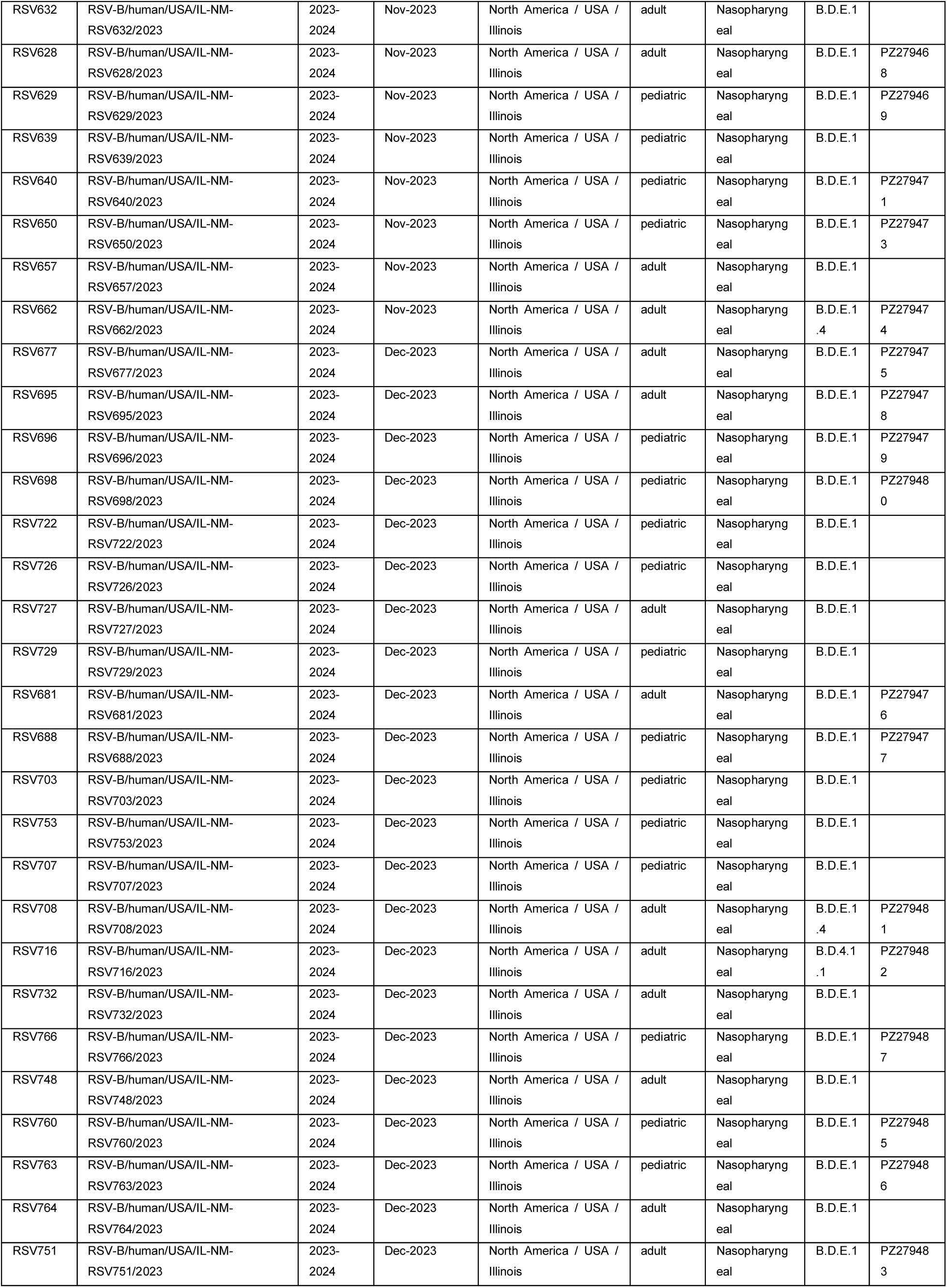
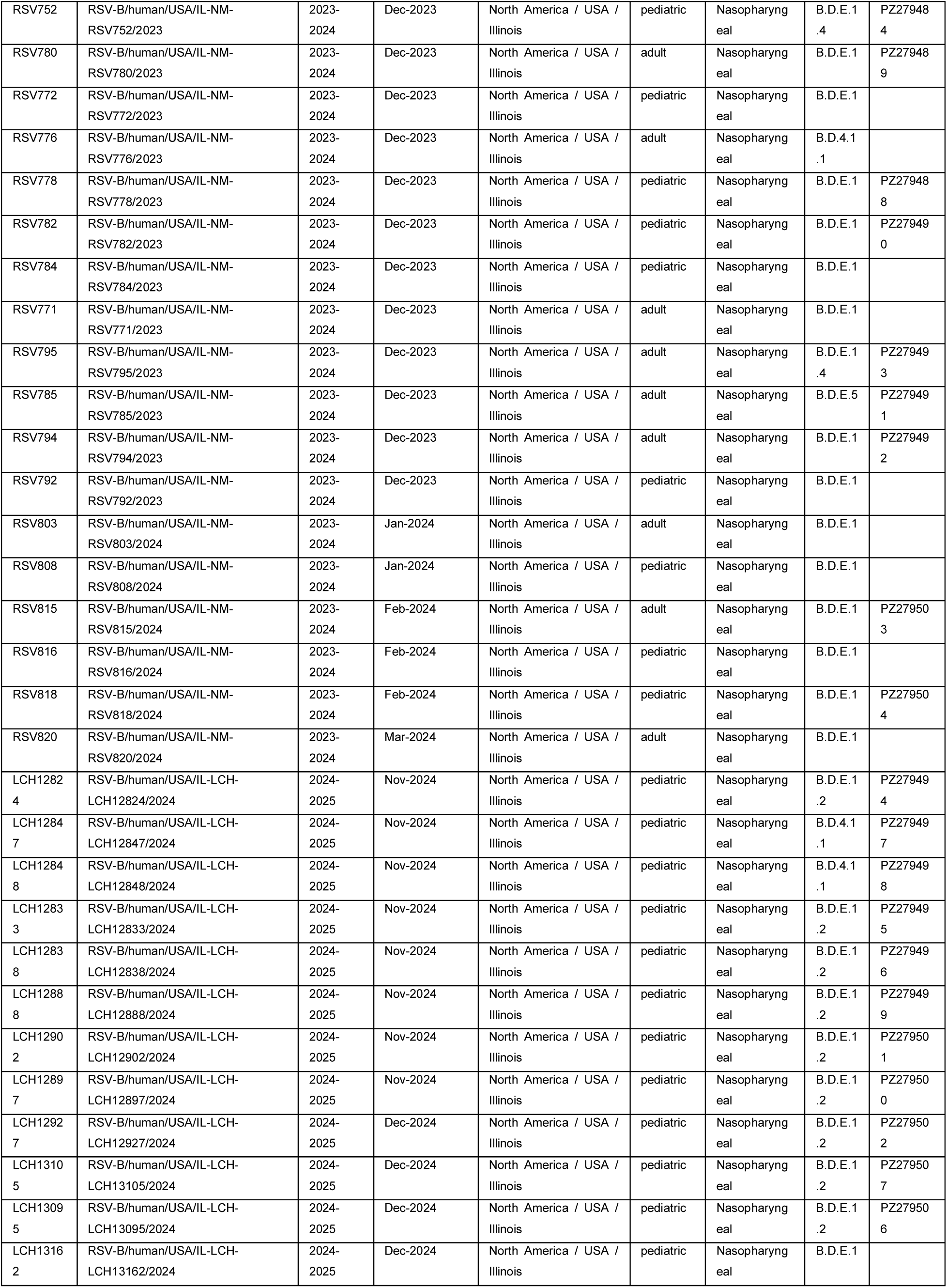
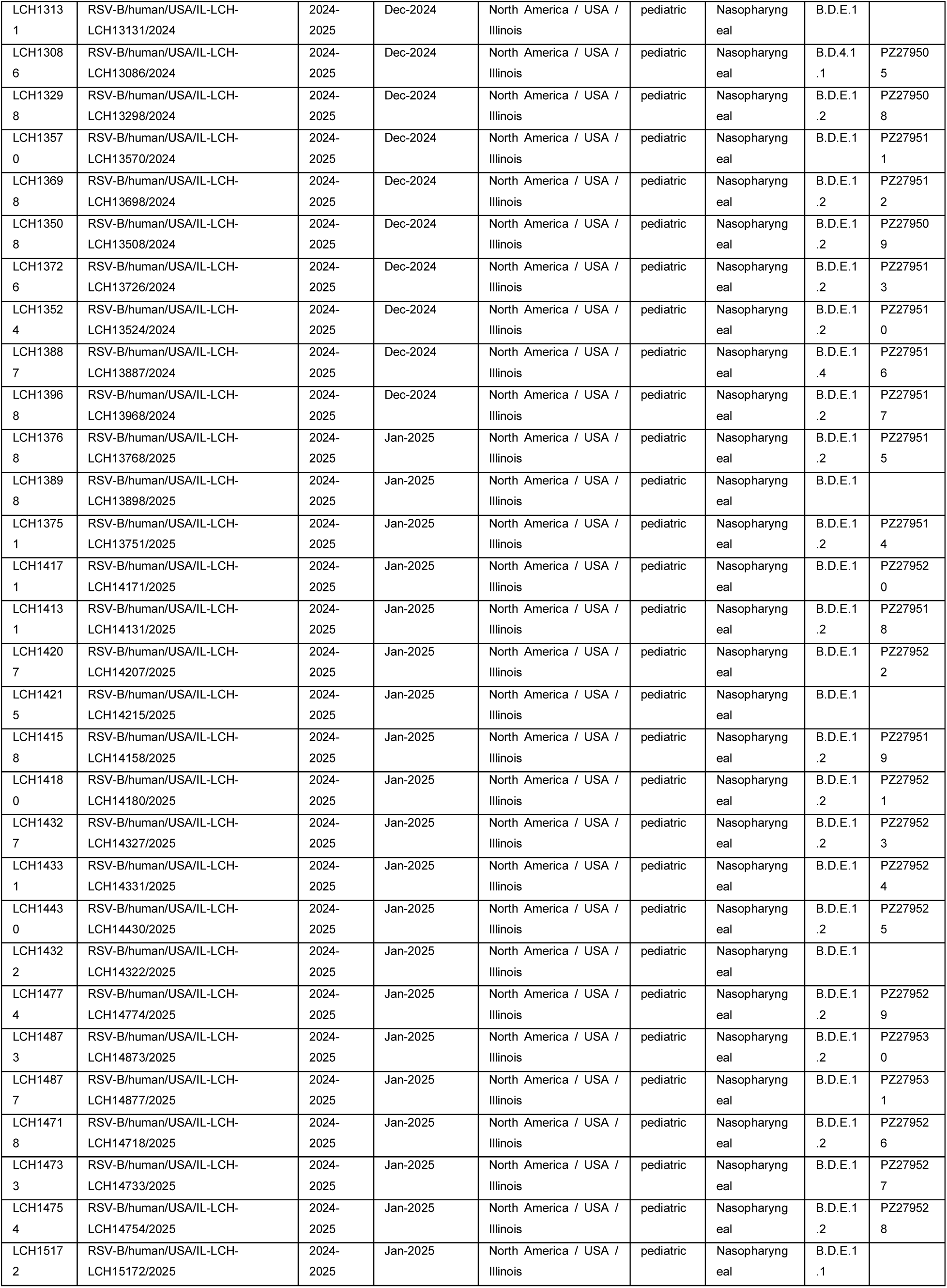
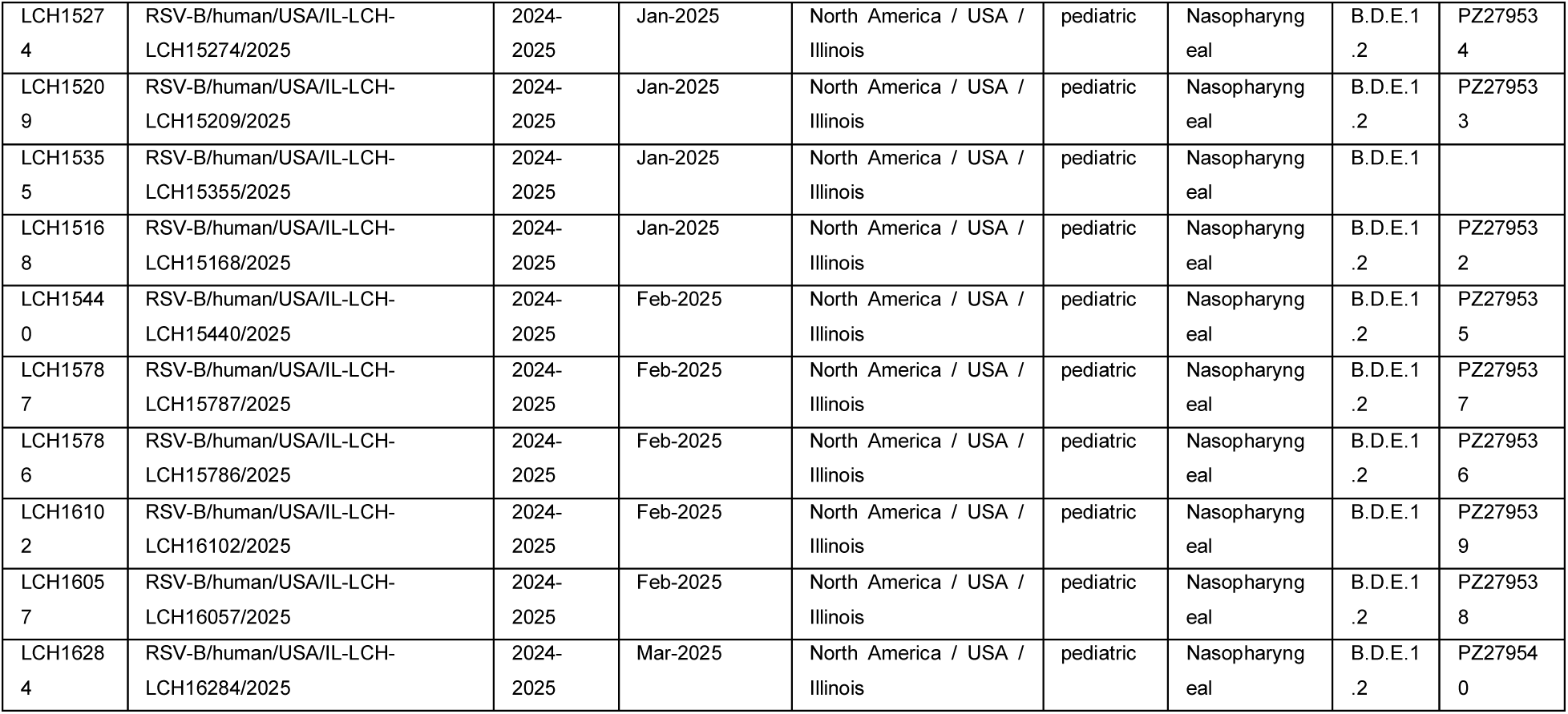
| NCBI accession ID numbers for deposited RSV whole genome sequences. *Note several sequences are in queue for NCBI sequence submission.

**Supplementary Table 3.**
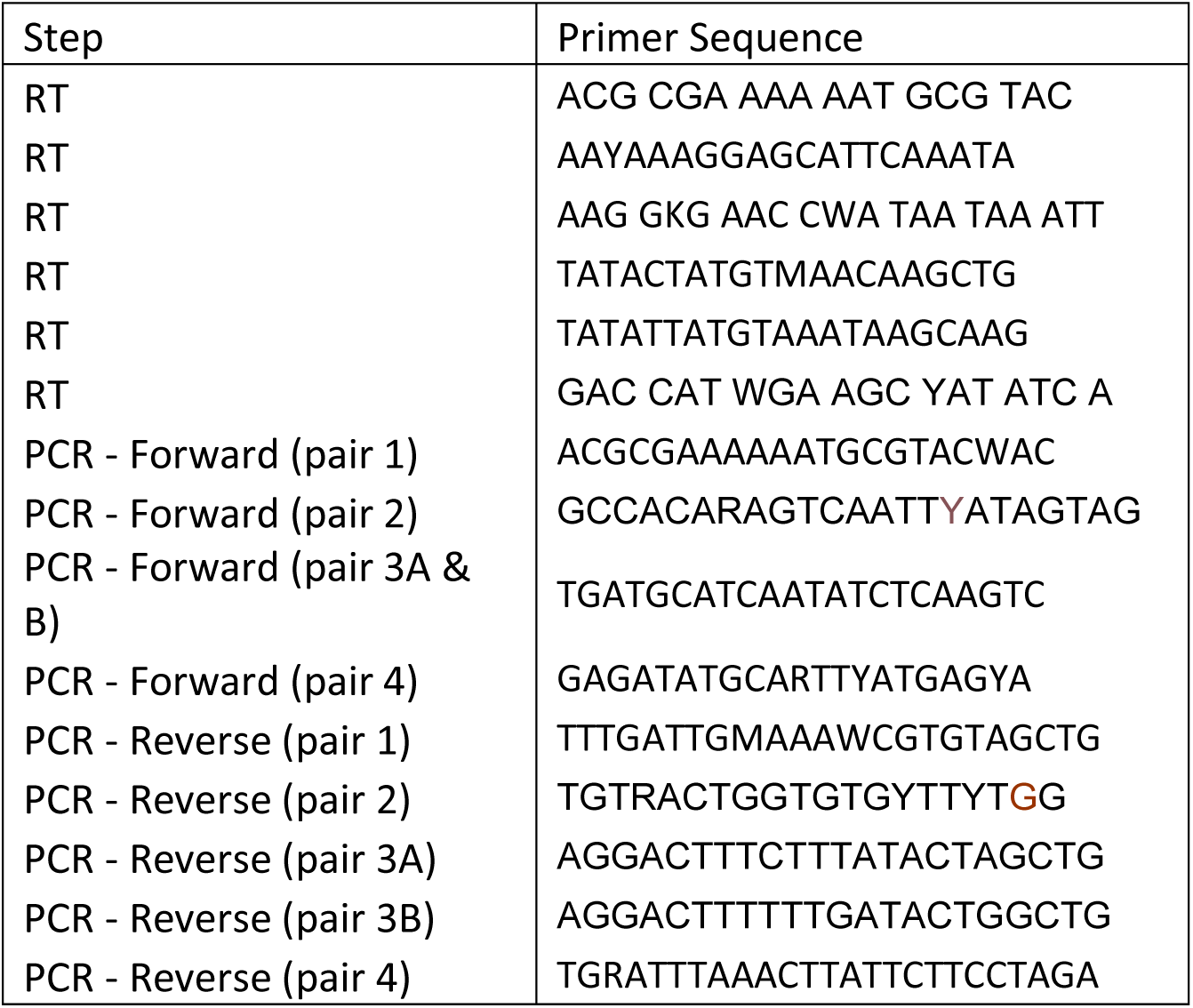
| Primer information for RSV whole genome sequencing pipeline.

**Supplementary Table 4.**
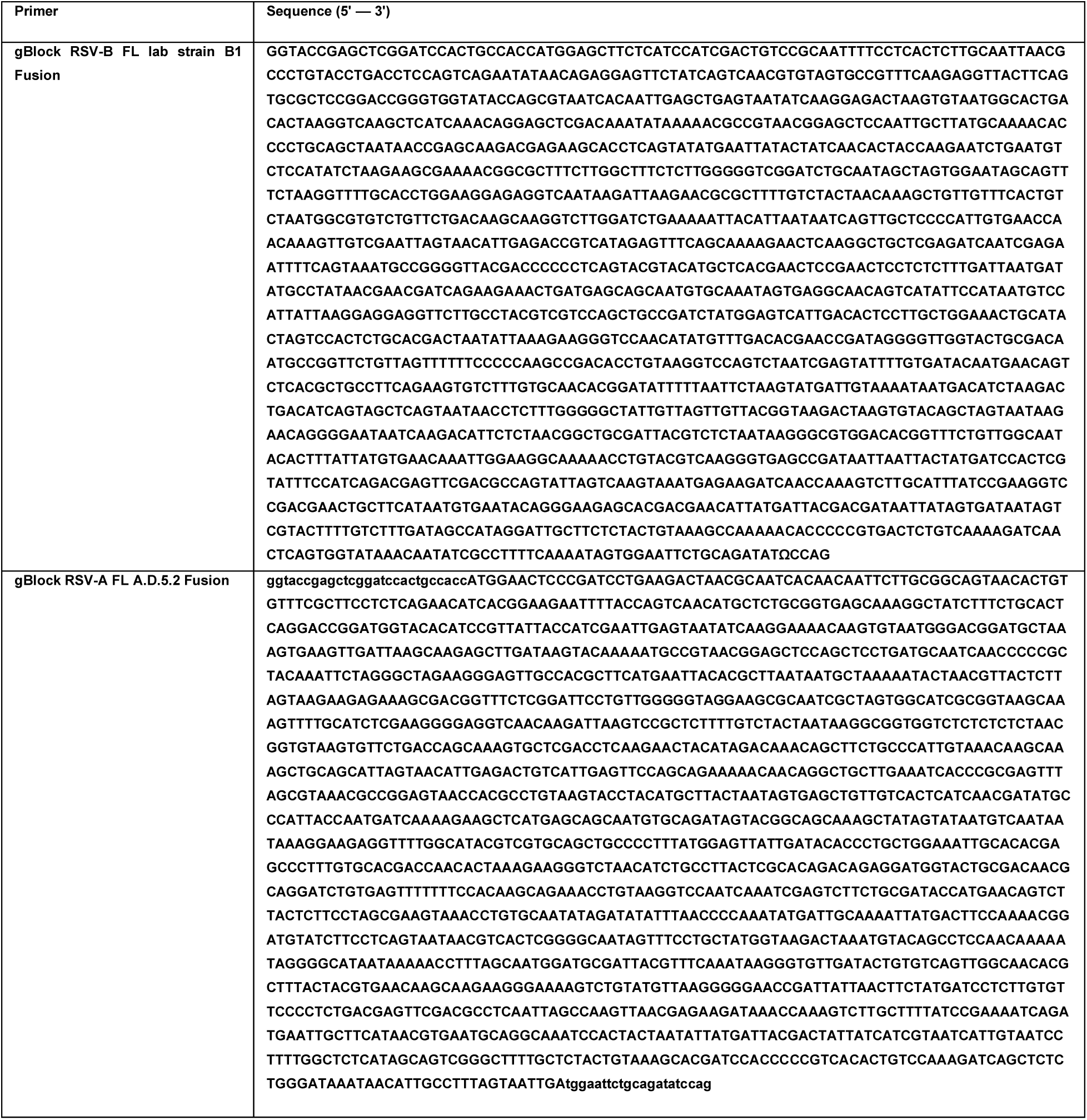
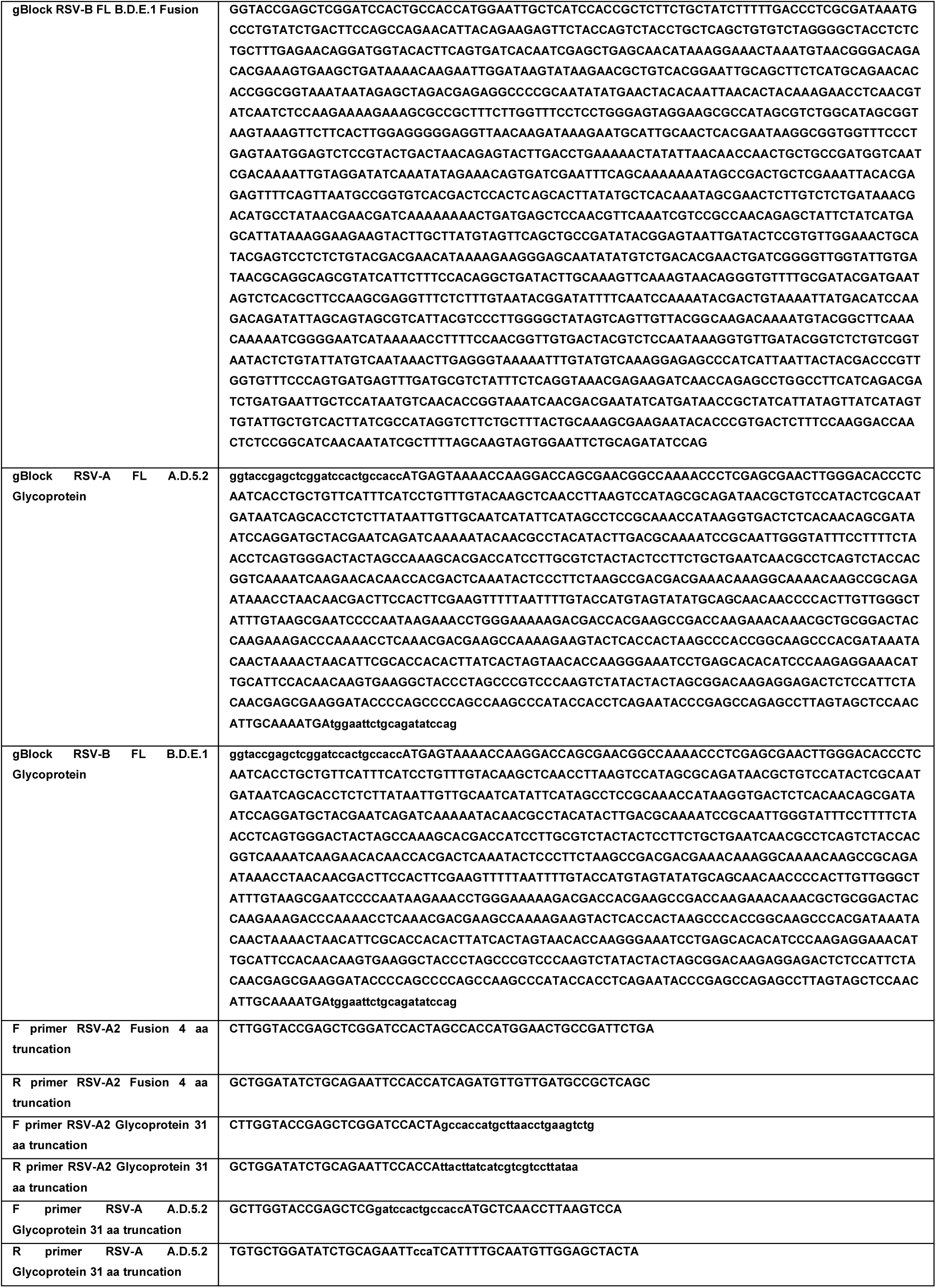
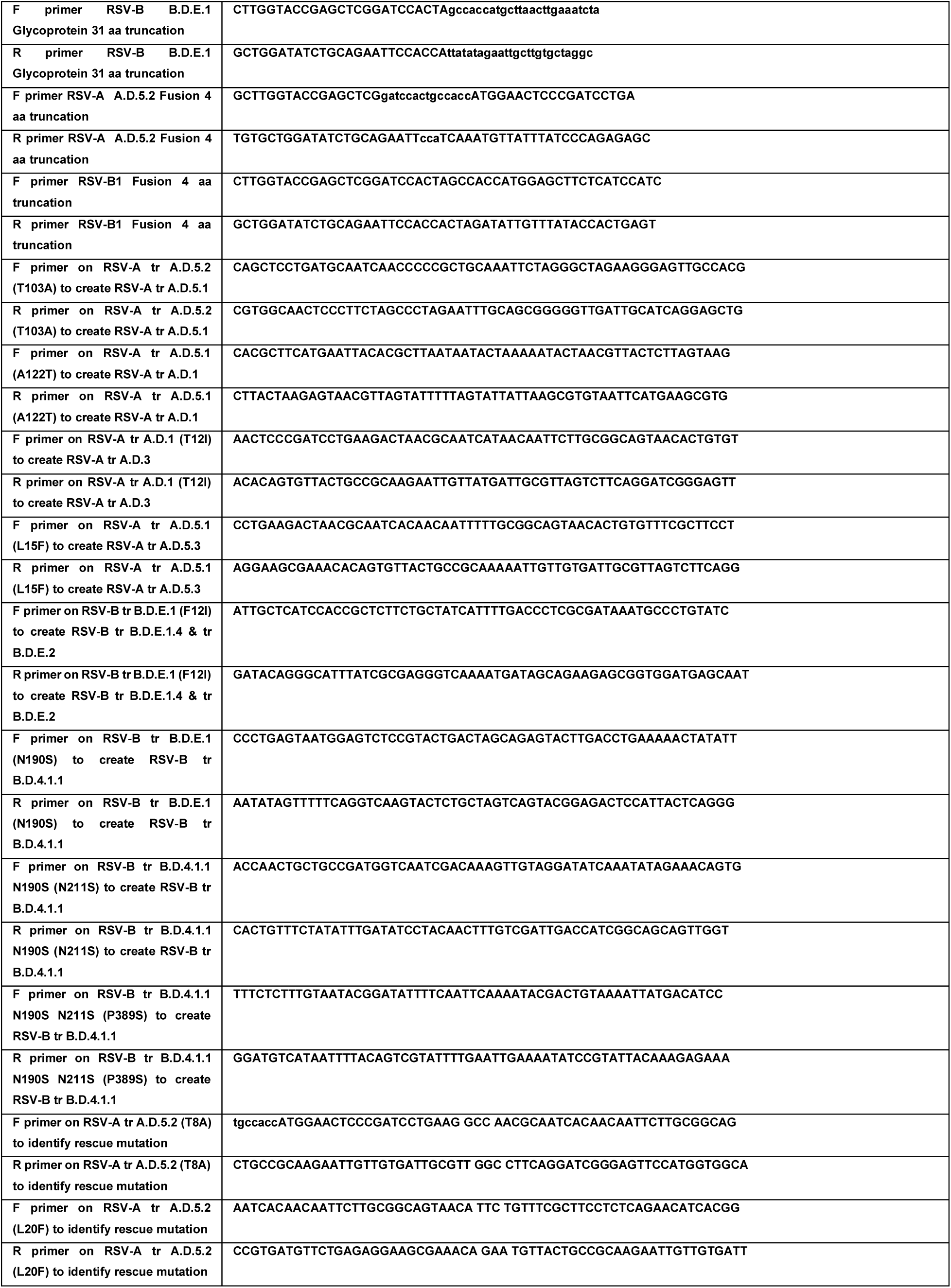
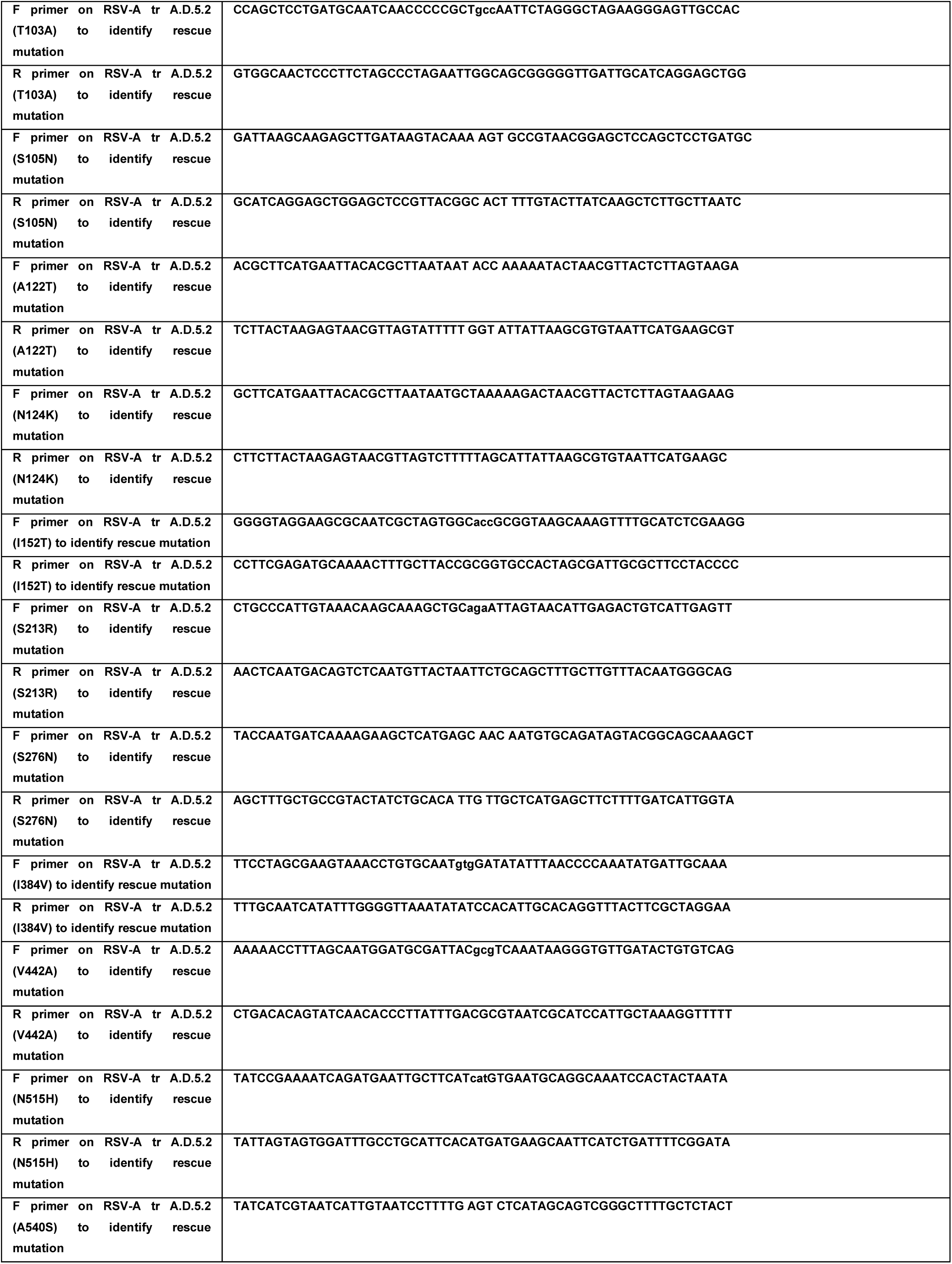
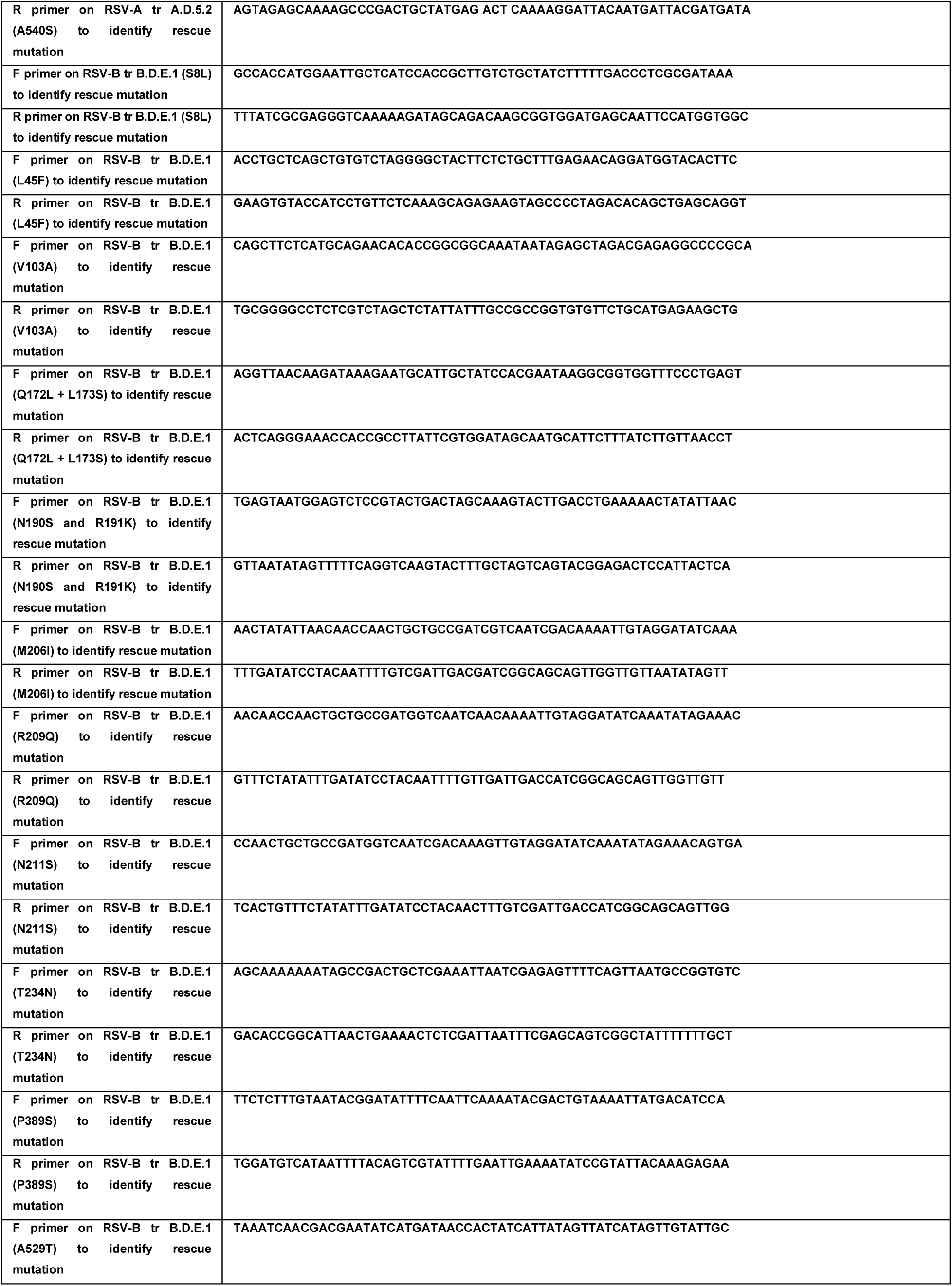
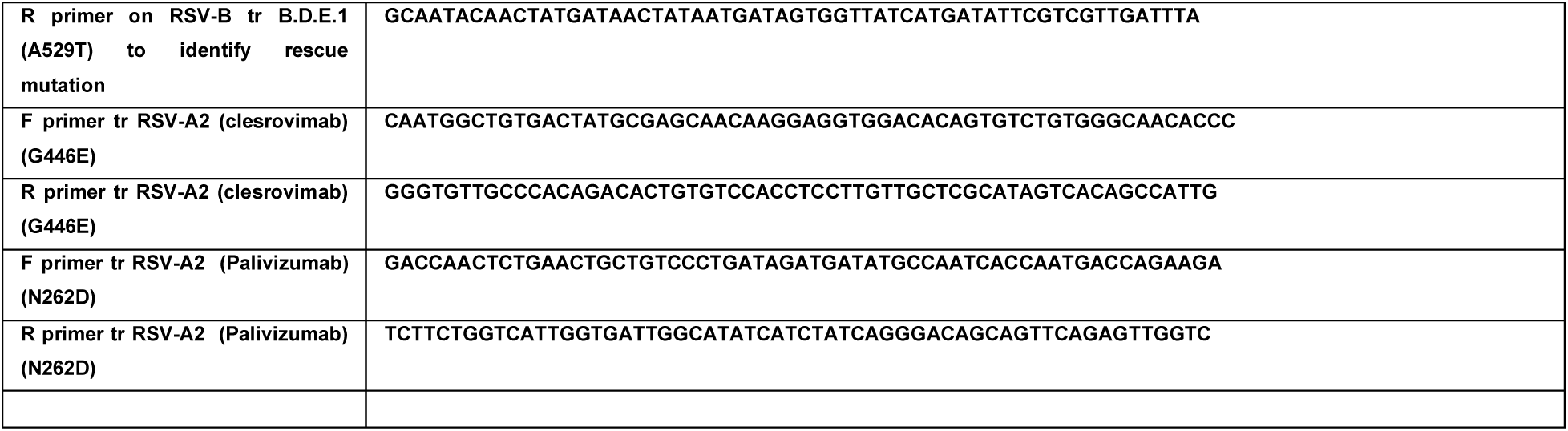
| Primer and oligo design for RSV pseudotyping and Fusion assays.

