## Supplementary Figures and Tables for "Functional Characterization of RSV Clades Associated with Prophylactic Breakthrough Infections in Pediatric Transmission Clusters"

### 1 SUPPLEMENTARY FIGURES

| Category, n (%) | **NM Cohort (n=80)** | **LCH Cohort (n=340)** |
|---|---|---|
| **Age Group at Admission, n (%)** | | |
| 0 - 2 years | 25 (31.2) | 189 (55.6) |
| 2 - 18 years | 19 (23.8) | 151 (44.6) |
| 18 - 50 years | 15 (18.8) | 0 (0.0) |
| 50-65 years | 11 (13.8) | 0 (0.0) |
| 65+ years | 10 (12.5) | 0 (0.0) |
| **Race, n (%)** | | |
| White | 57 (71.2) | 88 (25.9) |
| Black or African American | 8 (10.0) | 51 (15.0) |
| Other | 7 (8.8) | 186 (54.7) |
| Asian | 4 (5.0) | 10 (2.9) |
| Declined or Unable to Answer | 4 (5.0) | 5 (1.5) |
| **Ethnicity, n (%)** | | |
| Hispanic or Latino | 13 (16.2) | 192 (56.5) |
| Not Hispanic or Latino | 63 (78.8) | 146 (42.9) |
| Declined or Unable to Respond | 4 (5.0) | 2 (0.6) |
| **Sex, n (%)** | | |
| Female | 45 (56.2) | 167 (49.1) |
| Male | 35 (43.8) | 173 (50.9) |
| **Outcome, n (%)** | | |
| Not Hospitalized | 72 (90.0) | 288 (84.7) |
| Hospitalized, No ICU or PICU | 3 (3.8) | 39 (11.5) |
| ICU or PICU | 1 (1.2) | 11 (3.2) |
| Death | 1 (1.2) | 2 (0.6) |
| Unknown | 3 (3.8) | 0 (0.0) |
| **Comorbidities, n (%)** | | |
| Other | 20 (25.0) | 30 (8.8) |
| Cardiovascular | 14 (17.5) | 9 (2.6) |
| Chronic Lung Disease | 11 (13.8) | 14 (4.1) |

| | | |
|---|---|---|
| Immunocompromised | 2 (2.5) | 10 (2.9) |
| Prematurity | Unknown | 7 (2.0) |
| **Nirsevimab Administered, n (%)** | | |
| Ineligible | 0 (0.0) | 156 (45.9) |
| Yes | 0 (0.0) | 42 (12.4) |
| No | 0 (0.0) | 112 (32.9) |
| Administered After Infection | 0 (0.0) | 2 (0.6) |
| Unknown | 80 (100.0) | 28 (8.2) |
| **Season, n (%)** | | |
| 2023-2024 | 74 (92.5) | 0 (0.0) |
| 2024-2025 | 6 (7.5) | 340 (100.0) |
| **Subtype, n (%)** | | |
| A | 24 (30.0) | 289 (85.0) |
| B | 56 (70.0) | 51 (15.0) |

**Supplementary Table 1 | 2023-2025 RSV cohort demographics.**

Demographic and clinical features of RSV-positive patients with paired RSV whole genome sequences from NM hospital systems (n=80, left) and LCH (n=338, right). * Nirsevimab eligibility includes infants under 8 months of age encountering their first RSV season between October 1st and March 31st of each year as recommended by the American Academy of Pediatrics.

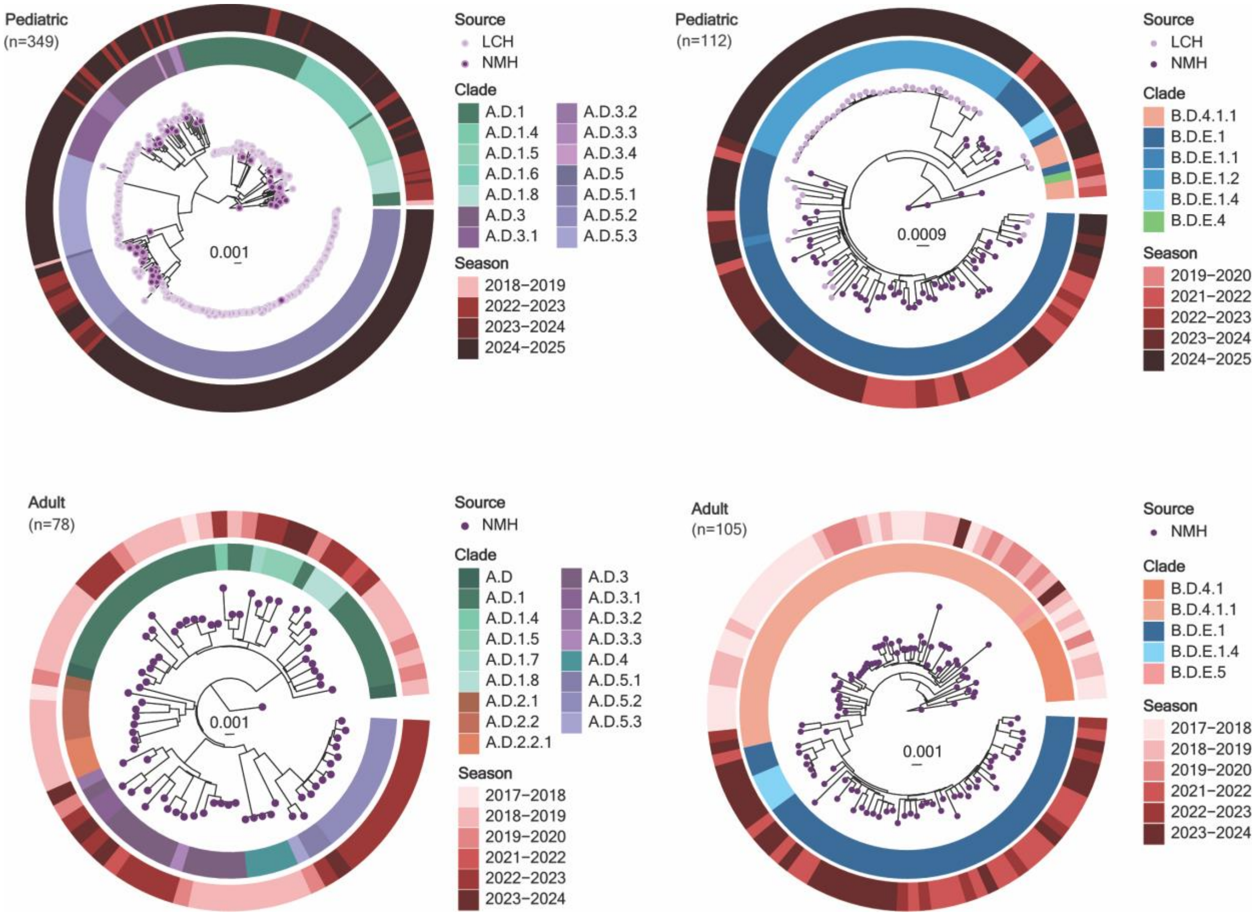

**Supplementary Figure 1 | Historical Sequencing Efforts for adult and pediatric RSV sequences.**

ML phylogenetic analysis of RSV genomes from NM and LCH in Chicago, Illinois from the 2017 to 2025 RSV seasons (n=644). For all trees, branch tips are colored by patient encounter age group, the inner ring is colored by Nextclade v3.21.0 WGS clade designation, and the outer ring is colored by RSV season in which the isolate is sampled from.

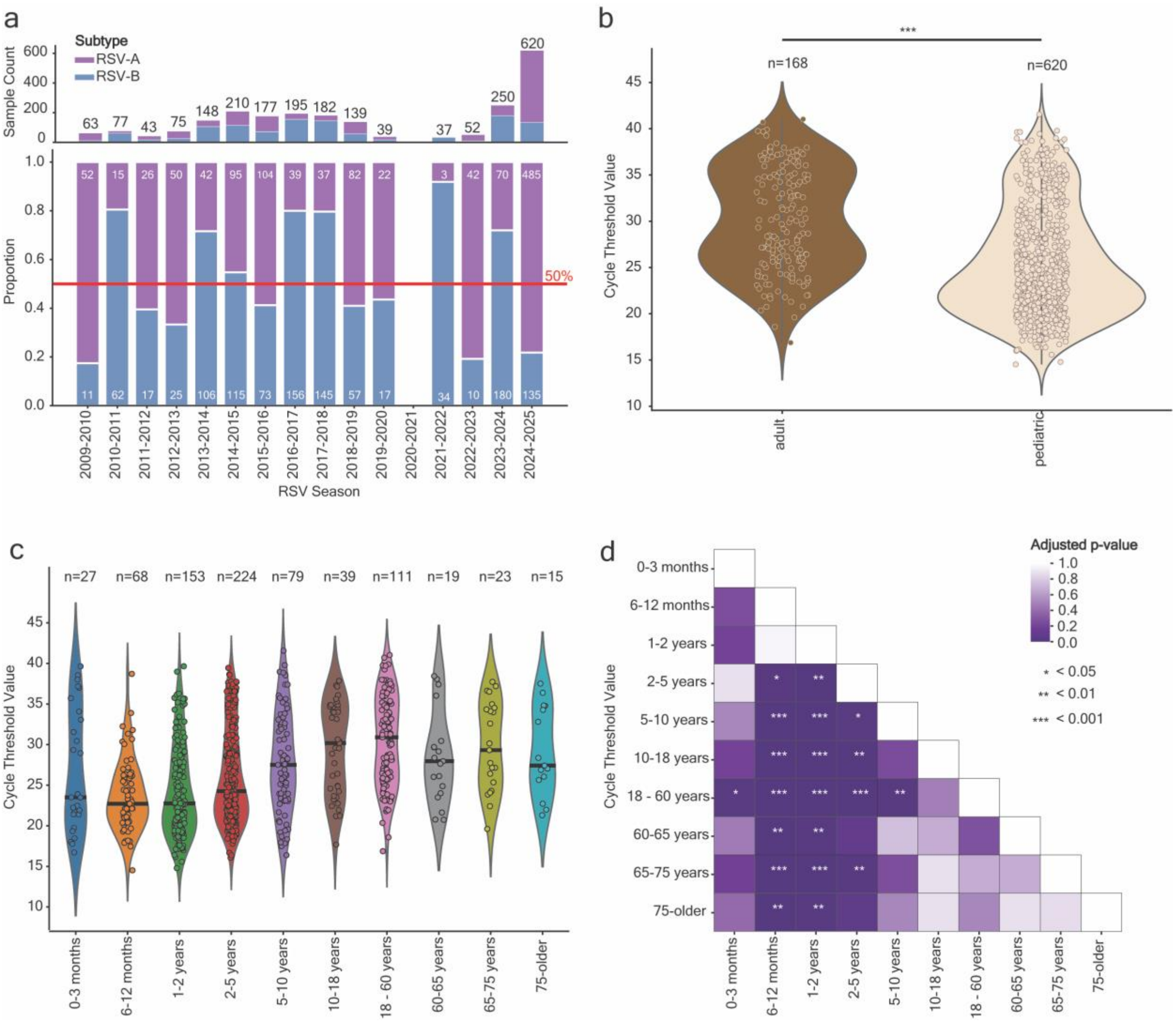

**Supplementary Figure 2 | RSV subtype distribution and viral load characteristics in adult and pediatric populations.**

**a,** Case count (top) and distribution (bottom) of RSV subtype from patient encounters in NM and LCH from April 2009 to March 2025 among 1,992 specimens with typing information [RSV-A in purple (n=828), RSV-B in blue (n=1164)]. **b,** Violin plot with scatterplot overlay of the Ct distribution among adult (brown, n=168) and pediatric (beige, n=620) RSV diagnostic specimens with available typing data. c, Violin plot with scatterplot overlay of the Ct distributions among patient encounters with available Ct data (n=788) stratified by patient age. **d,** pairwise comparison heatmap of Benjamini–Hochberg adjusted p values. Significant p values denoted by asterisks (* indicates 0.05, ** indicates <0.01, and *** indicates <0.001)

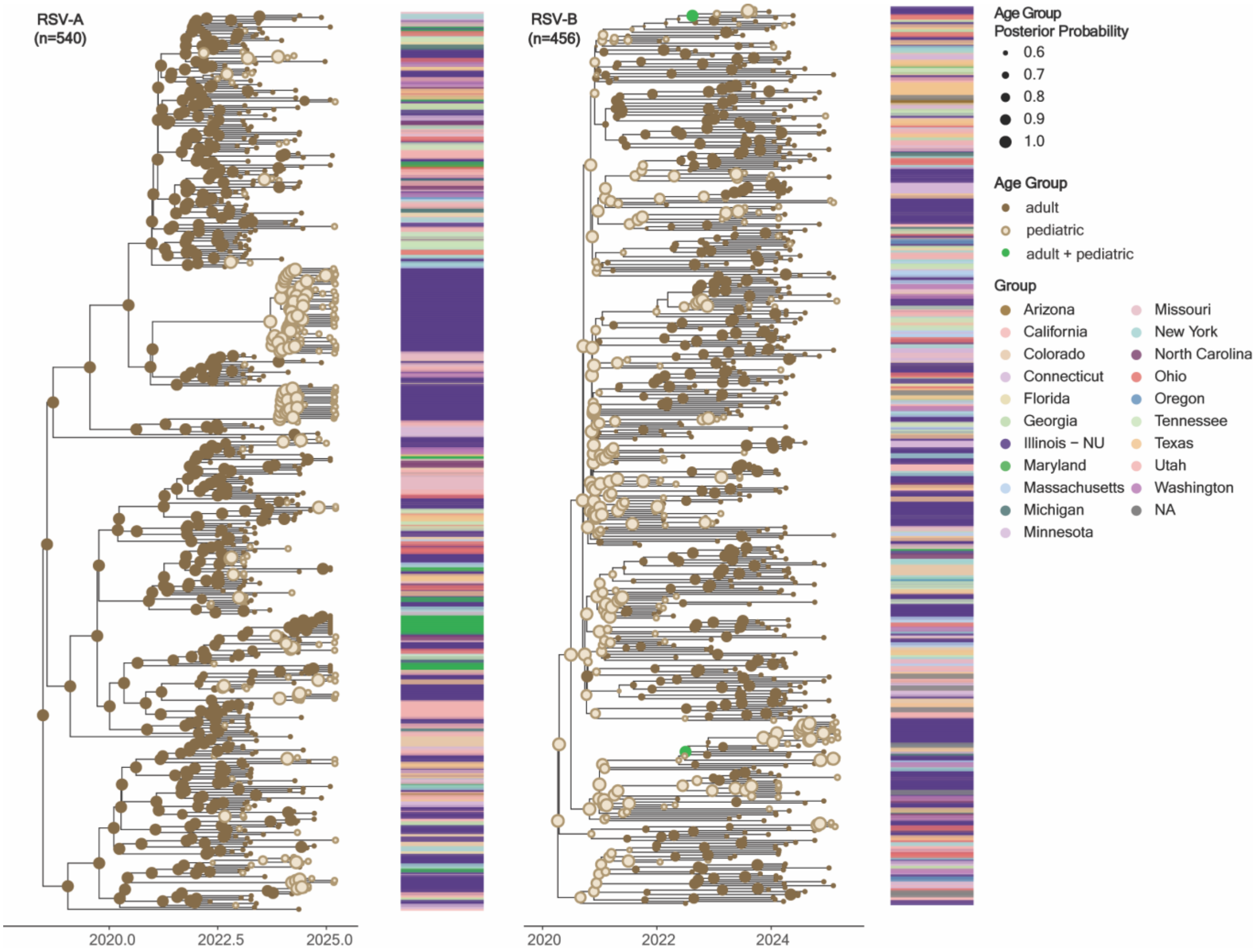

**Supplementary Figure 3 | Temporal Bayesian analysis of in-house and US-based RSV-A and RSV-B sequences by patient age group.** Bayesian phylogenetic temporal tree of circulating RSV-A (n=540) and RSV-B (n=456) genomes sampled in-house and from publicly available US genomes with known age group status. For both trees, branch tips are colored by patient age group status, and node color colored by the most probable sampled population (pediatric vs. adult). The size of the node circle represents the probability of the ancestral origin for the node of either pediatric or adult age groups. The bar adjacent to each tree is colored by sampling state.

a

| | Abrysvo (n=5) | 1st Season Nirsevimab Breakthrough (n=7) | 2nd Season Nirsevimab Breakthrough (n=35) |
|---|---|---|---|
| **Age Group at Admission, n (%)** | | | |
| 0 - 3 months | 3 (60.0) | 2 (28.6) | 0 (0.0) |
| 3 - 6 months | 1 (20.0) | 2 (28.6) | 0 (0.0) |
| 6 - 12 months | 1 (20.0) | 3 (42.9) | 10 (28.6) |
| 1-2 years | 0 (0.0) | 0 (0.0) | 25 (71.4) |
| **Race, n (%)** | | | |
| Other | 2 (40.0) | 4 (57.1) | 17 (48.6) |
| White | 1 (20.0) | 3 (42.9) | 15 (42.9) |
| Black/African American | 2 (40.0) | 0 (0.0) | 3 (8.6) |
| **Ethnicity, n (%)** | | | |
| Hispanic/Latino | 2 (40.0) | 2 (28.6) | 16 (45.7) |
| Not Hispanic or Latino | 3 (60.0) | 5 (71.4) | 19 (54.3) |
| **Sex, n (%)** | | | |
| Male | 3 (60.0) | 4 (57.1) | 16 (45.7) |
| Female | 2 (40.0) | 3 (42.9) | 19 (54.3) |
| **Outcome, n (%)** | | | |
| No Hospitalization | 2 (40.0) | 6 (85.7) | 26 (74.3) |
| Hospitalized, No PICU | 2 (40.0) | 1 (14.3) | 9 (25.7) |
| PICU | 1 (20.0) | 0 (0.0) | 0 (0.0) |
| Death | 0 (0.0) | 0 (0.0) | 0 (0.0) |
| **Comorbidities, n (%)** | | | |
| Prematurity | 0 (0.0) | 0 (0.0) | 3 (8.6) |
| Chronic Lung Condition | 0 (0.0) | 1 (14.3) | 0 (0.0) |
| Cardiovascular | 0 (0.0) | 0 (0.0) | 0 (0.0) |
| Immunocompromised | 0 (0.0) | 0 (0.0) | 0 (0.0) |
| Other | 1 (20.0) | 1 (14.3) | 3 (8.6) |
| **Season, n (%)** | | | |
| 2024-2025 | 5 (100.0) | 7 (100.0) | 35 (100.0) |
| **Subtype, n (%)** | | | |
| A | 4 (80.0) | 7 (100.0) | 31 (88.6) |
| B | 1 (20.0) | 0 (0.0) | 4 (11.4) |
| **Clade, n (%)** | | | |
| A.D.1 | 1 (20.0) | 3 (42.9) | 6 (17.1) |
| A.D.1.6 | 1 (20.0) | 0 (0.0) | 2 (5.7) |
| A.D.3 | 0 (0.0) | 0 (0.0) | 5 (14.3) |
| A.D.3.3 | 0 (0.0) | 0 (0.0) | 1 (2.9) |
| A.D.5.1 | 2 (40.0) | 3 (42.9) | 12 (34.3) |
| A.D.5.2 | 0 (0.0) | 0 (0.0) | 1 (2.9) |
| A.D.5.3 | 0 (0.0) | 1 (14.3) | 4 (11.4) |
| B.D.E.1 | 0 (0.0) | 0 (0.0) | 2 (5.7) |
| B.D.E.1.2 | 1 (20.0) | 0 (0.0) | 2 (5.7) |

b

Nirsevimab Administered vs Not
full model

| Characteristic | OR | 95% CI | p-value |
|---|---|---|---|
| Sex | | | |
| Female | — | — | |
| Male | 1.00 | 0.39, 2.59 | >0.9 |
| Race | | | |
| White | — | — | |
| Black/African American | 0.19 | 0.03, 0.80 | **0.033** |
| Other | 0.71 | 0.14, 3.66 | 0.7 |
| Ethnicity | | | |
| Not Hispanic or Latino | — | — | |
| Hispanic/Latino | 0.85 | 0.17, 4.17 | 0.8 |
| Age | 0.53 | 0.20, 1.31 | 0.2 |
| Days of Hospitalization | 0.47 | 0.06, 0.98 | 0.3 |
| Outcome | | | |
| No Hospitalization | — | — | |
| Hospitalization or PICU | 41.7 | 2.87, 15,292 | 0.077 |
| RSV Subtype | | | |
| A | — | — | |
| B | 0.56 | 0.14, 1.84 | 0.4 |
| Comorbidity Sum | 2.50 | 0.67, 9.53 | 0.2 |

Abbreviations: CI = Confidence Interval, OR = Odds Ratio

**Supplementary Figure 4 | Demographics of LCH patients with administered RSV prophylaxis and modeling patient outcome by nirsevimab administration.**

**a,** Demographic and clinical features of pediatric encounters with nirsevimab or Abrysvo-related breakthrough RSV infections. **b,** Demographic and virologic variables incorporated in a multivariable logistic regression to model nirsevimab administration among pediatric patients exposed to infection in their second RSV season (n = 116).

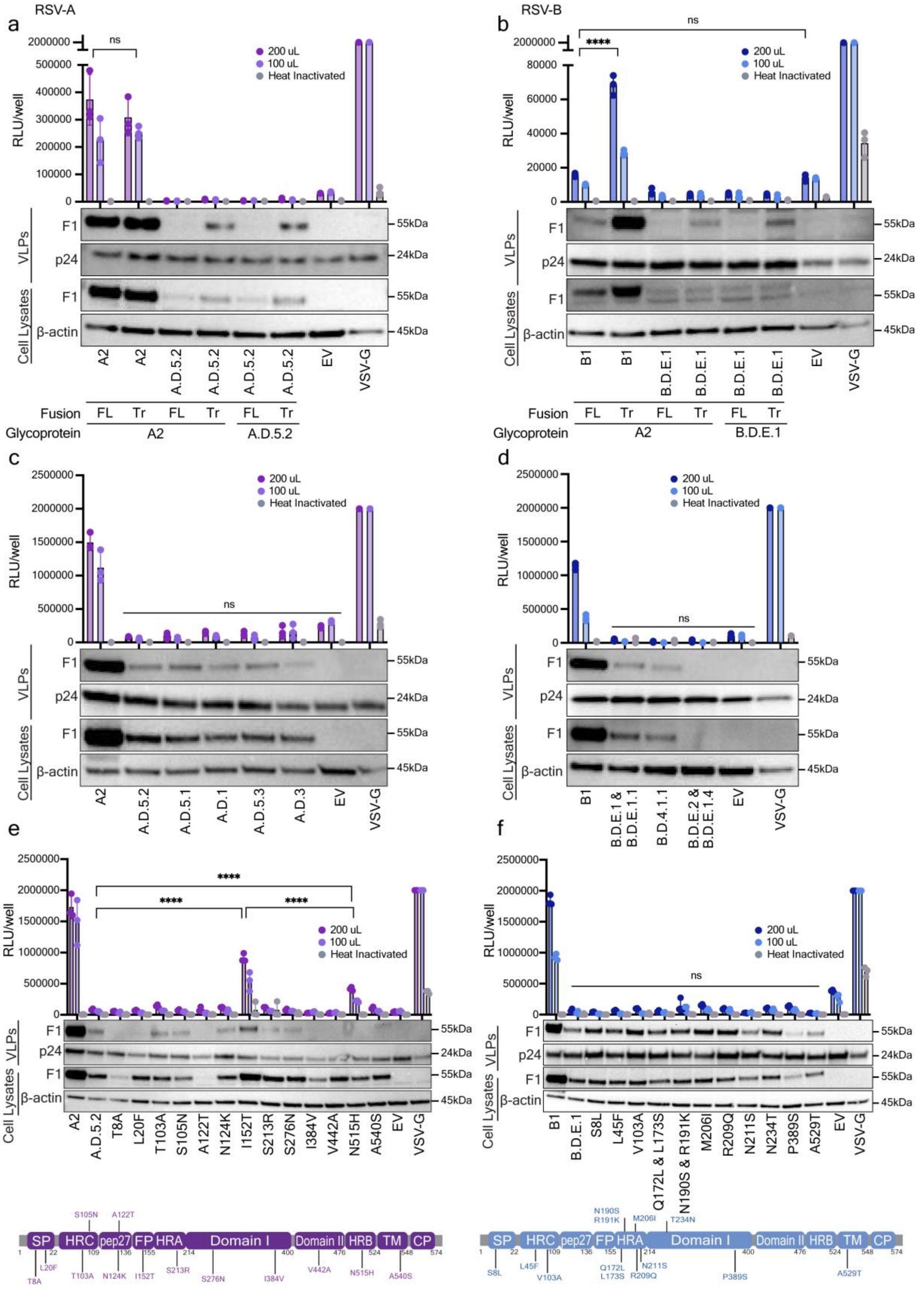
RSV-A
a
RSV-B
b
200 uL
100 uL
Heat Inactivated
RLU/well
ns
****
F1
p24
β-actin
VLPs
Cell Lysates
55kDa
24kDa
45kDa
A2
A.D.5.2
EV
VSV-G
B1
B.D.E.1
Fusion
FL
Tr
Glycoprotein
c
d
A.D.5.1
A.D.1
A.D.5.3
A.D.3
B.D.E.1 & B.D.E.1.1
B.D.4.1.1
B.D.E.2 & B.D.E.1.4
e
f
T8A
L20F
T103A
S105N
A122T
N124K
I152T
S213R
S276N
I384V
V442A
N515H
A540S
S8L
L45F
V103A
Q172L & L173S
N190S & R191K
M206I
R209Q
N211S
N234T
P389S
A529T
SP
HRC
pep27
FP
HRA
Domain I
Domain II
HRB
TM
CP

**Supplementary Figure 5 | Optimization of contemporary RSV clades in pseudotyping system.**

**a,** Lentiviral pseudoviruses bearing full-length (FL) or truncated (Tr) [A2, A.D.5.2] F and [G glycoprotein from RSV A2 with 31 AA deletion or G glycoprotein from A.D.5.2 31 AA deletion] to recover viral entry activity **b,** Lentiviral pseudoviruses bearing full-length (FL) or truncated (Tr) [B1, B.D.E.1] F and [G glycoprotein from RSV A2 with 31 AA deletion or G glycoprotein from B.D.E.1 31 AA deletion] to recover viral entry activity **c,** Truncated (Tr) pseudoviruses constructs based on A.D.5.2 backbone were engineered to contain individual A2 -specific point mutations using site-directed mutagenesis (SDM). **d**, Truncated (Tr) pseudoviruses constructs based on B.D.E1 backbone were engineered to contain individual B1 -specific point mutations using SDM. **e,** Lentiviral pseudoviruses bearing truncated [A2, A.D.5.2, A.D.5.1, A.D.1, A.D.5.3, A.D.3] F to confirm no activity in any contemporary clades. **f,** HEK293T effector cells were transiently transfected with plasmids encoding truncated [A2, A.D.5.2, A.D.5.1, A.D.1, A.D.5.3, A.D.3] F with and without mutation (I152T), as well as a T7 polymerase. Target cells were transfected with T7 promoter-driven luciferase reporter effector HEK293T cells were mixed at a 1:1 ratio with target HEK293T cells and co-cultured fusion-dependent luciferase activity was measured 48 HPI. **a-d**, all pseudoviruses were quantified and normalized to p24 levels before challenging. Viral entry was quantified by luciferase activity at 72 hours post-infection. Cell lysates (CL) and purified virus-like particles (VLPs) normalized to p24 were analyzed by immunoblotting for each RSV F. p24 was used as a VLP loading control, and β-actin as a cellular loading control. Data points represent the mean ± s.d. from three independent biological replicates. Statistics were calculated using two-way ANOVA followed by Tukey's multiple comparisons test. Pointed brackets indicate statistical comparisons between two specific conditions. Flat horizontal lines denote comparisons across all conditions. ns, not significant; $p < 0.05$ (*); $p < 0.01$ (**); $p < 0.001$ (***); $p < 0.0001$ (****).

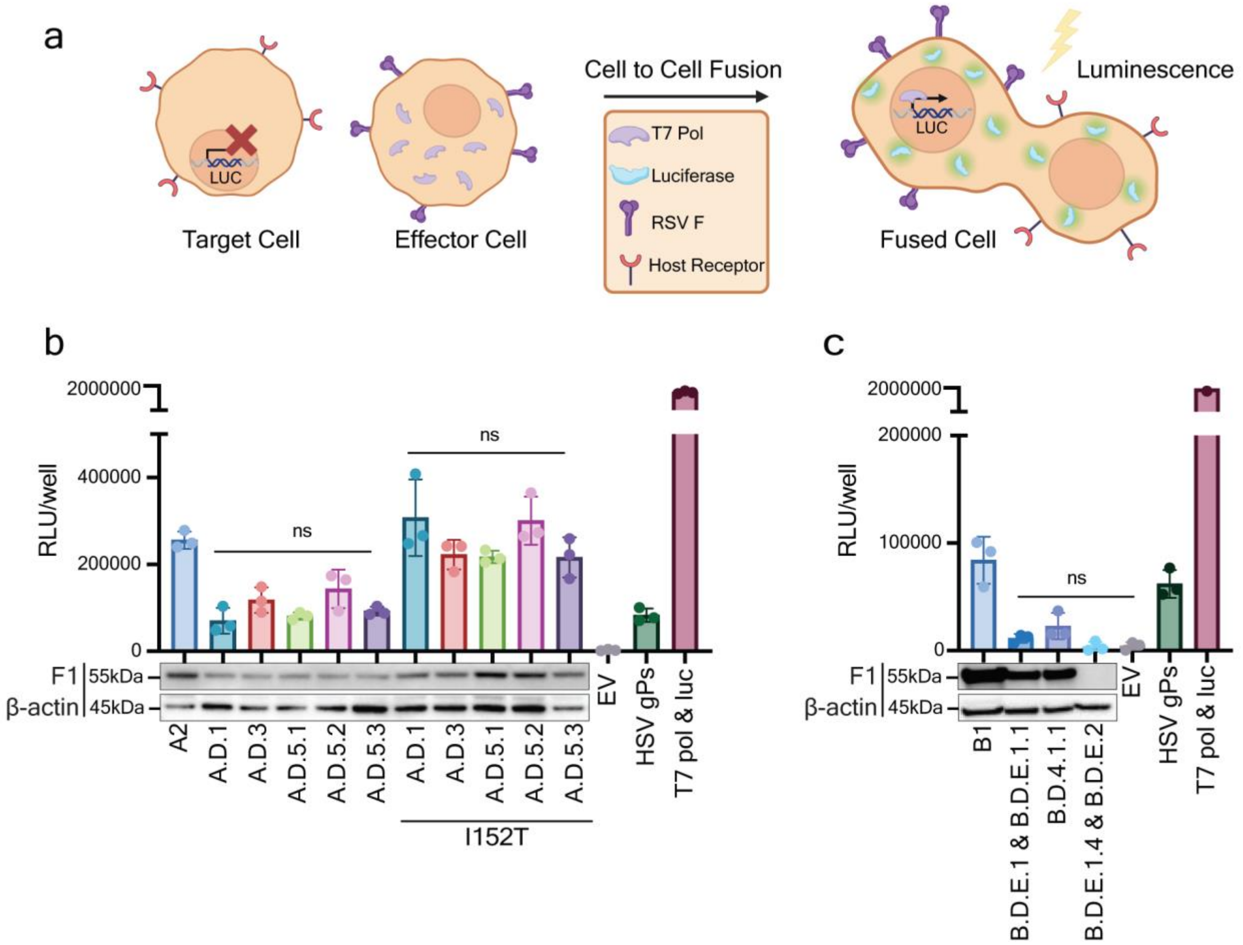

**Supplementary Figure 6 | Characterization of contemporary RSV F activity through Fusion assay.**

**a**, A schema of the RSV Fusion Assay. HEK293T target cells were transfected with T7 promoter-driven luciferase reporter. HEK293T effector cells were transiently transfected with plasmids encoding viral F glycoproteins together as well as T7 polymerase. After 24-hour transfection, HEK293T effector cells were mixed at a 1:1 ratio with HEK293T target cells and co-cultured. Fusion was quantified via luminescence 48-hour post-infection (HPI). **b**, Effector cells were transiently transfected with plasmids encoding truncated [A2, A.D.5.2, A.D.5.1, A.D.1, A.D.5.3, A.D.3] F with and without mutation (I152T), as well as T7 polymerase. Target cells were transfected with T7 promoter-driven luciferase reporter. Target and effector cells were mixed at a 1:1 ratio and co-cultured. Fusion-dependent luciferase activity was measured 48 HPI. **c**, Effector cells were transiently transfected with plasmids encoding truncated [B1, B.D.E.1 & B.D.E.1.1, B.D.4.1.1, B.D.E.1.4 & B.D.E.2] F, as well as a T7 polymerase. Target cells were transfected with T7 promoter-driven luciferase reporter. Target and effector cells were mixed at a 1:1 ratio and co-cultured. Fusion-dependent luciferase activity was measured 48 HPI. **b-c**, Cell lysates (CL) were analyzed by immunoblotting for each RSV F. β-actin was used as a cellular loading control. Data points represent the mean ± s.d. from three independent biological replicates. Statistics were calculated using one-way ANOVA followed by Tukey's multiple comparisons test. Flat horizontal lines denote comparisons across all conditions. ns, not significant; $p < 0.05$ (*); $p < 0.01$ (**); $p < 0.001$ (***); $p < 0.0001$ (****).

| seqName | Isolate | Season | Collection_date | Location | Age Group Status | Isolation_source | Genotype | NCBI Accession Number* |
|---|---|---|---|---|---|---|---|---|
| RSV566 | RSV-A/human/USA/IL-NM-RSV566/2023 | 2022-2023 | Jan-2023 | North America/USA/Illinois | adult | Nasopharyngeal | A.D.5.2 | PZ279136 |
| RSV564 | RSV-A/human/USA/IL-NM-RSV564/2023 | 2022-2023 | Jan-2023 | North America/USA/Illinois | adult | Nasopharyngeal | A.D.3.1 | PZ279135 |
| RSV573 | RSV-A/human/USA/IL-NM-RSV573/2023 | 2022-2023 | Feb-2023 | North America/USA/Illinois | adult | Nasopharyngeal | A.D.5.2 | PZ279137 |
| RSV578 | RSV-A/human/USA/IL-NM-RSV578/2023 | 2022-2023 | Feb-2023 | North America/USA/Illinois | adult | Nasopharyngeal | A.D.5.1 | PZ279138 |
| RSV579 | RSV-A/human/USA/IL-NM-RSV579/2023 | 2022-2023 | Feb-2023 | North America/USA/Illinois | adult | Nasopharyngeal | A.D.5.2 | PZ279139 |
| RSV585 | RSV-A/human/USA/IL-NM-RSV585/2023 | 2023-2024 | Jul-2023 | North America/USA/Illinois | pediatric | Nasopharyngeal | A.D.5.2 | PZ279140 |
| RSV592 | RSV-A/human/USA/IL-NM-RSV592/2023 | 2023-2024 | Aug-2023 | North America/USA/Illinois | pediatric | Nasopharyngeal | A.D.1.6 | PZ279142 |
| RSV591 | RSV-A/human/USA/IL-NM-RSV591/2023 | 2023-2024 | Sep-2023 | North America/USA/Illinois | adult | Nasopharyngeal | A.D.3 | PZ279141 |
| RSV608 | RSV-A/human/USA/IL-NM-RSV608/2023 | 2023-2024 | Oct-2023 | North America/USA/Illinois | pediatric | Nasopharyngeal | A.D.1.5 | PZ279143 |
| RSV611 | RSV-A/human/USA/IL-NM-RSV611/2023 | 2023-2024 | Oct-2023 | North America/USA/Illinois | pediatric | Nasopharyngeal | A.D.5.2 | PZ279144 |
| RSV615 | RSV-A/human/USA/IL-NM-RSV615/2023 | 2023-2024 | Oct-2023 | North America/USA/Illinois | adult | Nasopharyngeal | A.D.5.1 | PZ279145 |
| RSV619 | RSV-A/human/USA/IL-NM-RSV619/2023 | 2023-2024 | Oct-2023 | North America/USA/Illinois | pediatric | Nasopharyngeal | A.D.1.8 | PZ279146 |
| RSV623 | RSV-A/human/USA/IL-NM-RSV623/2023 | 2023-2024 | Oct-2023 | North America/USA/Illinois | pediatric | Nasopharyngeal | A.D.1 | PZ279147 |
| RSV636 | RSV-A/human/USA/IL-NM-RSV636/2023 | 2023-2024 | Nov-2023 | North America/USA/Illinois | pediatric | Nasopharyngeal | A.D.5.2 | |
| RSV637 | RSV-A/human/USA/IL-NM-RSV637/2023 | 2023-2024 | Nov-2023 | North America/USA/Illinois | pediatric | Nasopharyngeal | A.D.5.2 | PZ279148 |
| RSV651 | RSV-A/human/USA/IL-NM-RSV651/2023 | 2023-2024 | Nov-2023 | North America/USA/Illinois | adult | Nasopharyngeal | A.D.3.1 | |
| RSV658 | RSV-A/human/USA/IL-NM-RSV658/2023 | 2023-2024 | Nov-2023 | North America/USA/Illinois | pediatric | Nasopharyngeal | A.D.1.8 | PZ279149 |
| RSV661 | RSV-A/human/USA/IL-NM-RSV661/2023 | 2023-2024 | Nov-2023 | North America/USA/Illinois | adult | Nasopharyngeal | A.D.3.2 | PZ279150 |
| RSV668 | RSV-A/human/USA/IL-NM-RSV668/2023 | 2023-2024 | Nov-2023 | North America/USA/Illinois | adult | Nasopharyngeal | A.D.1.5 | PZ279151 |
| RSV684 | RSV-A/human/USA/IL-NM-RSV684/2023 | 2023-2024 | Dec-2023 | North America/USA/Illinois | pediatric | Nasopharyngeal | A.D.5.2 | PZ279152 |
| RSV710 | RSV-A/human/USA/IL-NM-RSV710/2023 | 2023-2024 | Dec-2023 | North America/USA/Illinois | adult | Nasopharyngeal | A.D.1.5 | PZ279153 |
| RSV712 | RSV-A/human/USA/IL-NM-RSV712/2023 | 2023-2024 | Dec-2023 | North America/USA/Illinois | pediatric | Nasopharyngeal | A.D.5.2 | |
| RSV769 | RSV-A/human/USA/IL-NM-RSV769/2023 | 2023-2024 | Dec-2023 | North America/USA/Illinois | pediatric | Nasopharyngeal | A.D.3.2 | PZ279155 |
| RSV749 | RSV-A/human/USA/IL-NM-RSV749/2023 | 2023-2024 | Dec-2023 | North America/USA/Illinois | pediatric | Nasopharyngeal | A.D.5.2 | PZ279154 |
| RSV832 | RSV-A/human/USA/IL-NM-RSV832/2024 | 2024-2025 | Oct-2024 | North America/USA/Illinois | pediatric | Nasopharyngeal | A.D.5.1 | PZ279220 |
| LCH12826 | RSV-A/human/USA/IL-LCH-LCH12826/2024 | 2024-2025 | Nov-2024 | North America/USA/Illinois | pediatric | Nasopharyngeal | A.D.5.1 | PZ279159 |
| LCH12829 | RSV-A/human/USA/IL-LCH-LCH12829/2024 | 2024-2025 | Nov-2024 | North America/USA/Illinois | pediatric | Nasopharyngeal | A.D.5.3 | PZ279162 |
| LCH12827 | RSV-A/human/USA/IL-LCH-LCH12827/2024 | 2024-2025 | Nov-2024 | North America/USA/Illinois | pediatric | Nasopharyngeal | A.D.5.1 | PZ279160 |

| | | | | | | | | |
|---|---|---|---|---|---|---|---|---|
| LCH12828 | RSV-A/human/USA/IL-LCH-LCH12828/2024 | 2024-2025 | Nov-2024 | North America/USA/Illinois | pediatric | Nasopharyngeal | A.D.3 | PZ279161 |
| LCH12821 | RSV-A/human/USA/IL-LCH-LCH12821/2024 | 2024-2025 | Nov-2024 | North America/USA/Illinois | pediatric | Nasopharyngeal | A.D.5.3 | PZ279156 |
| LCH12823 | RSV-A/human/USA/IL-LCH-LCH12823/2024 | 2024-2025 | Nov-2024 | North America/USA/Illinois | pediatric | Nasopharyngeal | A.D.5.3 | PZ279157 |
| LCH12825 | RSV-A/human/USA/IL-LCH-LCH12825/2024 | 2024-2025 | Nov-2024 | North America/USA/Illinois | pediatric | Nasopharyngeal | A.D.5.1 | PZ279158 |
| LCH12849 | RSV-A/human/USA/IL-LCH-LCH12849/2024 | 2024-2025 | Nov-2024 | North America/USA/Illinois | pediatric | Nasopharyngeal | A.D.5.1 | PZ279168 |
| LCH12850 | RSV-A/human/USA/IL-LCH-LCH12850/2024 | 2024-2025 | Nov-2024 | North America/USA/Illinois | pediatric | Nasopharyngeal | A.D.5.1 | PZ279169 |
| LCH12844 | RSV-A/human/USA/IL-LCH-LCH12844/2024 | 2024-2025 | Nov-2024 | North America/USA/Illinois | pediatric | Nasopharyngeal | A.D.5.3 | PZ279166 |
| LCH12851 | RSV-A/human/USA/IL-LCH-LCH12851/2024 | 2024-2025 | Nov-2024 | North America/USA/Illinois | pediatric | Nasopharyngeal | A.D.5.3 | PZ279170 |
| LCH12846 | RSV-A/human/USA/IL-LCH-LCH12846/2024 | 2024-2025 | Nov-2024 | North America/USA/Illinois | pediatric | Nasopharyngeal | A.D.5.1 | PZ279167 |
| LCH12837 | RSV-A/human/USA/IL-LCH-LCH12837/2024 | 2024-2025 | Nov-2024 | North America/USA/Illinois | pediatric | Nasopharyngeal | A.D.5.3 | PZ279164 |
| LCH12843 | RSV-A/human/USA/IL-LCH-LCH12843/2024 | 2024-2025 | Nov-2024 | North America/USA/Illinois | pediatric | Nasopharyngeal | A.D.5.1 | PZ279165 |
| LCH12831 | RSV-A/human/USA/IL-LCH-LCH12831/2024 | 2024-2025 | Nov-2024 | North America/USA/Illinois | pediatric | Nasopharyngeal | A.D.1 | PZ279163 |
| LCH12858 | RSV-A/human/USA/IL-LCH-LCH12858/2024 | 2024-2025 | Nov-2024 | North America/USA/Illinois | pediatric | Nasopharyngeal | A.D.5.2 | PZ279173 |
| LCH12861 | RSV-A/human/USA/IL-LCH-LCH12861/2024 | 2024-2025 | Nov-2024 | North America/USA/Illinois | pediatric | Nasopharyngeal | A.D.3 | PZ279176 |
| LCH12867 | RSV-A/human/USA/IL-LCH-LCH12867/2024 | 2024-2025 | Nov-2024 | North America/USA/Illinois | pediatric | Nasopharyngeal | A.D.5.1 | PZ279178 |
| LCH12868 | RSV-A/human/USA/IL-LCH-LCH12868/2024 | 2024-2025 | Nov-2024 | North America/USA/Illinois | pediatric | Nasopharyngeal | A.D.5.1 | PZ279179 |
| RSV845 | RSV-A/human/USA/-NM-RSV845/2024 | 2024-2025 | Nov-2024 | North America/USA/Illinois | pediatric | Nasopharyngeal | A.D.1.6 | PZ279221 |
| LCH12854 | RSV-A/human/USA/IL-LCH-LCH12854/2024 | 2024-2025 | Nov-2024 | North America/USA/Illinois | pediatric | Nasopharyngeal | A.D.5.1 | PZ279171 |
| LCH12855 | RSV-A/human/USA/IL-LCH-LCH12855/2024 | 2024-2025 | Nov-2024 | North America/USA/Illinois | pediatric | Nasopharyngeal | A.D.5.3 | PZ279172 |
| LCH12859 | RSV-A/human/USA/IL-LCH-LCH12859/2024 | 2024-2025 | Nov-2024 | North America/USA/Illinois | pediatric | Nasopharyngeal | A.D.1 | PZ279174 |
| LCH12863 | RSV-A/human/USA/IL-LCH-LCH12863/2024 | 2024-2025 | Nov-2024 | North America/USA/Illinois | pediatric | Nasopharyngeal | A.D.5.3 | PZ279177 |
| LCH12872 | RSV-A/human/USA/IL-LCH-LCH12872/2024 | 2024-2025 | Nov-2024 | North America/USA/Illinois | pediatric | Nasopharyngeal | A.D.1.6 | PZ279180 |
| LCH12860 | RSV-A/human/USA/IL-LCH-LCH12860/2024 | 2024-2025 | Nov-2024 | North America/USA/Illinois | pediatric | Nasopharyngeal | A.D.5.1 | PZ279175 |
| LCH12874 | RSV-A/human/USA/IL-LCH-LCH12874/2024 | 2024-2025 | Nov-2024 | North America/USA/Illinois | pediatric | Nasopharyngeal | A.D.3 | PZ279181 |
| LCH12892 | RSV-A/human/USA/IL-LCH-LCH12892/2024 | 2024-2025 | Nov-2024 | North America/USA/Illinois | pediatric | Nasopharyngeal | A.D.3 | PZ279191 |
| LCH12876 | RSV-A/human/USA/IL-LCH-LCH12876/2024 | 2024-2025 | Nov-2024 | North America/USA/Illinois | pediatric | Nasopharyngeal | A.D.5.3 | PZ279183 |
| LCH12878 | RSV-A/human/USA/IL-LCH-LCH12878/2024 | 2024-2025 | Nov-2024 | North America/USA/Illinois | pediatric | Nasopharyngeal | A.D.1 | PZ279185 |
| LCH12879 | RSV-A/human/USA/IL-LCH-LCH12879/2024 | 2024-2025 | Nov-2024 | North America/USA/Illinois | pediatric | Nasopharyngeal | A.D.5.2 | PZ279186 |
| LCH12893 | RSV-A/human/USA/IL-LCH-LCH12893/2024 | 2024-2025 | Nov-2024 | North America/USA/Illinois | pediatric | Nasopharyngeal | A.D.5.3 | PZ279192 |
| LCH12875 | RSV-A/human/USA/IL-LCH-LCH12875/2024 | 2024-2025 | Nov-2024 | North America/USA/Illinois | pediatric | Nasopharyngeal | A.D.5.1 | PZ279182 |

| | | | | | | | | |
|---|---|---|---|---|---|---|---|---|
| LCH12877 | RSV-A/human/USA/IL-LCH-LCH12877/2024 | 2024-2025 | Nov-2024 | North America/USA/Illinois | pediatric | Nasopharyngeal | A.D.5.1 | PZ279184 |
| LCH12882 | RSV-A/human/USA/IL-LCH-LCH12882/2024 | 2024-2025 | Nov-2024 | North America/USA/Illinois | pediatric | Nasopharyngeal | A.D.5.3 | PZ279188 |
| LCH12881 | RSV-A/human/USA/IL-LCH-LCH12881/2024 | 2024-2025 | Nov-2024 | North America/USA/Illinois | pediatric | Nasopharyngeal | A.D.5.3 | PZ279187 |
| LCH12887 | RSV-A/human/USA/IL-LCH-LCH12887/2024 | 2024-2025 | Nov-2024 | North America/USA/Illinois | pediatric | Nasopharyngeal | A.D.5.1 | PZ279189 |
| LCH12891 | RSV-A/human/USA/IL-LCH-LCH12891/2024 | 2024-2025 | Nov-2024 | North America/USA/Illinois | pediatric | Nasopharyngeal | A.D.1 | PZ279190 |
| LCH12991 | RSV-A/human/USA/IL-LCH-LCH12991/2024 | 2024-2025 | Nov-2024 | North America/USA/Illinois | pediatric | Nasopharyngeal | A.D.3 | PZ279351 |
| LCH12922 | RSV-A/human/USA/IL-LCH-LCH12922/2024 | 2024-2025 | Nov-2024 | North America/USA/Illinois | pediatric | Nasopharyngeal | A.D.5.1 | PZ279194 |
| LCH12947 | RSV-A/human/USA/IL-LCH-LCH12947/2024 | 2024-2025 | Nov-2024 | North America/USA/Illinois | pediatric | Nasopharyngeal | A.D.5.1 | PZ279199 |
| LCH12961 | RSV-A/human/USA/IL-LCH-LCH12961/2024 | 2024-2025 | Nov-2024 | North America/USA/Illinois | pediatric | Nasopharyngeal | A.D.5.1 | PZ279203 |
| LCH12964 | RSV-A/human/USA/IL-LCH-LCH12964/2024 | 2024-2025 | Nov-2024 | North America/USA/Illinois | pediatric | Nasopharyngeal | A.D.1.5 | PZ279204 |
| LCH12971 | RSV-A/human/USA/IL-LCH-LCH12971/2024 | 2024-2025 | Nov-2024 | North America/USA/Illinois | pediatric | Nasopharyngeal | A.D.5.3 | PZ279210 |
| LCH12996 | RSV-A/human/USA/IL-LCH-LCH12996/2024 | 2024-2025 | Nov-2024 | North America/USA/Illinois | pediatric | Nasopharyngeal | A.D.1.5 | PZ279259 |
| LCH12950 | RSV-A/human/USA/IL-LCH-LCH12950/2024 | 2024-2025 | Nov-2024 | North America/USA/Illinois | pediatric | Nasopharyngeal | A.D.5.1 | PZ279200 |
| LCH12959 | RSV-A/human/USA/IL-LCH-LCH12959/2024 | 2024-2025 | Nov-2024 | North America/USA/Illinois | pediatric | Nasopharyngeal | A.D.1 | PZ279202 |
| LCH12966 | RSV-A/human/USA/IL-LCH-LCH12966/2024 | 2024-2025 | Nov-2024 | North America/USA/Illinois | pediatric | Nasopharyngeal | A.D.5.1 | PZ279205 |
| LCH12978 | RSV-A/human/USA/IL-LCH-LCH12978/2024 | 2024-2025 | Nov-2024 | North America/USA/Illinois | pediatric | Nasopharyngeal | A.D.5.3 | |
| LCH12984 | RSV-A/human/USA/IL-LCH-LCH12984/2024 | 2024-2025 | Nov-2024 | North America/USA/Illinois | pediatric | Nasopharyngeal | A.D.5.1 | PZ279214 |
| LCH13003 | RSV-A/human/USA/IL-LCH-LCH13003/2024 | 2024-2025 | Nov-2024 | North America/USA/Illinois | pediatric | Nasopharyngeal | A.D.1.6 | PZ279352 |
| LCH12914 | RSV-A/human/USA/IL-LCH-LCH12914/2024 | 2024-2025 | Dec-2024 | North America/USA/Illinois | pediatric | Nasopharyngeal | A.D.5.3 | PZ279348 |
| LCH12943 | RSV-A/human/USA/IL-LCH-LCH12943/2024 | 2024-2025 | Dec-2024 | North America/USA/Illinois | pediatric | Nasopharyngeal | A.D.5.1 | |
| LCH12944 | RSV-A/human/USA/IL-LCH-LCH12944/2024 | 2024-2025 | Dec-2024 | North America/USA/Illinois | pediatric | Nasopharyngeal | A.D.5.1 | PZ279198 |
| LCH12946 | RSV-A/human/USA/IL-LCH-LCH12946/2024 | 2024-2025 | Dec-2024 | North America/USA/Illinois | pediatric | Nasopharyngeal | A.D.1.6 | PZ279349 |
| LCH12948 | RSV-A/human/USA/IL-LCH-LCH12948/2024 | 2024-2025 | Dec-2024 | North America/USA/Illinois | pediatric | Nasopharyngeal | A.D.1.6 | PZ279350 |
| LCH12967 | RSV-A/human/USA/IL-LCH-LCH12967/2024 | 2024-2025 | Dec-2024 | North America/USA/Illinois | pediatric | Nasopharyngeal | A.D.5.1 | PZ279206 |
| LCH12979 | RSV-A/human/USA/IL-LCH-LCH12979/2024 | 2024-2025 | Dec-2024 | North America/USA/Illinois | pediatric | Nasopharyngeal | A.D.3 | PZ279212 |
| LCH12990 | RSV-A/human/USA/IL-LCH-LCH12990/2024 | 2024-2025 | Dec-2024 | North America/USA/Illinois | pediatric | Nasopharyngeal | A.D.3 | PZ279216 |
| LCH13004 | RSV-A/human/USA/IL-LCH-LCH13004/2024 | 2024-2025 | Dec-2024 | North America/USA/Illinois | pediatric | Nasopharyngeal | A.D.5.3 | PZ279219 |
| LCH12908 | RSV-A/human/USA/IL-LCH-LCH12908/2024 | 2024-2025 | Dec-2024 | North America/USA/Illinois | pediatric | Nasopharyngeal | A.D.5.1 | PZ279347 |
| LCH12920 | RSV-A/human/USA/IL-LCH-LCH12920/2024 | 2024-2025 | Dec-2024 | North America/USA/Illinois | pediatric | Nasopharyngeal | A.D.3 | PZ279222 |
| LCH12932 | RSV-A/human/USA/IL-LCH-LCH12932/2024 | 2024-2025 | Dec-2024 | North America/USA/Illinois | pediatric | Nasopharyngeal | A.D.5.3 | PZ279197 |

| | | | | | | | | |
|---|---|---|---|---|---|---|---|---|
| LCH12969 | RSV-A/human/USA/IL-LCH-LCH12969/2024 | 2024-2025 | Dec-2024 | North America/USA/Illinois | pediatric | Nasopharyngeal | A.D.1 | PZ279208 |
| LCH12972 | RSV-A/human/USA/IL-LCH-LCH12972/2024 | 2024-2025 | Dec-2024 | North America/USA/Illinois | pediatric | Nasopharyngeal | A.D.3.2 | PZ279211 |
| LCH12983 | RSV-A/human/USA/IL-LCH-LCH12983/2024 | 2024-2025 | Dec-2024 | North America/USA/Illinois | pediatric | Nasopharyngeal | A.D.5.1 | PZ279213 |
| LCH12989 | RSV-A/human/USA/IL-LCH-LCH12989/2024 | 2024-2025 | Dec-2024 | North America/USA/Illinois | pediatric | Nasopharyngeal | A.D.5.1 | PZ279215 |
| LCH13001 | RSV-A/human/USA/IL-LCH-LCH13001/2024 | 2024-2025 | Dec-2024 | North America/USA/Illinois | pediatric | Nasopharyngeal | A.D.5.1 | PZ279218 |
| LCH12918 | RSV-A/human/USA/IL-LCH-LCH12918/2024 | 2024-2025 | Dec-2024 | North America/USA/Illinois | pediatric | Nasopharyngeal | A.D.5.3 | PZ279193 |
| LCH12925 | RSV-A/human/USA/IL-LCH-LCH12925/2024 | 2024-2025 | Dec-2024 | North America/USA/Illinois | pediatric | Nasopharyngeal | A.D.5.1 | PZ279195 |
| LCH12928 | RSV-A/human/USA/IL-LCH-LCH12928/2024 | 2024-2025 | Dec-2024 | North America/USA/Illinois | pediatric | Nasopharyngeal | A.D.5.1 | |
| LCH12931 | RSV-A/human/USA/IL-LCH-LCH12931/2024 | 2024-2025 | Dec-2024 | North America/USA/Illinois | pediatric | Nasopharyngeal | A.D.1.5 | PZ279196 |
| LCH12945 | RSV-A/human/USA/IL-LCH-LCH12945/2024 | 2024-2025 | Dec-2024 | North America/USA/Illinois | pediatric | Nasopharyngeal | A.D.5.1 | |
| LCH12952 | RSV-A/human/USA/IL-LCH-LCH12952/2024 | 2024-2025 | Dec-2024 | North America/USA/Illinois | pediatric | Nasopharyngeal | A.D.5.1 | PZ279201 |
| LCH12970 | RSV-A/human/USA/IL-LCH-LCH12970/2024 | 2024-2025 | Dec-2024 | North America/USA/Illinois | pediatric | Nasopharyngeal | A.D.3.1 | PZ279209 |
| LCH12968 | RSV-A/human/USA/IL-LCH-LCH12968/2024 | 2024-2025 | Dec-2024 | North America/USA/Illinois | pediatric | Nasopharyngeal | A.D.3.1 | PZ279207 |
| LCH12995 | RSV-A/human/USA/IL-LCH-LCH12995/2024 | 2024-2025 | Dec-2024 | North America/USA/Illinois | pediatric | Nasopharyngeal | A.D.3.2 | PZ279217 |
| LCH13014 | RSV-A/human/USA/IL-LCH-LCH13014/2024 | 2024-2025 | Dec-2024 | North America/USA/Illinois | pediatric | Nasopharyngeal | A.D.5.1 | PZ279262 |
| LCH13054 | RSV-A/human/USA/IL-LCH-LCH13054/2024 | 2024-2025 | Dec-2024 | North America/USA/Illinois | pediatric | Nasopharyngeal | A.D.1 | PZ279355 |
| LCH13056 | RSV-A/human/USA/IL-LCH-LCH13056/2024 | 2024-2025 | Dec-2024 | North America/USA/Illinois | pediatric | Nasopharyngeal | A.D.3.1 | PZ279356 |
| LCH13091 | RSV-A/human/USA/IL-LCH-LCH13091/2024 | 2024-2025 | Dec-2024 | North America/USA/Illinois | pediatric | Nasopharyngeal | A.D.5.1 | PZ279362 |
| LCH13161 | RSV-A/human/USA/IL-LCH-LCH13161/2024 | 2024-2025 | Dec-2024 | North America/USA/Illinois | pediatric | Nasopharyngeal | A.D.3 | PZ279371 |
| LCH13037 | RSV-A/human/USA/IL-LCH-LCH13037/2024 | 2024-2025 | Dec-2024 | North America/USA/Illinois | pediatric | Nasopharyngeal | A.D.5.1 | PZ279224 |
| LCH13063 | RSV-A/human/USA/IL-LCH-LCH13063/2024 | 2024-2025 | Dec-2024 | North America/USA/Illinois | pediatric | Nasopharyngeal | A.D.1.6 | PZ279357 |
| LCH13067 | RSV-A/human/USA/IL-LCH-LCH13067/2024 | 2024-2025 | Dec-2024 | North America/USA/Illinois | pediatric | Nasopharyngeal | A.D.5.1 | PZ279358 |
| LCH13078 | RSV-A/human/USA/IL-LCH-LCH13078/2024 | 2024-2025 | Dec-2024 | North America/USA/Illinois | pediatric | Nasopharyngeal | A.D.5.1 | PZ279359 |
| LCH13107 | RSV-A/human/USA/IL-LCH-LCH13107/2024 | 2024-2025 | Dec-2024 | North America/USA/Illinois | pediatric | Nasopharyngeal | A.D.1.5 | PZ279364 |
| LCH13125 | RSV-A/human/USA/IL-LCH-LCH13125/2024 | 2024-2025 | Dec-2024 | North America/USA/Illinois | pediatric | Nasopharyngeal | A.D.5.1 | PZ279368 |
| LCH13084 | RSV-A/human/USA/IL-LCH-LCH13084/2024 | 2024-2025 | Dec-2024 | North America/USA/Illinois | pediatric | Nasopharyngeal | A.D.5.1 | PZ279360 |
| LCH13099 | RSV-A/human/USA/IL-LCH-LCH13099/2024 | 2024-2025 | Dec-2024 | North America/USA/Illinois | pediatric | Nasopharyngeal | A.D.5.1 | PZ279363 |
| LCH13012 | RSV-A/human/USA/IL-LCH-LCH13012/2024 | 2024-2025 | Dec-2024 | North America/USA/Illinois | pediatric | Nasopharyngeal | A.D.1.5 | PZ279261 |
| LCH13118 | RSV-A/human/USA/IL-LCH-LCH13118/2024 | 2024-2025 | Dec-2024 | North America/USA/Illinois | pediatric | Nasopharyngeal | A.D.1 | PZ279366 |
| LCH13122 | RSV-A/human/USA/IL-LCH-LCH13122/2024 | 2024-2025 | Dec-2024 | North America/USA/Illinois | pediatric | Nasopharyngeal | A.D.5.1 | PZ279367 |

| | | | | | | | | |
|---|---|---|---|---|---|---|---|---|
| LCH13022 | RSV-A/human/USA/IL-LCH-LCH13022/2024 | 2024-2025 | Dec-2024 | North America/USA/Illinois | pediatric | Nasopharyngeal | A.D.5.1 | PZ279263 |
| LCH13038 | RSV-A/human/USA/IL-LCH-LCH13038/2024 | 2024-2025 | Dec-2024 | North America/USA/Illinois | pediatric | Nasopharyngeal | A.D.5.1 | PZ279225 |
| LCH13048 | RSV-A/human/USA/IL-LCH-LCH13048/2024 | 2024-2025 | Dec-2024 | North America/USA/Illinois | pediatric | Nasopharyngeal | A.D.5.1 | PZ279353 |
| LCH13088 | RSV-A/human/USA/IL-LCH-LCH13088/2024 | 2024-2025 | Dec-2024 | North America/USA/Illinois | pediatric | Nasopharyngeal | A.D.5.3 | PZ279361 |
| LCH13128 | RSV-A/human/USA/IL-LCH-LCH13128/2024 | 2024-2025 | Dec-2024 | North America/USA/Illinois | pediatric | Nasopharyngeal | A.D.5.1 | PZ279369 |
| LCH13005 | RSV-A/human/USA/IL-LCH-LCH13005/2024 | 2024-2025 | Dec-2024 | North America/USA/Illinois | pediatric | Nasopharyngeal | A.D.5.1 | PZ279260 |
| LCH13031 | RSV-A/human/USA/IL-LCH-LCH13031/2024 | 2024-2025 | Dec-2024 | North America/USA/Illinois | pediatric | Nasopharyngeal | A.D.1 | PZ279223 |
| LCH13049 | RSV-A/human/USA/IL-LCH-LCH13049/2024 | 2024-2025 | Dec-2024 | North America/USA/Illinois | pediatric | Nasopharyngeal | A.D.5.3 | PZ279354 |
| LCH13117 | RSV-A/human/USA/IL-LCH-LCH13117/2024 | 2024-2025 | Dec-2024 | North America/USA/Illinois | pediatric | Nasopharyngeal | A.D.5.1 | PZ279365 |
| LCH13153 | RSV-A/human/USA/IL-LCH-LCH13153/2024 | 2024-2025 | Dec-2024 | North America/USA/Illinois | pediatric | Nasopharyngeal | A.D.5.3 | PZ279370 |
| LCH13172 | RSV-A/human/USA/IL-LCH-LCH13172/2024 | 2024-2025 | Dec-2024 | North America/USA/Illinois | pediatric | Nasopharyngeal | A.D.5.1 | PZ279373 |
| LCH13244 | RSV-A/human/USA/IL-LCH-LCH13244/2024 | 2024-2025 | Dec-2024 | North America/USA/Illinois | pediatric | Nasopharyngeal | A.D.5.1 | PZ279381 |
| LCH13329 | RSV-A/human/USA/IL-LCH-LCH13329/2024 | 2024-2025 | Dec-2024 | North America/USA/Illinois | pediatric | Nasopharyngeal | A.D.5.1 | PZ279390 |
| LCH13179 | RSV-A/human/USA/IL-LCH-LCH13179/2024 | 2024-2025 | Dec-2024 | North America/USA/Illinois | pediatric | Nasopharyngeal | A.D.5.1 | PZ279374 |
| LCH13193 | RSV-A/human/USA/IL-LCH-LCH13193/2024 | 2024-2025 | Dec-2024 | North America/USA/Illinois | pediatric | Nasopharyngeal | A.D.3.1 | PZ279375 |
| LCH13380 | RSV-A/human/USA/IL-LCH-LCH13380/2024 | 2024-2025 | Dec-2024 | North America/USA/Illinois | pediatric | Nasopharyngeal | A.D.5.1 | PZ279395 |
| LCH13428 | RSV-A/human/USA/IL-LCH-LCH13428/2024 | 2024-2025 | Dec-2024 | North America/USA/Illinois | pediatric | Nasopharyngeal | A.D.1 | PZ279265 |
| LCH13216 | RSV-A/human/USA/IL-LCH-LCH13216/2024 | 2024-2025 | Dec-2024 | North America/USA/Illinois | pediatric | Nasopharyngeal | A.D.5.1 | PZ279378 |
| LCH13325 | RSV-A/human/USA/IL-LCH-LCH13325/2024 | 2024-2025 | Dec-2024 | North America/USA/Illinois | pediatric | Nasopharyngeal | A.D.5.1 | PZ279388 |
| LCH13328 | RSV-A/human/USA/IL-LCH-LCH13328/2024 | 2024-2025 | Dec-2024 | North America/USA/Illinois | pediatric | Nasopharyngeal | A.D.1 | PZ279389 |
| LCH13336 | RSV-A/human/USA/IL-LCH-LCH13336/2024 | 2024-2025 | Dec-2024 | North America/USA/Illinois | pediatric | Nasopharyngeal | A.D.1 | PZ279391 |
| LCH13344 | RSV-A/human/USA/IL-LCH-LCH13344/2024 | 2024-2025 | Dec-2024 | North America/USA/Illinois | pediatric | Nasopharyngeal | A.D.5.1 | PZ279392 |
| LCH13389 | RSV-A/human/USA/IL-LCH-LCH13389/2024 | 2024-2025 | Dec-2024 | North America/USA/Illinois | pediatric | Nasopharyngeal | A.D.1.5 | PZ279397 |
| LCH13170 | RSV-A/human/USA/IL-LCH-LCH13170/2024 | 2024-2025 | Dec-2024 | North America/USA/Illinois | pediatric | Nasopharyngeal | A.D.1 | PZ279372 |
| LCH13200 | RSV-A/human/USA/IL-LCH-LCH13200/2024 | 2024-2025 | Dec-2024 | North America/USA/Illinois | pediatric | Nasopharyngeal | A.D.3 | PZ279376 |
| LCH13212 | RSV-A/human/USA/IL-LCH-LCH13212/2024 | 2024-2025 | Dec-2024 | North America/USA/Illinois | pediatric | Nasopharyngeal | A.D.5.1 | PZ279377 |
| LCH13250 | RSV-A/human/USA/IL-LCH-LCH13250/2024 | 2024-2025 | Dec-2024 | North America/USA/Illinois | pediatric | Nasopharyngeal | A.D.1 | PZ279383 |
| LCH13232 | RSV-A/human/USA/IL-LCH-LCH13232/2024 | 2024-2025 | Dec-2024 | North America/USA/Illinois | pediatric | Nasopharyngeal | A.D.5.1 | PZ279380 |
| LCH13294 | RSV-A/human/USA/IL-LCH-LCH13294/2024 | 2024-2025 | Dec-2024 | North America/USA/Illinois | pediatric | Nasopharyngeal | A.D.5.2 | PZ279387 |
| LCH13386 | RSV-A/human/USA/IL-LCH-LCH13386/2024 | 2024-2025 | Dec-2024 | North America/USA/Illinois | pediatric | Nasopharyngeal | A.D.3.1 | PZ279396 |

| | | | | | | | | |
|---|---|---|---|---|---|---|---|---|
| LCH13262 | RSV-A/human/USA/IL-LCH-LCH13262/2024 | 2024-2025 | Dec-2024 | North America/USA/Illinois | pediatric | Nasopharyngeal | A.D.5.1 | PZ279385 |
| LCH13265 | RSV-A/human/USA/IL-LCH-LCH13265/2024 | 2024-2025 | Dec-2024 | North America/USA/Illinois | pediatric | Nasopharyngeal | A.D.5.1 | PZ279386 |
| LCH13357 | RSV-A/human/USA/IL-LCH-LCH13357/2024 | 2024-2025 | Dec-2024 | North America/USA/Illinois | pediatric | Nasopharyngeal | A.D.5.1 | PZ279393 |
| LCH13369 | RSV-A/human/USA/IL-LCH-LCH13369/2024 | 2024-2025 | Dec-2024 | North America/USA/Illinois | pediatric | Nasopharyngeal | A.D.5.3 | PZ279394 |
| LCH13406 | RSV-A/human/USA/IL-LCH-LCH13406/2024 | 2024-2025 | Dec-2024 | North America/USA/Illinois | pediatric | Nasopharyngeal | A.D.5.1 | PZ279264 |
| LCH13225 | RSV-A/human/USA/IL-LCH-LCH13225/2024 | 2024-2025 | Dec-2024 | North America/USA/Illinois | pediatric | Nasopharyngeal | A.D.1 | PZ279379 |
| LCH13248 | RSV-A/human/USA/IL-LCH-LCH13248/2024 | 2024-2025 | Dec-2024 | North America/USA/Illinois | pediatric | Nasopharyngeal | A.D.1 | PZ279382 |
| LCH13256 | RSV-A/human/USA/IL-LCH-LCH13256/2024 | 2024-2025 | Dec-2024 | North America/USA/Illinois | pediatric | Nasopharyngeal | A.D.5.1 | PZ279384 |
| LCH13533 | RSV-A/human/USA/IL-LCH-LCH13533/2024 | 2024-2025 | Dec-2024 | North America/USA/Illinois | pediatric | Nasopharyngeal | A.D.3.1 | PZ279276 |
| LCH13548 | RSV-A/human/USA/IL-LCH-LCH13548/2024 | 2024-2025 | Dec-2024 | North America/USA/Illinois | pediatric | Nasopharyngeal | A.D.5.1 | PZ279280 |
| LCH13699 | RSV-A/human/USA/IL-LCH-LCH13699/2024 | 2024-2025 | Dec-2024 | North America/USA/Illinois | pediatric | Nasopharyngeal | A.D.5.1 | PZ279292 |
| LCH13730 | RSV-A/human/USA/IL-LCH-LCH13730/2024 | 2024-2025 | Dec-2024 | North America/USA/Illinois | pediatric | Nasopharyngeal | A.D.5.1 | PZ279398 |
| LCH13500 | RSV-A/human/USA/IL-LCH-LCH13500/2024 | 2024-2025 | Dec-2024 | North America/USA/Illinois | pediatric | Nasopharyngeal | A.D.5.1 | PZ279271 |
| LCH13503 | RSV-A/human/USA/IL-LCH-LCH13503/2024 | 2024-2025 | Dec-2024 | North America/USA/Illinois | pediatric | Nasopharyngeal | A.D.5.1 | PZ279272 |
| LCH13696 | RSV-A/human/USA/IL-LCH-LCH13696/2024 | 2024-2025 | Dec-2024 | North America/USA/Illinois | pediatric | Nasopharyngeal | A.D.3 | PZ279291 |
| LCH13735 | RSV-A/human/USA/IL-LCH-LCH13735/2024 | 2024-2025 | Dec-2024 | North America/USA/Illinois | pediatric | Nasopharyngeal | A.D.1 | PZ279399 |
| LCH13433 | RSV-A/human/USA/IL-LCH-LCH13433/2024 | 2024-2025 | Dec-2024 | North America/USA/Illinois | pediatric | Nasopharyngeal | A.D.5.3 | PZ279266 |
| LCH13470 | RSV-A/human/USA/IL-LCH-LCH13470/2024 | 2024-2025 | Dec-2024 | North America/USA/Illinois | pediatric | Nasopharyngeal | A.D.5.1 | PZ279269 |
| LCH13561 | RSV-A/human/USA/IL-LCH-LCH13561/2024 | 2024-2025 | Dec-2024 | North America/USA/Illinois | pediatric | Nasopharyngeal | A.D.5.3 | PZ279281 |
| LCH13567 | RSV-A/human/USA/IL-LCH-LCH13567/2024 | 2024-2025 | Dec-2024 | North America/USA/Illinois | pediatric | Nasopharyngeal | A.D.3.2 | PZ279283 |
| LCH13717 | RSV-A/human/USA/IL-LCH-LCH13717/2024 | 2024-2025 | Dec-2024 | North America/USA/Illinois | pediatric | Nasopharyngeal | A.D.1.6 | PZ279295 |
| LCH13506 | RSV-A/human/USA/IL-LCH-LCH13506/2024 | 2024-2025 | Dec-2024 | North America/USA/Illinois | pediatric | Nasopharyngeal | A.D.1.5 | PZ279274 |
| LCH13540 | RSV-A/human/USA/IL-LCH-LCH13540/2024 | 2024-2025 | Dec-2024 | North America/USA/Illinois | pediatric | Nasopharyngeal | A.D.5.1 | PZ279278 |
| LCH13558 | RSV-A/human/USA/IL-LCH-LCH13558/2024 | 2024-2025 | Dec-2024 | North America/USA/Illinois | pediatric | Nasopharyngeal | A.D.5.1 | |
| LCH13573 | RSV-A/human/USA/IL-LCH-LCH13573/2024 | 2024-2025 | Dec-2024 | North America/USA/Illinois | pediatric | Nasopharyngeal | A.D.5.1 | PZ279284 |
| LCH13700 | RSV-A/human/USA/IL-LCH-LCH13700/2024 | 2024-2025 | Dec-2024 | North America/USA/Illinois | pediatric | Nasopharyngeal | A.D.5.1 | PZ279293 |
| LCH13708 | RSV-A/human/USA/IL-LCH-LCH13708/2024 | 2024-2025 | Dec-2024 | North America/USA/Illinois | pediatric | Nasopharyngeal | A.D.5.1 | PZ279294 |
| LCH13722 | RSV-A/human/USA/IL-LCH-LCH13722/2024 | 2024-2025 | Dec-2024 | North America/USA/Illinois | pediatric | Nasopharyngeal | A.D.5.1 | PZ279296 |
| LCH13728 | RSV-A/human/USA/IL-LCH-LCH13728/2024 | 2024-2025 | Dec-2024 | North America/USA/Illinois | pediatric | Nasopharyngeal | A.D.5.1 | PZ279298 |
| LCH13747 | RSV-A/human/USA/IL-LCH-LCH13747/2024 | 2024-2025 | Dec-2024 | North America/USA/Illinois | pediatric | Nasopharyngeal | A.D.5.1 | PZ279400 |

| | | | | | | | | |
|---|---|---|---|---|---|---|---|---|
| LCH13488 | RSV-A/human/USA/IL-LCH-LCH13488/2024 | 2024-2025 | Dec-2024 | North America/USA/Illinois | pediatric | Nasopharyngeal | A.D.3.1 | PZ279270 |
| LCH13504 | RSV-A/human/USA/IL-LCH-LCH13504/2024 | 2024-2025 | Dec-2024 | North America/USA/Illinois | pediatric | Nasopharyngeal | A.D.5.1 | PZ279273 |
| LCH13529 | RSV-A/human/USA/IL-LCH-LCH13529/2024 | 2024-2025 | Dec-2024 | North America/USA/Illinois | pediatric | Nasopharyngeal | A.D.5.1 | PZ279275 |
| LCH13542 | RSV-A/human/USA/IL-LCH-LCH13542/2024 | 2024-2025 | Dec-2024 | North America/USA/Illinois | pediatric | Nasopharyngeal | A.D.1.6 | PZ279279 |
| LCH13725 | RSV-A/human/USA/IL-LCH-LCH13725/2024 | 2024-2025 | Dec-2024 | North America/USA/Illinois | pediatric | Nasopharyngeal | A.D.5.3 | PZ279297 |
| LCH13442 | RSV-A/human/USA/IL-LCH-LCH13442/2024 | 2024-2025 | Dec-2024 | North America/USA/Illinois | pediatric | Nasopharyngeal | A.D.5.1 | PZ279267 |
| LCH13459 | RSV-A/human/USA/IL-LCH-LCH13459/2024 | 2024-2025 | Dec-2024 | North America/USA/Illinois | pediatric | Nasopharyngeal | A.D.5.1 | PZ279268 |
| LCH13537 | RSV-A/human/USA/IL-LCH-LCH13537/2024 | 2024-2025 | Dec-2024 | North America/USA/Illinois | pediatric | Nasopharyngeal | A.D.5.1 | PZ279277 |
| LCH13565 | RSV-A/human/USA/IL-LCH-LCH13565/2024 | 2024-2025 | Dec-2024 | North America/USA/Illinois | pediatric | Nasopharyngeal | A.D.1.6 | PZ279282 |
| LCH13595 | RSV-A/human/USA/IL-LCH-LCH13595/2024 | 2024-2025 | Dec-2024 | North America/USA/Illinois | pediatric | Nasopharyngeal | A.D.1.4 | |
| LCH13597 | RSV-A/human/USA/IL-LCH-LCH13597/2024 | 2024-2025 | Dec-2024 | North America/USA/Illinois | pediatric | Nasopharyngeal | A.D.1.6 | PZ279285 |
| LCH13666 | RSV-A/human/USA/IL-LCH-LCH13666/2024 | 2024-2025 | Dec-2024 | North America/USA/Illinois | pediatric | Nasopharyngeal | A.D.1.5 | PZ279288 |
| LCH13697 | RSV-A/human/USA/IL-LCH-LCH13697/2024 | 2024-2025 | Dec-2024 | North America/USA/Illinois | pediatric | Nasopharyngeal | A.D.5.1 | |
| LCH13634 | RSV-A/human/USA/IL-LCH-LCH13634/2024 | 2024-2025 | Dec-2024 | North America/USA/Illinois | pediatric | Nasopharyngeal | A.D.1 | PZ279286 |
| LCH13640 | RSV-A/human/USA/IL-LCH-LCH13640/2024 | 2024-2025 | Dec-2024 | North America/USA/Illinois | pediatric | Nasopharyngeal | A.D.1 | PZ279287 |
| LCH13673 | RSV-A/human/USA/IL-LCH-LCH13673/2024 | 2024-2025 | Dec-2024 | North America/USA/Illinois | pediatric | Nasopharyngeal | A.D.5.1 | PZ279289 |
| LCH13677 | RSV-A/human/USA/IL-LCH-LCH13677/2024 | 2024-2025 | Dec-2024 | North America/USA/Illinois | pediatric | Nasopharyngeal | A.D.5.1 | PZ279290 |
| LCH13824 | RSV-A/human/USA/IL-LCH-LCH13824/2024 | 2024-2025 | Dec-2024 | North America/USA/Illinois | pediatric | Nasopharyngeal | A.D.3 | PZ279404 |
| LCH13825 | RSV-A/human/USA/IL-LCH-LCH13825/2024 | 2024-2025 | Dec-2024 | North America/USA/Illinois | pediatric | Nasopharyngeal | A.D.1 | PZ279405 |
| LCH13849 | RSV-A/human/USA/IL-LCH-LCH13849/2024 | 2024-2025 | Dec-2024 | North America/USA/Illinois | pediatric | Nasopharyngeal | A.D.5.1 | PZ279410 |
| LCH13851 | RSV-A/human/USA/IL-LCH-LCH13851/2024 | 2024-2025 | Dec-2024 | North America/USA/Illinois | pediatric | Nasopharyngeal | A.D.1.6 | PZ279411 |
| LCH14008 | RSV-A/human/USA/IL-LCH-LCH14008/2024 | 2024-2025 | Dec-2024 | North America/USA/Illinois | pediatric | Nasopharyngeal | A.D.1 | PZ279431 |
| LCH14059 | RSV-A/human/USA/IL-LCH-LCH14059/2024 | 2024-2025 | Dec-2024 | North America/USA/Illinois | pediatric | Nasopharyngeal | A.D.5.1 | PZ279436 |
| LCH13822 | RSV-A/human/USA/IL-LCH-LCH13822/2024 | 2024-2025 | Dec-2024 | North America/USA/Illinois | pediatric | Nasopharyngeal | A.D.5.1 | PZ279403 |
| LCH13837 | RSV-A/human/USA/IL-LCH-LCH13837/2024 | 2024-2025 | Dec-2024 | North America/USA/Illinois | pediatric | Nasopharyngeal | A.D.1 | PZ279407 |
| LCH13840 | RSV-A/human/USA/IL-LCH-LCH13840/2024 | 2024-2025 | Dec-2024 | North America/USA/Illinois | pediatric | Nasopharyngeal | A.D.5.1 | PZ279408 |
| LCH13882 | RSV-A/human/USA/IL-LCH-LCH13882/2024 | 2024-2025 | Dec-2024 | North America/USA/Illinois | pediatric | Nasopharyngeal | A.D.5.1 | PZ279415 |
| LCH14060 | RSV-A/human/USA/IL-LCH-LCH14060/2024 | 2024-2025 | Dec-2024 | North America/USA/Illinois | pediatric | Nasopharyngeal | A.D.1.5 | PZ279437 |
| LCH13833 | RSV-A/human/USA/IL-LCH-LCH13833/2024 | 2024-2025 | Dec-2024 | North America/USA/Illinois | pediatric | Nasopharyngeal | A.D.5.3 | PZ279406 |
| LCH13842 | RSV-A/human/USA/IL-LCH-LCH13842/2024 | 2024-2025 | Dec-2024 | North America/USA/Illinois | pediatric | Nasopharyngeal | A.D.1 | PZ279409 |

| | | | | | | | | |
|---|---|---|---|---|---|---|---|---|
| LCH13856 | RSV-A/human/USA/IL-LCH-LCH13856/2024 | 2024-2025 | Dec-2024 | North America/USA/Illinois | pediatric | Nasopharyngeal | A.D.5.3 | PZ279412 |
| LCH13870 | RSV-A/human/USA/IL-LCH-LCH13870/2024 | 2024-2025 | Dec-2024 | North America/USA/Illinois | pediatric | Nasopharyngeal | A.D.5.3 | PZ279413 |
| LCH13877 | RSV-A/human/USA/IL-LCH-LCH13877/2024 | 2024-2025 | Dec-2024 | North America/USA/Illinois | pediatric | Nasopharyngeal | A.D.5.3 | PZ279414 |
| LCH14016 | RSV-A/human/USA/IL-LCH-LCH14016/2024 | 2024-2025 | Dec-2024 | North America/USA/Illinois | pediatric | Nasopharyngeal | A.D.5.1 | PZ279432 |
| LCH14043 | RSV-A/human/USA/IL-LCH-LCH14043/2024 | 2024-2025 | Dec-2024 | North America/USA/Illinois | pediatric | Nasopharyngeal | A.D.3.1 | PZ279434 |
| LCH14057 | RSV-A/human/USA/IL-LCH-LCH14057/2024 | 2024-2025 | Dec-2024 | North America/USA/Illinois | pediatric | Nasopharyngeal | A.D.5.1 | PZ279435 |
| LCH14062 | RSV-A/human/USA/IL-LCH-LCH14062/2024 | 2024-2025 | Dec-2024 | North America/USA/Illinois | pediatric | Nasopharyngeal | A.D.5.1 | PZ279438 |
| LCH14075 | RSV-A/human/USA/IL-LCH-LCH14075/2024 | 2024-2025 | Dec-2024 | North America/USA/Illinois | pediatric | Nasopharyngeal | A.D.3 | PZ279226 |
| LCH13883 | RSV-A/human/USA/IL-LCH-LCH13883/2024 | 2024-2025 | Dec-2024 | North America/USA/Illinois | pediatric | Nasopharyngeal | A.D.5.1 | PZ279416 |
| LCH13895 | RSV-A/human/USA/IL-LCH-LCH13895/2024 | 2024-2025 | Dec-2024 | North America/USA/Illinois | pediatric | Nasopharyngeal | A.D.1.5 | PZ279417 |
| LCH13896 | RSV-A/human/USA/IL-LCH-LCH13896/2024 | 2024-2025 | Dec-2024 | North America/USA/Illinois | pediatric | Nasopharyngeal | A.D.5.1 | PZ279418 |
| LCH13899 | RSV-A/human/USA/IL-LCH-LCH13899/2024 | 2024-2025 | Dec-2024 | North America/USA/Illinois | pediatric | Nasopharyngeal | A.D.1.5 | PZ279419 |
| LCH13904 | RSV-A/human/USA/IL-LCH-LCH13904/2024 | 2024-2025 | Dec-2024 | North America/USA/Illinois | pediatric | Nasopharyngeal | A.D.5.1 | PZ279420 |
| LCH13905 | RSV-A/human/USA/IL-LCH-LCH13905/2024 | 2024-2025 | Dec-2024 | North America/USA/Illinois | pediatric | Nasopharyngeal | A.D.5.1 | PZ279421 |
| LCH13916 | RSV-A/human/USA/IL-LCH-LCH13916/2024 | 2024-2025 | Dec-2024 | North America/USA/Illinois | pediatric | Nasopharyngeal | A.D.5.1 | PZ279423 |
| LCH13970 | RSV-A/human/USA/IL-LCH-LCH13970/2024 | 2024-2025 | Dec-2024 | North America/USA/Illinois | pediatric | Nasopharyngeal | A.D.1 | PZ279430 |
| LCH14027 | RSV-A/human/USA/IL-LCH-LCH14027/2024 | 2024-2025 | Dec-2024 | North America/USA/Illinois | pediatric | Nasopharyngeal | A.D.1 | PZ279433 |
| LCH13906 | RSV-A/human/USA/IL-LCH-LCH13906/2024 | 2024-2025 | Dec-2024 | North America/USA/Illinois | pediatric | Nasopharyngeal | A.D.1 | PZ279422 |
| LCH13749 | RSV-A/human/USA/IL-LCH-LCH13749/2025 | 2024-2025 | Jan-2025 | North America/USA/Illinois | pediatric | Nasopharyngeal | A.D.1.5 | PZ279401 |
| LCH13750 | RSV-A/human/USA/IL-LCH-LCH13750/2025 | 2024-2025 | Jan-2025 | North America/USA/Illinois | pediatric | Nasopharyngeal | A.D.5.1 | PZ279402 |
| LCH13939 | RSV-A/human/USA/IL-LCH-LCH13939/2025 | 2024-2025 | Jan-2025 | North America/USA/Illinois | pediatric | Nasopharyngeal | A.D.5.1 | PZ279426 |
| LCH13954 | RSV-A/human/USA/IL-LCH-LCH13954/2025 | 2024-2025 | Jan-2025 | North America/USA/Illinois | pediatric | Nasopharyngeal | A.D.5.1 | PZ279427 |
| LCH13956 | RSV-A/human/USA/IL-LCH-LCH13956/2025 | 2024-2025 | Jan-2025 | North America/USA/Illinois | pediatric | Nasopharyngeal | A.D.1.6 | PZ279428 |
| LCH13924 | RSV-A/human/USA/IL-LCH-LCH13924/2025 | 2024-2025 | Jan-2025 | North America/USA/Illinois | pediatric | Nasopharyngeal | A.D.5.1 | PZ279424 |
| LCH13959 | RSV-A/human/USA/IL-LCH-LCH13959/2025 | 2024-2025 | Jan-2025 | North America/USA/Illinois | pediatric | Nasopharyngeal | A.D.5.1 | PZ279429 |
| LCH13938 | RSV-A/human/USA/IL-LCH-LCH13938/2025 | 2024-2025 | Jan-2025 | North America/USA/Illinois | pediatric | Nasopharyngeal | A.D.5.1 | PZ279425 |
| LCH14099 | RSV-A/human/USA/IL-LCH-LCH14099/2025 | 2024-2025 | Jan-2025 | North America/USA/Illinois | pediatric | Nasopharyngeal | A.D.1 | PZ279227 |
| LCH14226 | RSV-A/human/USA/IL-LCH-LCH14226/2025 | 2024-2025 | Jan-2025 | North America/USA/Illinois | pediatric | Nasopharyngeal | A.D.1.6 | PZ279300 |
| LCH14233 | RSV-A/human/USA/IL-LCH-LCH14233/2025 | 2024-2025 | Jan-2025 | North America/USA/Illinois | pediatric | Nasopharyngeal | A.D.5.1 | PZ279301 |
| LCH14268 | RSV-A/human/USA/IL-LCH-LCH14268/2025 | 2024-2025 | Jan-2025 | North America/USA/Illinois | pediatric | Nasopharyngeal | A.D.1 | PZ279302 |

| | | | | | | | | |
|---|---|---|---|---|---|---|---|---|
| LCH14277 | RSV-A/human/USA/IL-LCH-LCH14277/2025 | 2024-2025 | Jan-2025 | North America/USA/Illinois | pediatric | Nasopharyngeal | A.D.5.3 | PZ279304 |
| LCH14317 | RSV-A/human/USA/IL-LCH-LCH14317/2025 | 2024-2025 | Jan-2025 | North America/USA/Illinois | pediatric | Nasopharyngeal | A.D.5.3 | PZ279306 |
| LCH14118 | RSV-A/human/USA/IL-LCH-LCH14118/2025 | 2024-2025 | Jan-2025 | North America/USA/Illinois | pediatric | Nasopharyngeal | A.D.5.1 | PZ279228 |
| LCH14191 | RSV-A/human/USA/IL-LCH-LCH14191/2025 | 2024-2025 | Jan-2025 | North America/USA/Illinois | pediatric | Nasopharyngeal | A.D.1 | PZ279229 |
| LCH14195 | RSV-A/human/USA/IL-LCH-LCH14195/2025 | 2024-2025 | Jan-2025 | North America/USA/Illinois | pediatric | Nasopharyngeal | A.D.1 | PZ279299 |
| LCH14201 | RSV-A/human/USA/IL-LCH-LCH14201/2025 | 2024-2025 | Jan-2025 | North America/USA/Illinois | pediatric | Nasopharyngeal | A.D.5.2 | |
| LCH14276 | RSV-A/human/USA/IL-LCH-LCH14276/2025 | 2024-2025 | Jan-2025 | North America/USA/Illinois | pediatric | Nasopharyngeal | A.D.5.1 | PZ279303 |
| LCH14294 | RSV-A/human/USA/IL-LCH-LCH14294/2025 | 2024-2025 | Jan-2025 | North America/USA/Illinois | pediatric | Nasopharyngeal | A.D.5.2 | |
| LCH14296 | RSV-A/human/USA/IL-LCH-LCH14296/2025 | 2024-2025 | Jan-2025 | North America/USA/Illinois | pediatric | Nasopharyngeal | A.D.5.2 | PZ279305 |
| LCH14554 | RSV-A/human/USA/IL-LCH-LCH14554/2025 | 2024-2025 | Jan-2025 | North America/USA/Illinois | pediatric | Nasopharyngeal | A.D.5.3 | PZ279314 |
| LCH14588 | RSV-A/human/USA/IL-LCH-LCH14588/2025 | 2024-2025 | Jan-2025 | North America/USA/Illinois | pediatric | Nasopharyngeal | A.D.1.6 | PZ279318 |
| LCH14355 | RSV-A/human/USA/IL-LCH-LCH14355/2025 | 2024-2025 | Jan-2025 | North America/USA/Illinois | pediatric | Nasopharyngeal | A.D.1 | PZ279311 |
| LCH14556 | RSV-A/human/USA/IL-LCH-LCH14556/2025 | 2024-2025 | Jan-2025 | North America/USA/Illinois | pediatric | Nasopharyngeal | A.D.5.1 | PZ279315 |
| LCH14577 | RSV-A/human/USA/IL-LCH-LCH14577/2025 | 2024-2025 | Jan-2025 | North America/USA/Illinois | pediatric | Nasopharyngeal | A.D.1 | PZ279317 |
| LCH14337 | RSV-A/human/USA/IL-LCH-LCH14337/2025 | 2024-2025 | Jan-2025 | North America/USA/Illinois | pediatric | Nasopharyngeal | A.D.1.6 | PZ279308 |
| LCH14533 | RSV-A/human/USA/IL-LCH-LCH14533/2025 | 2024-2025 | Jan-2025 | North America/USA/Illinois | pediatric | Nasopharyngeal | A.D.3.3 | PZ279313 |
| LCH14557 | RSV-A/human/USA/IL-LCH-LCH14557/2025 | 2024-2025 | Jan-2025 | North America/USA/Illinois | pediatric | Nasopharyngeal | A.D.3 | PZ279316 |
| LCH14413 | RSV-A/human/USA/IL-LCH-LCH14413/2025 | 2024-2025 | Jan-2025 | North America/USA/Illinois | pediatric | Nasopharyngeal | A.D.5.3 | |
| LCH14636 | RSV-A/human/USA/IL-LCH-LCH14636/2025 | 2024-2025 | Jan-2025 | North America/USA/Illinois | pediatric | Nasopharyngeal | A.D.1.6 | PZ279320 |
| LCH14345 | RSV-A/human/USA/IL-LCH-LCH14345/2025 | 2024-2025 | Jan-2025 | North America/USA/Illinois | pediatric | Nasopharyngeal | A.D.3.1 | PZ279309 |
| LCH14350 | RSV-A/human/USA/IL-LCH-LCH14350/2025 | 2024-2025 | Jan-2025 | North America/USA/Illinois | pediatric | Nasopharyngeal | A.D.1.6 | PZ279310 |
| LCH14335 | RSV-A/human/USA/IL-LCH-LCH14335/2025 | 2024-2025 | Jan-2025 | North America/USA/Illinois | pediatric | Nasopharyngeal | A.D.5.1 | PZ279307 |
| LCH14617 | RSV-A/human/USA/IL-LCH-LCH14617/2025 | 2024-2025 | Jan-2025 | North America/USA/Illinois | pediatric | Nasopharyngeal | A.D.1 | PZ279319 |
| LCH14456 | RSV-A/human/USA/IL-LCH-LCH14456/2025 | 2024-2025 | Jan-2025 | North America/USA/Illinois | pediatric | Nasopharyngeal | A.D.5.1 | PZ279312 |
| LCH14871 | RSV-A/human/USA/IL-LCH-LCH14871/2025 | 2024-2025 | Jan-2025 | North America/USA/Illinois | pediatric | Nasopharyngeal | A.D.3.1 | PZ279327 |
| LCH14750 | RSV-A/human/USA/IL-LCH-LCH14750/2025 | 2024-2025 | Jan-2025 | North America/USA/Illinois | pediatric | Nasopharyngeal | A.D.5.1 | PZ279323 |
| LCH14783 | RSV-A/human/USA/IL-LCH-LCH14783/2025 | 2024-2025 | Jan-2025 | North America/USA/Illinois | pediatric | Nasopharyngeal | A.D.1 | PZ279325 |
| LCH14738 | RSV-A/human/USA/IL-LCH-LCH14738/2025 | 2024-2025 | Jan-2025 | North America/USA/Illinois | pediatric | Nasopharyngeal | A.D.1.6 | PZ279322 |
| LCH14781 | RSV-A/human/USA/IL-LCH-LCH14781/2025 | 2024-2025 | Jan-2025 | North America/USA/Illinois | pediatric | Nasopharyngeal | A.D.5.1 | PZ279324 |
| LCH14870 | RSV-A/human/USA/IL-LCH-LCH14870/2025 | 2024-2025 | Jan-2025 | North America/USA/Illinois | pediatric | Nasopharyngeal | A.D.3.2 | PZ279326 |

| | | | | | | | | |
|---|---|---|---|---|---|---|---|---|
| LCH14730 | RSV-A/human/USA/IL-LCH-LCH14730/2025 | 2024-2025 | Jan-2025 | North America/USA/Illinois | pediatric | Nasopharyngeal | A.D.1 | PZ279321 |
| LCH15352 | RSV-A/human/USA/IL-LCH-LCH15352/2025 | 2024-2025 | Jan-2025 | North America/USA/Illinois | pediatric | Nasopharyngeal | A.D.5.1 | PZ279334 |
| LCH15150 | RSV-A/human/USA/IL-LCH-LCH15150/2025 | 2024-2025 | Jan-2025 | North America/USA/Illinois | pediatric | Nasopharyngeal | A.D.3.1 | PZ279328 |
| LCH15165 | RSV-A/human/USA/IL-LCH-LCH15165/2025 | 2024-2025 | Jan-2025 | North America/USA/Illinois | pediatric | Nasopharyngeal | A.D.1.5 | PZ279329 |
| LCH15171 | RSV-A/human/USA/IL-LCH-LCH15171/2025 | 2024-2025 | Jan-2025 | North America/USA/Illinois | pediatric | Nasopharyngeal | A.D.5.1 | PZ279330 |
| LCH15242 | RSV-A/human/USA/IL-LCH-LCH15242/2025 | 2024-2025 | Feb-2025 | North America/USA/Illinois | pediatric | Nasopharyngeal | A.D.5.3 | PZ279332 |
| LCH15279 | RSV-A/human/USA/IL-LCH-LCH15279/2025 | 2024-2025 | Feb-2025 | North America/USA/Illinois | pediatric | Nasopharyngeal | A.D.5.1 | PZ279333 |
| LCH15548 | RSV-A/human/USA/IL-LCH-LCH15548/2025 | 2024-2025 | Feb-2025 | North America/USA/Illinois | pediatric | Nasopharyngeal | A.D.1.6 | PZ279343 |
| LCH15554 | RSV-A/human/USA/IL-LCH-LCH15554/2025 | 2024-2025 | Feb-2025 | North America/USA/Illinois | pediatric | Nasopharyngeal | A.D.5.1 | PZ279344 |
| LCH15238 | RSV-A/human/USA/IL-LCH-LCH15238/2025 | 2024-2025 | Feb-2025 | North America/USA/Illinois | pediatric | Nasopharyngeal | A.D.5.1 | PZ279331 |
| LCH15430 | RSV-A/human/USA/IL-LCH-LCH15430/2025 | 2024-2025 | Feb-2025 | North America/USA/Illinois | pediatric | Nasopharyngeal | A.D.5.1 | PZ279336 |
| LCH15507 | RSV-A/human/USA/IL-LCH-LCH15507/2025 | 2024-2025 | Feb-2025 | North America/USA/Illinois | pediatric | Nasopharyngeal | A.D.1.6 | PZ279341 |
| LCH15513 | RSV-A/human/USA/IL-LCH-LCH15513/2025 | 2024-2025 | Feb-2025 | North America/USA/Illinois | pediatric | Nasopharyngeal | A.D.3 | PZ279342 |
| LCH15438 | RSV-A/human/USA/IL-LCH-LCH15438/2025 | 2024-2025 | Feb-2025 | North America/USA/Illinois | pediatric | Nasopharyngeal | A.D.5.1 | PZ279337 |
| LCH15473 | RSV-A/human/USA/IL-LCH-LCH15473/2025 | 2024-2025 | Feb-2025 | North America/USA/Illinois | pediatric | Nasopharyngeal | A.D.5.2 | PZ279339 |
| LCH15459 | RSV-A/human/USA/IL-LCH-LCH15459/2025 | 2024-2025 | Feb-2025 | North America/USA/Illinois | pediatric | Nasopharyngeal | A.D.3.3 | PZ279338 |
| LCH15483 | RSV-A/human/USA/IL-LCH-LCH15483/2025 | 2024-2025 | Feb-2025 | North America/USA/Illinois | pediatric | Nasopharyngeal | A.D.1 | PZ279340 |
| LCH15367 | RSV-A/human/USA/IL-LCH-LCH15367/2025 | 2024-2025 | Feb-2025 | North America/USA/Illinois | pediatric | Nasopharyngeal | A.D.5.1 | PZ279335 |
| LCH15737 | RSV-A/human/USA/IL-LCH-LCH15737/2025 | 2024-2025 | Feb-2025 | North America/USA/Illinois | pediatric | Nasopharyngeal | A.D.5.1 | PZ279232 |
| LCH15798 | RSV-A/human/USA/IL-LCH-LCH15798/2025 | 2024-2025 | Feb-2025 | North America/USA/Illinois | pediatric | Nasopharyngeal | A.D.5.1 | PZ279235 |
| LCH15785 | RSV-A/human/USA/IL-LCH-LCH15785/2025 | 2024-2025 | Feb-2025 | North America/USA/Illinois | pediatric | Nasopharyngeal | A.D.1.6 | PZ279233 |
| LCH15788 | RSV-A/human/USA/IL-LCH-LCH15788/2025 | 2024-2025 | Feb-2025 | North America/USA/Illinois | pediatric | Nasopharyngeal | A.D.1 | PZ279234 |
| LCH15802 | RSV-A/human/USA/IL-LCH-LCH15802/2025 | 2024-2025 | Feb-2025 | North America/USA/Illinois | pediatric | Nasopharyngeal | A.D.1.6 | PZ279236 |
| LCH15576 | RSV-A/human/USA/IL-LCH-LCH15576/2025 | 2024-2025 | Feb-2025 | North America/USA/Illinois | pediatric | Nasopharyngeal | A.D.5.1 | PZ279345 |
| LCH15618 | RSV-A/human/USA/IL-LCH-LCH15618/2025 | 2024-2025 | Feb-2025 | North America/USA/Illinois | pediatric | Nasopharyngeal | A.D.3.1 | PZ279346 |
| LCH15684 | RSV-A/human/USA/IL-LCH-LCH15684/2025 | 2024-2025 | Feb-2025 | North America/USA/Illinois | pediatric | Nasopharyngeal | A.D.3.2 | PZ279230 |
| LCH15707 | RSV-A/human/USA/IL-LCH-LCH15707/2025 | 2024-2025 | Feb-2025 | North America/USA/Illinois | pediatric | Nasopharyngeal | A.D.5.1 | PZ279231 |
| LCH15861 | RSV-A/human/USA/IL-LCH-LCH15861/2025 | 2024-2025 | Feb-2025 | North America/USA/Illinois | pediatric | Nasopharyngeal | A.D.5.2 | PZ279237 |
| LCH15962 | RSV-A/human/USA/IL-LCH-LCH15962/2025 | 2024-2025 | Feb-2025 | North America/USA/Illinois | pediatric | Nasopharyngeal | A.D.3.1 | PZ279241 |
| LCH15916 | RSV-A/human/USA/IL-LCH-LCH15916/2025 | 2024-2025 | Feb-2025 | North America/USA/Illinois | pediatric | Nasopharyngeal | A.D.3 | PZ279239 |

| | | | | | | | | |
|---|---|---|---|---|---|---|---|---|
| LCH16015 | RSV-A/human/USA/IL-LCH-LCH16015/2025 | 2024-2025 | Feb-2025 | North America/USA/Illinois | pediatric | Nasopharyngeal | A.D.3 | PZ279242 |
| LCH15937 | RSV-A/human/USA/IL-LCH-LCH15937/2025 | 2024-2025 | Feb-2025 | North America/USA/Illinois | pediatric | Nasopharyngeal | A.D.3.1 | PZ279240 |
| LCH15900 | RSV-A/human/USA/IL-LCH-LCH15900/2025 | 2024-2025 | Feb-2025 | North America/USA/Illinois | pediatric | Nasopharyngeal | A.D.5.1 | PZ279238 |
| LCH16099 | RSV-A/human/USA/IL-LCH-LCH16099/2025 | 2024-2025 | Feb-2025 | North America/USA/Illinois | pediatric | Nasopharyngeal | A.D.1 | PZ279243 |
| LCH16213 | RSV-A/human/USA/IL-LCH-LCH16213/2025 | 2024-2025 | Mar-2025 | North America/USA/Illinois | pediatric | Nasopharyngeal | A.D.1.6 | PZ279244 |
| LCH16232 | RSV-A/human/USA/IL-LCH-LCH16232/2025 | 2024-2025 | Mar-2025 | North America/USA/Illinois | pediatric | Nasopharyngeal | A.D.1.6 | PZ279245 |
| LCH16252 | RSV-A/human/USA/IL-LCH-LCH16252/2025 | 2024-2025 | Mar-2025 | North America/USA/Illinois | pediatric | Nasopharyngeal | A.D.5.1 | PZ279246 |
| LCH16295 | RSV-A/human/USA/IL-LCH-LCH16295/2025 | 2024-2025 | Mar-2025 | North America/USA/Illinois | pediatric | Nasopharyngeal | A.D.5.1 | PZ279247 |
| LCH16296 | RSV-A/human/USA/IL-LCH-LCH16296/2025 | 2024-2025 | Mar-2025 | North America/USA/Illinois | pediatric | Nasopharyngeal | A.D.3.1 | PZ279248 |
| LCH16310 | RSV-A/human/USA/IL-LCH-LCH16310/2025 | 2024-2025 | Mar-2025 | North America/USA/Illinois | pediatric | Nasopharyngeal | A.D.3 | PZ279249 |
| LCH16355 | RSV-A/human/USA/IL-LCH-LCH16355/2025 | 2024-2025 | Mar-2025 | North America/USA/Illinois | pediatric | Nasopharyngeal | A.D.1.6 | PZ279250 |
| LCH16400 | RSV-A/human/USA/IL-LCH-LCH16400/2025 | 2024-2025 | Mar-2025 | North America/USA/Illinois | pediatric | Nasopharyngeal | A.D.5.1 | PZ279252 |
| LCH16395 | RSV-A/human/USA/IL-LCH-LCH16395/2025 | 2024-2025 | Mar-2025 | North America/USA/Illinois | pediatric | Nasopharyngeal | A.D.1 | PZ279251 |
| LCH16401 | RSV-A/human/USA/IL-LCH-LCH16401/2025 | 2024-2025 | Mar-2025 | North America/USA/Illinois | pediatric | Nasopharyngeal | A.D.5.1 | PZ279253 |
| LCH16451 | RSV-A/human/USA/IL-LCH-LCH16451/2025 | 2024-2025 | Mar-2025 | North America/USA/Illinois | pediatric | Nasopharyngeal | A.D.5.1 | PZ279254 |
| LCH16504 | RSV-A/human/USA/IL-LCH-LCH16504/2025 | 2024-2025 | Mar-2025 | North America/USA/Illinois | pediatric | Nasopharyngeal | A.D.3.1 | PZ279255 |
| LCH16529 | RSV-A/human/USA/IL-LCH-LCH16529/2025 | 2024-2025 | Mar-2025 | North America/USA/Illinois | pediatric | Nasopharyngeal | A.D.3 | PZ279256 |
| LCH16562 | RSV-A/human/USA/IL-LCH-LCH16562/2025 | 2024-2025 | Mar-2025 | North America/USA/Illinois | pediatric | Nasopharyngeal | A.D.5.1 | PZ279257 |
| LCH16617 | RSV-A/human/USA/IL-LCH-LCH16617/2025 | 2024-2025 | Mar-2025 | North America/USA/Illinois | pediatric | Nasopharyngeal | A.D.1.6 | PZ279258 |
| RSV556 | RSV-B/human/USA/IL-NM-RSV556/2023 | 2022-2023 | Jan-2023 | North America / USA / Illinois | adult | Nasopharyngeal | B.D.4.1.1 | PZ279463 |
| RSV569 | RSV-B/human/USA/IL-NM-RSV569/2023 | 2022-2023 | Feb-2023 | North America / USA / Illinois | adult | Nasopharyngeal | B.D.E.1 | PZ279464 |
| RSV570 | RSV-B/human/USA/IL-NM-RSV570/2023 | 2022-2023 | Feb-2023 | North America / USA / Illinois | adult | Nasopharyngeal | B.D.E.1 | PZ279465 |
| RSV602 | RSV-B/human/USA/IL-NM-RSV602/2023 | 2023-2024 | Oct-2023 | North America / USA / Illinois | adult | Nasopharyngeal | B.D.E.1 | |
| RSV606 | RSV-B/human/USA/IL-NM-RSV606/2023 | 2023-2024 | Oct-2023 | North America / USA / Illinois | pediatric | Nasopharyngeal | B.D.E.1 | PZ279466 |
| RSV607 | RSV-B/human/USA/IL-NM-RSV607/2023 | 2023-2024 | Oct-2023 | North America / USA / Illinois | pediatric | Nasopharyngeal | B.D.E.1 | |
| RSV622 | RSV-B/human/USA/IL-NM-RSV622/2023 | 2023-2024 | Oct-2023 | North America / USA / Illinois | adult | Nasopharyngeal | B.D.E.1.4 | PZ279467 |
| RSV638 | RSV-B/human/USA/IL-NM-RSV638/2023 | 2023-2024 | Oct-2023 | North America / USA / Illinois | pediatric | Nasopharyngeal | B.D.E.1 | PZ279470 |
| RSV648 | RSV-B/human/USA/IL-NM-RSV648/2023 | 2023-2024 | Oct-2023 | North America / USA / Illinois | adult | Nasopharyngeal | B.D.E.1 | PZ279472 |
| RSV625 | RSV-B/human/USA/IL-NM-RSV625/2023 | 2023-2024 | Oct-2023 | North America / USA / Illinois | pediatric | Nasopharyngeal | B.D.E.1 | |
| RSV626 | RSV-B/human/USA/IL-NM-RSV626/2023 | 2023-2024 | Nov-2023 | North America / USA / Illinois | pediatric | Nasopharyngeal | B.D.E.1 | |

| | | | | | | | | |
|---|---|---|---|---|---|---|---|---|
| RSV632 | RSV-B/human/USA/IL-NM-RSV632/2023 | 2023-2024 | Nov-2023 | North America / USA / Illinois | adult | Nasopharyngeal | B.D.E.1 | |
| RSV628 | RSV-B/human/USA/IL-NM-RSV628/2023 | 2023-2024 | Nov-2023 | North America / USA / Illinois | adult | Nasopharyngeal | B.D.E.1 | PZ279468 |
| RSV629 | RSV-B/human/USA/IL-NM-RSV629/2023 | 2023-2024 | Nov-2023 | North America / USA / Illinois | pediatric | Nasopharyngeal | B.D.E.1 | PZ279469 |
| RSV639 | RSV-B/human/USA/IL-NM-RSV639/2023 | 2023-2024 | Nov-2023 | North America / USA / Illinois | pediatric | Nasopharyngeal | B.D.E.1 | |
| RSV640 | RSV-B/human/USA/IL-NM-RSV640/2023 | 2023-2024 | Nov-2023 | North America / USA / Illinois | pediatric | Nasopharyngeal | B.D.E.1 | PZ279471 |
| RSV650 | RSV-B/human/USA/IL-NM-RSV650/2023 | 2023-2024 | Nov-2023 | North America / USA / Illinois | pediatric | Nasopharyngeal | B.D.E.1 | PZ279473 |
| RSV657 | RSV-B/human/USA/IL-NM-RSV657/2023 | 2023-2024 | Nov-2023 | North America / USA / Illinois | adult | Nasopharyngeal | B.D.E.1 | |
| RSV662 | RSV-B/human/USA/IL-NM-RSV662/2023 | 2023-2024 | Nov-2023 | North America / USA / Illinois | adult | Nasopharyngeal | B.D.E.1.4 | PZ279474 |
| RSV677 | RSV-B/human/USA/IL-NM-RSV677/2023 | 2023-2024 | Dec-2023 | North America / USA / Illinois | adult | Nasopharyngeal | B.D.E.1 | PZ279475 |
| RSV695 | RSV-B/human/USA/IL-NM-RSV695/2023 | 2023-2024 | Dec-2023 | North America / USA / Illinois | adult | Nasopharyngeal | B.D.E.1 | PZ279478 |
| RSV696 | RSV-B/human/USA/IL-NM-RSV696/2023 | 2023-2024 | Dec-2023 | North America / USA / Illinois | pediatric | Nasopharyngeal | B.D.E.1 | PZ279479 |
| RSV698 | RSV-B/human/USA/IL-NM-RSV698/2023 | 2023-2024 | Dec-2023 | North America / USA / Illinois | pediatric | Nasopharyngeal | B.D.E.1 | PZ279480 |
| RSV722 | RSV-B/human/USA/IL-NM-RSV722/2023 | 2023-2024 | Dec-2023 | North America / USA / Illinois | pediatric | Nasopharyngeal | B.D.E.1 | |
| RSV726 | RSV-B/human/USA/IL-NM-RSV726/2023 | 2023-2024 | Dec-2023 | North America / USA / Illinois | pediatric | Nasopharyngeal | B.D.E.1 | |
| RSV727 | RSV-B/human/USA/IL-NM-RSV727/2023 | 2023-2024 | Dec-2023 | North America / USA / Illinois | adult | Nasopharyngeal | B.D.E.1 | |
| RSV729 | RSV-B/human/USA/IL-NM-RSV729/2023 | 2023-2024 | Dec-2023 | North America / USA / Illinois | pediatric | Nasopharyngeal | B.D.E.1 | |
| RSV681 | RSV-B/human/USA/IL-NM-RSV681/2023 | 2023-2024 | Dec-2023 | North America / USA / Illinois | adult | Nasopharyngeal | B.D.E.1 | PZ279476 |
| RSV688 | RSV-B/human/USA/IL-NM-RSV688/2023 | 2023-2024 | Dec-2023 | North America / USA / Illinois | pediatric | Nasopharyngeal | B.D.E.1 | PZ279477 |
| RSV703 | RSV-B/human/USA/IL-NM-RSV703/2023 | 2023-2024 | Dec-2023 | North America / USA / Illinois | pediatric | Nasopharyngeal | B.D.E.1 | |
| RSV753 | RSV-B/human/USA/IL-NM-RSV753/2023 | 2023-2024 | Dec-2023 | North America / USA / Illinois | pediatric | Nasopharyngeal | B.D.E.1 | |
| RSV707 | RSV-B/human/USA/IL-NM-RSV707/2023 | 2023-2024 | Dec-2023 | North America / USA / Illinois | pediatric | Nasopharyngeal | B.D.E.1 | |
| RSV708 | RSV-B/human/USA/IL-NM-RSV708/2023 | 2023-2024 | Dec-2023 | North America / USA / Illinois | adult | Nasopharyngeal | B.D.E.1.4 | PZ279481 |
| RSV716 | RSV-B/human/USA/IL-NM-RSV716/2023 | 2023-2024 | Dec-2023 | North America / USA / Illinois | adult | Nasopharyngeal | B.D.4.1.1 | PZ279482 |
| RSV732 | RSV-B/human/USA/IL-NM-RSV732/2023 | 2023-2024 | Dec-2023 | North America / USA / Illinois | adult | Nasopharyngeal | B.D.E.1 | |
| RSV766 | RSV-B/human/USA/IL-NM-RSV766/2023 | 2023-2024 | Dec-2023 | North America / USA / Illinois | pediatric | Nasopharyngeal | B.D.E.1 | PZ279487 |
| RSV748 | RSV-B/human/USA/IL-NM-RSV748/2023 | 2023-2024 | Dec-2023 | North America / USA / Illinois | adult | Nasopharyngeal | B.D.E.1 | |
| RSV760 | RSV-B/human/USA/IL-NM-RSV760/2023 | 2023-2024 | Dec-2023 | North America / USA / Illinois | pediatric | Nasopharyngeal | B.D.E.1 | PZ279485 |
| RSV763 | RSV-B/human/USA/IL-NM-RSV763/2023 | 2023-2024 | Dec-2023 | North America / USA / Illinois | pediatric | Nasopharyngeal | B.D.E.1 | PZ279486 |
| RSV764 | RSV-B/human/USA/IL-NM-RSV764/2023 | 2023-2024 | Dec-2023 | North America / USA / Illinois | adult | Nasopharyngeal | B.D.E.1 | |
| RSV751 | RSV-B/human/USA/IL-NM-RSV751/2023 | 2023-2024 | Dec-2023 | North America / USA / Illinois | adult | Nasopharyngeal | B.D.E.1 | PZ279483 |

| | | | | | | | | |
|---|---|---|---|---|---|---|---|---|
| RSV752 | RSV-B/human/USA/IL-NM-RSV752/2023 | 2023-2024 | Dec-2023 | North America / USA / Illinois | pediatric | Nasopharyngeal | B.D.E.1.4 | PZ279484 |
| RSV780 | RSV-B/human/USA/IL-NM-RSV780/2023 | 2023-2024 | Dec-2023 | North America / USA / Illinois | adult | Nasopharyngeal | B.D.E.1 | PZ279489 |
| RSV772 | RSV-B/human/USA/IL-NM-RSV772/2023 | 2023-2024 | Dec-2023 | North America / USA / Illinois | pediatric | Nasopharyngeal | B.D.E.1 | |
| RSV776 | RSV-B/human/USA/IL-NM-RSV776/2023 | 2023-2024 | Dec-2023 | North America / USA / Illinois | adult | Nasopharyngeal | B.D.4.1.1 | |
| RSV778 | RSV-B/human/USA/IL-NM-RSV778/2023 | 2023-2024 | Dec-2023 | North America / USA / Illinois | pediatric | Nasopharyngeal | B.D.E.1 | PZ279488 |
| RSV782 | RSV-B/human/USA/IL-NM-RSV782/2023 | 2023-2024 | Dec-2023 | North America / USA / Illinois | pediatric | Nasopharyngeal | B.D.E.1 | PZ279490 |
| RSV784 | RSV-B/human/USA/IL-NM-RSV784/2023 | 2023-2024 | Dec-2023 | North America / USA / Illinois | pediatric | Nasopharyngeal | B.D.E.1 | |
| RSV771 | RSV-B/human/USA/IL-NM-RSV771/2023 | 2023-2024 | Dec-2023 | North America / USA / Illinois | adult | Nasopharyngeal | B.D.E.1 | |
| RSV795 | RSV-B/human/USA/IL-NM-RSV795/2023 | 2023-2024 | Dec-2023 | North America / USA / Illinois | adult | Nasopharyngeal | B.D.E.1.4 | PZ279493 |
| RSV785 | RSV-B/human/USA/IL-NM-RSV785/2023 | 2023-2024 | Dec-2023 | North America / USA / Illinois | adult | Nasopharyngeal | B.D.E.5 | PZ279491 |
| RSV794 | RSV-B/human/USA/IL-NM-RSV794/2023 | 2023-2024 | Dec-2023 | North America / USA / Illinois | adult | Nasopharyngeal | B.D.E.1 | PZ279492 |
| RSV792 | RSV-B/human/USA/IL-NM-RSV792/2023 | 2023-2024 | Dec-2023 | North America / USA / Illinois | pediatric | Nasopharyngeal | B.D.E.1 | |
| RSV803 | RSV-B/human/USA/IL-NM-RSV803/2024 | 2023-2024 | Jan-2024 | North America / USA / Illinois | adult | Nasopharyngeal | B.D.E.1 | |
| RSV808 | RSV-B/human/USA/IL-NM-RSV808/2024 | 2023-2024 | Jan-2024 | North America / USA / Illinois | pediatric | Nasopharyngeal | B.D.E.1 | |
| RSV815 | RSV-B/human/USA/IL-NM-RSV815/2024 | 2023-2024 | Feb-2024 | North America / USA / Illinois | adult | Nasopharyngeal | B.D.E.1 | PZ279503 |
| RSV816 | RSV-B/human/USA/IL-NM-RSV816/2024 | 2023-2024 | Feb-2024 | North America / USA / Illinois | pediatric | Nasopharyngeal | B.D.E.1 | |
| RSV818 | RSV-B/human/USA/IL-NM-RSV818/2024 | 2023-2024 | Feb-2024 | North America / USA / Illinois | pediatric | Nasopharyngeal | B.D.E.1 | PZ279504 |
| RSV820 | RSV-B/human/USA/IL-NM-RSV820/2024 | 2023-2024 | Mar-2024 | North America / USA / Illinois | adult | Nasopharyngeal | B.D.E.1 | |
| LCH12824 | RSV-B/human/USA/IL-LCH-LCH12824/2024 | 2024-2025 | Nov-2024 | North America / USA / Illinois | pediatric | Nasopharyngeal | B.D.E.1.2 | PZ279494 |
| LCH12847 | RSV-B/human/USA/IL-LCH-LCH12847/2024 | 2024-2025 | Nov-2024 | North America / USA / Illinois | pediatric | Nasopharyngeal | B.D.4.1.1 | PZ279497 |
| LCH12848 | RSV-B/human/USA/IL-LCH-LCH12848/2024 | 2024-2025 | Nov-2024 | North America / USA / Illinois | pediatric | Nasopharyngeal | B.D.4.1.1 | PZ279498 |
| LCH12833 | RSV-B/human/USA/IL-LCH-LCH12833/2024 | 2024-2025 | Nov-2024 | North America / USA / Illinois | pediatric | Nasopharyngeal | B.D.E.1.2 | PZ279495 |
| LCH12838 | RSV-B/human/USA/IL-LCH-LCH12838/2024 | 2024-2025 | Nov-2024 | North America / USA / Illinois | pediatric | Nasopharyngeal | B.D.E.1.2 | PZ279496 |
| LCH12888 | RSV-B/human/USA/IL-LCH-LCH12888/2024 | 2024-2025 | Nov-2024 | North America / USA / Illinois | pediatric | Nasopharyngeal | B.D.E.1.2 | PZ279499 |
| LCH12902 | RSV-B/human/USA/IL-LCH-LCH12902/2024 | 2024-2025 | Nov-2024 | North America / USA / Illinois | pediatric | Nasopharyngeal | B.D.E.1.2 | PZ279501 |
| LCH12897 | RSV-B/human/USA/IL-LCH-LCH12897/2024 | 2024-2025 | Nov-2024 | North America / USA / Illinois | pediatric | Nasopharyngeal | B.D.E.1.2 | PZ279500 |
| LCH12927 | RSV-B/human/USA/IL-LCH-LCH12927/2024 | 2024-2025 | Dec-2024 | North America / USA / Illinois | pediatric | Nasopharyngeal | B.D.E.1.2 | PZ279502 |
| LCH13105 | RSV-B/human/USA/IL-LCH-LCH13105/2024 | 2024-2025 | Dec-2024 | North America / USA / Illinois | pediatric | Nasopharyngeal | B.D.E.1.2 | PZ279507 |
| LCH13095 | RSV-B/human/USA/IL-LCH-LCH13095/2024 | 2024-2025 | Dec-2024 | North America / USA / Illinois | pediatric | Nasopharyngeal | B.D.E.1.2 | PZ279506 |
| LCH13162 | RSV-B/human/USA/IL-LCH-LCH13162/2024 | 2024-2025 | Dec-2024 | North America / USA / Illinois | pediatric | Nasopharyngeal | B.D.E.1 | |

| | | | | | | | | |
|---|---|---|---|---|---|---|---|---|
| LCH13131 | RSV-B/human/USA/IL-LCH-LCH13131/2024 | 2024-2025 | Dec-2024 | North America / USA / Illinois | pediatric | Nasopharyngeal | B.D.E.1 | |
| LCH13086 | RSV-B/human/USA/IL-LCH-LCH13086/2024 | 2024-2025 | Dec-2024 | North America / USA / Illinois | pediatric | Nasopharyngeal | B.D.4.1.1 | PZ279505 |
| LCH13298 | RSV-B/human/USA/IL-LCH-LCH13298/2024 | 2024-2025 | Dec-2024 | North America / USA / Illinois | pediatric | Nasopharyngeal | B.D.E.1.2 | PZ279508 |
| LCH13570 | RSV-B/human/USA/IL-LCH-LCH13570/2024 | 2024-2025 | Dec-2024 | North America / USA / Illinois | pediatric | Nasopharyngeal | B.D.E.1 | PZ279511 |
| LCH13698 | RSV-B/human/USA/IL-LCH-LCH13698/2024 | 2024-2025 | Dec-2024 | North America / USA / Illinois | pediatric | Nasopharyngeal | B.D.E.1.2 | PZ279512 |
| LCH13508 | RSV-B/human/USA/IL-LCH-LCH13508/2024 | 2024-2025 | Dec-2024 | North America / USA / Illinois | pediatric | Nasopharyngeal | B.D.E.1.2 | PZ279509 |
| LCH13726 | RSV-B/human/USA/IL-LCH-LCH13726/2024 | 2024-2025 | Dec-2024 | North America / USA / Illinois | pediatric | Nasopharyngeal | B.D.E.1.2 | PZ279513 |
| LCH13524 | RSV-B/human/USA/IL-LCH-LCH13524/2024 | 2024-2025 | Dec-2024 | North America / USA / Illinois | pediatric | Nasopharyngeal | B.D.E.1.2 | PZ279510 |
| LCH13887 | RSV-B/human/USA/IL-LCH-LCH13887/2024 | 2024-2025 | Dec-2024 | North America / USA / Illinois | pediatric | Nasopharyngeal | B.D.E.1.4 | PZ279516 |
| LCH13968 | RSV-B/human/USA/IL-LCH-LCH13968/2024 | 2024-2025 | Dec-2024 | North America / USA / Illinois | pediatric | Nasopharyngeal | B.D.E.1.2 | PZ279517 |
| LCH13768 | RSV-B/human/USA/IL-LCH-LCH13768/2025 | 2024-2025 | Jan-2025 | North America / USA / Illinois | pediatric | Nasopharyngeal | B.D.E.1.2 | PZ279515 |
| LCH13898 | RSV-B/human/USA/IL-LCH-LCH13898/2025 | 2024-2025 | Jan-2025 | North America / USA / Illinois | pediatric | Nasopharyngeal | B.D.E.1 | |
| LCH13751 | RSV-B/human/USA/IL-LCH-LCH13751/2025 | 2024-2025 | Jan-2025 | North America / USA / Illinois | pediatric | Nasopharyngeal | B.D.E.1.2 | PZ279514 |
| LCH14171 | RSV-B/human/USA/IL-LCH-LCH14171/2025 | 2024-2025 | Jan-2025 | North America / USA / Illinois | pediatric | Nasopharyngeal | B.D.E.1 | PZ279520 |
| LCH14131 | RSV-B/human/USA/IL-LCH-LCH14131/2025 | 2024-2025 | Jan-2025 | North America / USA / Illinois | pediatric | Nasopharyngeal | B.D.E.1.2 | PZ279518 |
| LCH14207 | RSV-B/human/USA/IL-LCH-LCH14207/2025 | 2024-2025 | Jan-2025 | North America / USA / Illinois | pediatric | Nasopharyngeal | B.D.E.1 | PZ279522 |
| LCH14215 | RSV-B/human/USA/IL-LCH-LCH14215/2025 | 2024-2025 | Jan-2025 | North America / USA / Illinois | pediatric | Nasopharyngeal | B.D.E.1 | |
| LCH14158 | RSV-B/human/USA/IL-LCH-LCH14158/2025 | 2024-2025 | Jan-2025 | North America / USA / Illinois | pediatric | Nasopharyngeal | B.D.E.1.2 | PZ279519 |
| LCH14180 | RSV-B/human/USA/IL-LCH-LCH14180/2025 | 2024-2025 | Jan-2025 | North America / USA / Illinois | pediatric | Nasopharyngeal | B.D.E.1.2 | PZ279521 |
| LCH14327 | RSV-B/human/USA/IL-LCH-LCH14327/2025 | 2024-2025 | Jan-2025 | North America / USA / Illinois | pediatric | Nasopharyngeal | B.D.E.1.2 | PZ279523 |
| LCH14331 | RSV-B/human/USA/IL-LCH-LCH14331/2025 | 2024-2025 | Jan-2025 | North America / USA / Illinois | pediatric | Nasopharyngeal | B.D.E.1 | PZ279524 |
| LCH14430 | RSV-B/human/USA/IL-LCH-LCH14430/2025 | 2024-2025 | Jan-2025 | North America / USA / Illinois | pediatric | Nasopharyngeal | B.D.E.1.2 | PZ279525 |
| LCH14322 | RSV-B/human/USA/IL-LCH-LCH14322/2025 | 2024-2025 | Jan-2025 | North America / USA / Illinois | pediatric | Nasopharyngeal | B.D.E.1 | |
| LCH14774 | RSV-B/human/USA/IL-LCH-LCH14774/2025 | 2024-2025 | Jan-2025 | North America / USA / Illinois | pediatric | Nasopharyngeal | B.D.E.1.2 | PZ279529 |
| LCH14873 | RSV-B/human/USA/IL-LCH-LCH14873/2025 | 2024-2025 | Jan-2025 | North America / USA / Illinois | pediatric | Nasopharyngeal | B.D.E.1.2 | PZ279530 |
| LCH14877 | RSV-B/human/USA/IL-LCH-LCH14877/2025 | 2024-2025 | Jan-2025 | North America / USA / Illinois | pediatric | Nasopharyngeal | B.D.E.1 | PZ279531 |
| LCH14718 | RSV-B/human/USA/IL-LCH-LCH14718/2025 | 2024-2025 | Jan-2025 | North America / USA / Illinois | pediatric | Nasopharyngeal | B.D.E.1.2 | PZ279526 |
| LCH14733 | RSV-B/human/USA/IL-LCH-LCH14733/2025 | 2024-2025 | Jan-2025 | North America / USA / Illinois | pediatric | Nasopharyngeal | B.D.E.1 | PZ279527 |
| LCH14754 | RSV-B/human/USA/IL-LCH-LCH14754/2025 | 2024-2025 | Jan-2025 | North America / USA / Illinois | pediatric | Nasopharyngeal | B.D.E.1.2 | PZ279528 |
| LCH15172 | RSV-B/human/USA/IL-LCH-LCH15172/2025 | 2024-2025 | Jan-2025 | North America / USA / Illinois | pediatric | Nasopharyngeal | B.D.E.1.1 | |

| | | | | | | | | |
|---|---|---|---|---|---|---|---|---|
| LCH15274 | RSV-B/human/USA/IL-LCH-LCH15274/2025 | 2024-2025 | Jan-2025 | North America / USA / Illinois | pediatric | Nasopharyngeal | B.D.E.1.2 | PZ279534 |
| LCH15209 | RSV-B/human/USA/IL-LCH-LCH15209/2025 | 2024-2025 | Jan-2025 | North America / USA / Illinois | pediatric | Nasopharyngeal | B.D.E.1.2 | PZ279533 |
| LCH15355 | RSV-B/human/USA/IL-LCH-LCH15355/2025 | 2024-2025 | Jan-2025 | North America / USA / Illinois | pediatric | Nasopharyngeal | B.D.E.1 | |
| LCH15168 | RSV-B/human/USA/IL-LCH-LCH15168/2025 | 2024-2025 | Jan-2025 | North America / USA / Illinois | pediatric | Nasopharyngeal | B.D.E.1.2 | PZ279532 |
| LCH15440 | RSV-B/human/USA/IL-LCH-LCH15440/2025 | 2024-2025 | Feb-2025 | North America / USA / Illinois | pediatric | Nasopharyngeal | B.D.E.1.2 | PZ279535 |
| LCH15787 | RSV-B/human/USA/IL-LCH-LCH15787/2025 | 2024-2025 | Feb-2025 | North America / USA / Illinois | pediatric | Nasopharyngeal | B.D.E.1.2 | PZ279537 |
| LCH15786 | RSV-B/human/USA/IL-LCH-LCH15786/2025 | 2024-2025 | Feb-2025 | North America / USA / Illinois | pediatric | Nasopharyngeal | B.D.E.1.2 | PZ279536 |
| LCH16102 | RSV-B/human/USA/IL-LCH-LCH16102/2025 | 2024-2025 | Feb-2025 | North America / USA / Illinois | pediatric | Nasopharyngeal | B.D.E.1 | PZ279539 |
| LCH16057 | RSV-B/human/USA/IL-LCH-LCH16057/2025 | 2024-2025 | Feb-2025 | North America / USA / Illinois | pediatric | Nasopharyngeal | B.D.E.1.2 | PZ279538 |
| LCH16284 | RSV-B/human/USA/IL-LCH-LCH16284/2025 | 2024-2025 | Mar-2025 | North America / USA / Illinois | pediatric | Nasopharyngeal | B.D.E.1.2 | PZ279540 |

**Supplementary Table 2 | NCBI accession ID numbers for deposited RSV whole genome sequences.** *Note several sequences are in queue for NCBI sequence submission.

| Step | Primer Sequence |
| --- | --- |
| RT | ACG CGA AAA AAT GCG TAC |
| RT | AAYAAAGGAGCATTCAAATA |
| RT | AAG GKG AAC CWA TAA TAA ATT |
| RT | TATACTATGTMAACAAGCTG |
| RT | TATATTATGTAAATAAGCAAG |
| RT | GAC CAT WGA AGC YAT ATC A |
| PCR - Forward (pair 1) | ACGCGAAAAAATGCGTACWAC |
| PCR - Forward (pair 2) | GCCACARAGTCAATTYATAGTAG |
| PCR - Forward (pair 3A & B) | TGATGCATCAATATCTCAAGTC |
| PCR - Forward (pair 4) | GAGATATGCARTTYATGAGYA |
| PCR - Reverse (pair 1) | TTTGATTGMAAAWCGTGTAGCTG |
| PCR - Reverse (pair 2) | TGTRACTGGTGTGYTTYTGG |
| PCR - Reverse (pair 3A) | AGGACTTTCTTTATACTAGCTG |
| PCR - Reverse (pair 3B) | AGGACTTTTTTGATACTGGCTG |
| PCR - Reverse (pair 4) | TGRATTTAAACTTATTCTTCCTAGA |

**Supplementary Table 3 | Primer information for RSV whole genome sequencing pipeline.**

| Primer | Sequence (5' — 3') |
|---|---|
| gBlock RSV-B FL lab strain B1 Fusion | GGTACCGAGCTCGGATCCACTGCCACCATGGAGCTTCTCATCCATCGACTGTCCGCAATTTTCCTCACTCTTGCAATTAACG<br>CCCTGTACCTGACCTCCAGTCAGAATATAACAGAGGAGTTCTATCAGTCAACGTGTAGTGCCGTTTCAAGAGGTTACTTCAG<br>TGCGCTCCGGACCGGGTGGTATACCAGCGTAATCACAATTGAGCTGAGTAATATCAAGGAGACTAAGTGTAATGGCACTGA<br>CACTAAGGTCAAGCTCATCAAACAGGAGCTCGACAAATATAAAAACGCCGTAACGGAGCTCCAATTGCTTATGCAAAACAC<br>CCCTGCAGCTAATAACCGAGCAAGACGAGAAGCACCTCAGTATATGAATTATACTATCAACACTACCAAGAATCTGAATGT<br>CTCCATATCTAAGAAGCGAAAACGGCGCTTTCTTGGCTTTCTCTTGGGGGTCGGATCTGCAATAGCTAGTGGAATAGCAGTT<br>TCTAAGGTTTTGCACCTGGAAGGAGAGGTCAATAAGATTAAGAACGCGCTTTTGTCTACTAACAAAGCTGTTGTTTCACTGT<br>CTAATGGCGTGTCTGTTCTGACAAGCAAGGTCTTGGATCTGAAAAATTACATTAATAATCAGTTGCTCCCCATTGTGAACCA<br>ACAAAGTTGTCGAATTAGTAACATTGAGACCGTCATAGAGTTTCAGCAAAAGAACTCAAGGCTGCTCGAGATCAATCGAGA<br>ATTTTCAGTAAATGCCGGGGTTACGACCCCCCTCAGTACGTACATGCTCACGAACTCCGAACTCCTCTCTTTGATTAATGAT<br>ATGCCTATAACGAACGATCAGAAGAAACTGATGAGCAGCAATGTGCAAATAGTGAGGCAACAGTCATATTCCATAATGTCC<br>ATTATTAAGGAGGAGGTTCTTGCCTACGTCGTCCAGCTGCCGATCTATGGAGTCATTGACACTCCTTGCTGGAAACTGCATA<br>CTAGTCCACTCTGCACGACTAATATTAAAGAAGGGTCCAACATATGTTTGACACGAACCGATAGGGGTTGGTACTGCGACA<br>ATGCCGGTTCTGTTAGTTTTTTCCCCCAAGCCGACACCTGTAAGGTCCAGTCTAATCGAGTATTTTGTGATACAATGAACAGT<br>CTCACGCTGCCTTCAGAAGTGTCTTTGTGCAACACGGATATTTTTAATTCTAAGTATGATTGTAAAATAATGACATCTAAGAC<br>TGACATCAGTAGCTCAGTAATAACCTCTTTGGGGGCTATTGTTAGTTGTTACGGTAAGACTAAGTGTACAGCTAGTAATAAG<br>AACAGGGGAATAATCAAGACATTCTCTAACGGCTGCGATTACGTCTCTAATAAGGGCGTGGACACGGTTTCTGTTGGCAAT<br>ACACTTTATTATGTGAACAAATTGGAAGGCAAAAACCTGTACGTCAAGGGTGAGCCGATAATTAATTACTATGATCCACTCG<br>TATTTCCATCAGACGAGTTCGACGCCAGTATTAGTCAAGTAAATGAGAAGATCAACCAAAGTCTTGCATTTATCCGAAGGTC<br>CGACGAACTGCTTCATAATGTGAATACAGGGAAGAGCACGACGAACATTATGATTACGACGATAATTATAGTGATAATAGT<br>CGTACTTTTGTCTTTGATAGCCATAGGATTGCTTCTCTACTGTAAAGCCAAAAACACCCCCGTGACTCTGTCAAAAGATCAA<br>CTCAGTGGTATAAACAATATCGCCTTTTCAAAATAGTGGAATTCTGCAGATATΩCCAG |
| gBlock RSV-A FL A.D.5.2 Fusion | ggtaccgagctcggatccactgccaccATGGAACTCCCGATCCTGAAGACTAACGCAATCACAACAATTCTTGCGGCAGTAACACTGT<br>GTTTCGCTTCCTCTCAGAACATCACGGAAGAATTTTACCAGTCAACATGCTCTGCGGTGAGCAAAGGCTATCTTTCTGCACT<br>CAGGACCGGATGGTACACATCCGTTATTACCATCGAATTGAGTAATATCAAGGAAAACAAGTGTAATGGGACGGATGCTAA<br>AGTGAAGTTGATTAAGCAAGAGCTTGATAAGTACAAAAATGCCGTAACGGAGCTCCAGCTCCTGATGCAATCAACCCCCGC<br>TACAAATTCTAGGGCTAGAAGGGAGTTGCCACGCTTCATGAATTACACGCTTAATAATGCTAAAAATACTAACGTTACTCTT<br>AGTAAGAAGAGAAAGCGACGGTTTCTCGGATTCCTGTTGGGGGTAGGAAGCGCAATCGCTAGTGGCATCGCGGTAAGCAA<br>AGTTTTGCATCTCGAAGGGGAGGTCAACAAGATTAAGTCCGCTCTTTTGTCTACTAATAAGGCGGTGGTCTCTCTCTCTAAC<br>GGTGTAAGTGTTCTGACCAGCAAAGTGCTCGACCTCAAGAACTACATAGACAAACAGCTTCTGCCCATTGTAAACAAGCAA<br>AGCTGCAGCATTAGTAACATTGAGACTGTCATTGAGTTCCAGCAGAAAAACAACAGGCTGCTTGAAATCACCCGCGAGTTT<br>AGCGTAAACGCCGGAGTAACCACGCCTGTAAGTACCTACATGCTTACTAATAGTGAGCTGTTGTCACTCATCAACGATATGC<br>CCATTACCAATGATCAAAAGAAGCTCATGAGCAGCAATGTGCAGATAGTACGGCAGCAAAGCTATAGTATAATGTCAATAA<br>TAAAGGAAGAGGTTTTGGCATACGTCGTGCAGCTGCCCCTTTATGGAGTTATTGATACACCCTGCTGGAAATTGCACACGA<br>GCCCTTTGTGCACGACCAACACTAAAGAAGGGTCTAACATCTGCCTTACTCGCACAGACAGAGGATGGTACTGCGACAACG<br>CAGGATCTGTGAGTTTTTTTCCACAAGCAGAAACCTGTAAGGTCCAATCAAATCGAGTCTTCTGCGATACCATGAACAGTCT<br>TACTCTTCCTAGCGAAGTAAACCTGTGCAATATAGATATATTTAACCCCAAATATGATTGCAAAATTATGACTTCCAAAACGG<br>ATGTATCTTCCTCAGTAATAACGTCACTCGGGGCAATAGTTTCCTGCTATGGTAAGACTAAATGTACAGCCTCCAACAAAAA<br>TAGGGGCATAATAAAAACCTTTAGCAATGGATGCGATTACGTTTCAAATAAGGGTGTTGATACTGTGTCAGTTGGCAACACG<br>CTTTACTACGTGAACAAGCAAGAAGGGAAAAGTCTGTATGTTAAGGGGGAACCGATTATTAACTTCTATGATCCTCTTGTGT<br>TCCCCTCTGACGAGTTCGACGCCTCAATTAGCCAAGTTAACGAGAAGATAAACCAAAGTCTTGCTTTTATCCGAAAATCAGA<br>TGAATTGCTTCATAACGTGAATGCAGGCAAATCCACTACTAATATTATGATTACGACTATTATCATCGTAATCATTGTAATCC<br>TTTTGGCTCTCATAGCAGTCGGGCTTTTGCTCTACTGTAAAGCACGATCCACCCCCGTCACACTGTCCAAAGATCAGCTCTC<br>TGGGATAAATAACATTGCCTTTAGTAATTGAtggaattctgcagatatccag |

| | |
|---|---|
| gBlock RSV-B FL B.D.E.1 Fusion | GGTACCGAGCTCGGATCCACTGCCACCATGGAATTGCTCATCCACCGCTCTTCTGCTATCTTTTTGACCCTCGCGATAAATG<br>CCCTGTATCTGACTTCCAGCCAGAACATTACAGAAGAGTTCTACCAGTCTACCTGCTCAGCTGTGTCTAGGGGCTACCTCTC<br>TGCTTTGAGAACAGGATGGTACACTTCAGTGATCACAATCGAGCTGAGCAACATAAAGGAAACTAAATGTAACGGGACAGA<br>CACGAAAGTGAAGCTGATAAAACAAGAATTGGATAAGTATAAGAACGCTGTCACGGAATTGCAGCTTCTCATGCAGAACAC<br>ACCGGCGGTAAATAATAGAGCTAGACGAGAGGCCCCGCAATATATGAACTACACAATTAACACTACAAAGAACCTCAACGT<br>ATCAATCTCCAAGAAAAGAAAGCGCCGCTTTCTTGGTTTCCTCCTGGGAGTAGGAAGCGCCATAGCGTCTGGCATAGCGGT<br>AAGTAAAGTTCTTCACTTGGAGGGGGAGGTTAACAAGATAAAGAATGCATTGCAACTCACGAATAAGGCGGTGGTTTCCCT<br>GAGTAATGGAGTCTCCGTACTGACTAACAGAGTACTTGACCTGAAAAACTATATTAACAACCAACTGCTGCCGATGGTCAAT<br>CGACAAAATTGTAGGATATCAAATATAGAAACAGTGATCGAATTTCAGCAAAAAAATAGCCGACTGCTCGAAATTACACGA<br>GAGTTTTCAGTTAATGCCGGTGTCACGACTCCACTCAGCACTTATATGCTCACAAATAGCGAACTCTTGTCTCTGATAAACG<br>ACATGCCTATAACGAACGATCAAAAAAAACTGATGAGCTCCAACGTTCAAATCGTCCGCCAACAGAGCTATTCTATCATGA<br>GCATTATAAAGGAAGAAGTACTTGCTTATGTAGTTCAGCTGCCGATATACGGAGTAATTGATACTCCGTGTTGGAAACTGCA<br>TACGAGTCCTCTCTGTACGACGAACATAAAAGAAGGGAGCAATATATGTCTGACACGAACTGATCGGGGTTGGTATTGTGA<br>TAACGCAGGCAGCGTATCATTCTTTCCACAGGCTGATACTTGCAAAGTTCAAAGTAACAGGGTGTTTTGCGATACGATGAAT<br>AGTCTCACGCTTCCAAGCGAGGTTTCTCTTTGTAATACGGATATTTTCAATCCAAAATACGACTGTAAAATTATGACATCCAA<br>GACAGATATTAGCAGTAGCGTCATTACGTCCCTTGGGGCTATAGTCAGTTGTTACGGCAAGACAAAATGTACGGCTTCAAA<br>CAAAAATCGGGGAATCATAAAAACCTTTTCCAACGGTTGTGACTACGTCTCCAATAAAGGTGTTGATACGGTCTCTGTCGGT<br>AATACTCTGTATTATGTCAATAAACTTGAGGGTAAAAATTTGTATGTCAAAGGAGAGCCCATCATTAATTACTACGACCCGTT<br>GGTGTTTCCCAGTGATGAGTTTGATGCGTCTATTTCTCAGGTAAACGAGAAGATCAACCAGAGCCTGGCCTTCATCAGACGA<br>TCTGATGAATTGCTCCATAATGTCAACACCGGTAAATCAACGACGAATATCATGATAACCGCTATCATTATAGTTATCATAGT<br>TGTATTGCTGTCACTTATCGCCATAGGTCTTCTGCTTTACTGCAAAGCGAAGAATACACCCGTGACTCTTTCCAAGGACCAA<br>CTCTCCGGCATCAACAATATCGCTTTTAGCAAGTAGTGGAATTCTGCAGATATCCAG |
| gBlock RSV-A FL A.D.5.2 Glycoprotein | ggtaccgagctcggatccactgccaccATGAGTAAAACCAAGGACCAGCGAACGGCCAAAACCCTCGAGCGAACTTGGGACACCCTC<br>AATCACCTGCTGTTCATTTCATCCTGTTTGTACAAGCTCAACCTTAAGTCCATAGCGCAGATAACGCTGTCCATACTCGCAAT<br>GATAATCAGCACCTCTCTTATAATTGTTGCAATCATATTCATAGCCTCCGCAAACCATAAGGTGACTCTCACAACAGCGATA<br>ATCCAGGATGCTACGAATCAGATCAAAAATACAACGCCTACATACTTGACGCAAAATCCGCAATTGGGTATTTCCTTTTCTA<br>ACCTCAGTGGGACTACTAGCCAAAGCACGACCATCCTTGCGTCTACTACTCCTTCTGCTGAATCAACGCCTCAGTCTACCAC<br>GGTCAAAATCAAGAACACAACCACGACTCAAATACTCCCTTCTAAGCCGACGACGAAACAAAGGCAAAACAAGCCGCAGA<br>ATAAACCTAACAACGACTTCCACTTCGAAGTTTTTAATTTTGTACCATGTAGTATATGCAGCAACAACCCCACTTGTTGGGCT<br>ATTTGTAAGCGAATCCCCAATAAGAAACCTGGGAAAAAGACGACCACGAAGCCGACCAAGAAACAAACGCTGCGGACTAC<br>CAAGAAAGACCCAAAACCTCAAACGACGAAGCCAAAAGAAGTACTCACCACTAAGCCCACCGGCAAGCCCACGATAAATA<br>CAACTAAAACTAACATTCGCACCACACTTATCACTAGTAACACCAAGGGAAATCCTGAGCACACATCCCAAGAGGAAACAT<br>TGCATTCCACAACAAGTGAAGGCTACCCTAGCCCGTCCCAAGTCTATACTACTAGCGGACAAGAGGAGACTCTCCATTCTA<br>CAACGAGCGAAGGATACCCCAGCCCCAGCCAAGCCCATACCACCTCAGAATACCCGAGCCAGAGCCTTAGTAGCTCCAAC<br>ATTGCAAAATGAtggaattctgcagatatccag |
| gBlock RSV-B FL B.D.E.1 Glycoprotein | ggtaccgagctcggatccactgccaccATGAGTAAAACCAAGGACCAGCGAACGGCCAAAACCCTCGAGCGAACTTGGGACACCCTC<br>AATCACCTGCTGTTCATTTCATCCTGTTTGTACAAGCTCAACCTTAAGTCCATAGCGCAGATAACGCTGTCCATACTCGCAAT<br>GATAATCAGCACCTCTCTTATAATTGTTGCAATCATATTCATAGCCTCCGCAAACCATAAGGTGACTCTCACAACAGCGATA<br>ATCCAGGATGCTACGAATCAGATCAAAAATACAACGCCTACATACTTGACGCAAAATCCGCAATTGGGTATTTCCTTTTCTA<br>ACCTCAGTGGGACTACTAGCCAAAGCACGACCATCCTTGCGTCTACTACTCCTTCTGCTGAATCAACGCCTCAGTCTACCAC<br>GGTCAAAATCAAGAACACAACCACGACTCAAATACTCCCTTCTAAGCCGACGACGAAACAAAGGCAAAACAAGCCGCAGA<br>ATAAACCTAACAACGACTTCCACTTCGAAGTTTTTAATTTTGTACCATGTAGTATATGCAGCAACAACCCCACTTGTTGGGCT<br>ATTTGTAAGCGAATCCCCAATAAGAAACCTGGGAAAAAGACGACCACGAAGCCGACCAAGAAACAAACGCTGCGGACTAC<br>CAAGAAAGACCCAAAACCTCAAACGACGAAGCCAAAAGAAGTACTCACCACTAAGCCCACCGGCAAGCCCACGATAAATA<br>CAACTAAAACTAACATTCGCACCACACTTATCACTAGTAACACCAAGGGAAATCCTGAGCACACATCCCAAGAGGAAACAT<br>TGCATTCCACAACAAGTGAAGGCTACCCTAGCCCGTCCCAAGTCTATACTACTAGCGGACAAGAGGAGACTCTCCATTCTA<br>CAACGAGCGAAGGATACCCCAGCCCCAGCCAAGCCCATACCACCTCAGAATACCCGAGCCAGAGCCTTAGTAGCTCCAAC<br>ATTGCAAAATGAtggaattctgcagatatccag |
| F primer RSV-A2 Fusion 4 aa truncation | CTTGGTACCGAGCTCGGATCCACTAGCCACCATGGAACTGCCGATTCTGA |
| R primer RSV-A2 Fusion 4 aa truncation | GCTGGATATCTGCAGAATTCCACCATCAGATGTTGTTGATGCCGCTCAGC |
| F primer RSV-A2 Glycoprotein 31 aa truncation | CTTGGTACCGAGCTCGGATCCACTAgccaccatgcttaacctgaagtctg |
| R primer RSV-A2 Glycoprotein 31 aa truncation | GCTGGATATCTGCAGAATTCCACCAttacttatcatcgtcgtccttataa |
| F primer RSV-A A.D.5.2 Glycoprotein 31 aa truncation | GCTTGGTACCGAGCTCGgatccactgccaccATGCTCAACCTTAAGTCCA |
| R primer RSV-A A.D.5.2 Glycoprotein 31 aa truncation | TGTGCTGGATATCTGCAGAATTccaTCATTTTGCAATGTTGGAGCTACTA |

| | |
|---|---|
| **F primer RSV-B B.D.E.1 Glycoprotein 31 aa truncation** | **CTTGGTACCGAGCTCGGATCCACTAgccaccatgcttaacttgaaatcta** |
| **R primer RSV-B B.D.E.1 Glycoprotein 31 aa truncation** | **GCTGGATATCTGCAGAATTCCACCAttatatagaattgcttgtgctaggc** |
| **F primer RSV-A A.D.5.2 Fusion 4 aa truncation** | **GCTTGGTACCGAGCTCGgatccactgccaccATGGAACTCCCGATCCTGA** |
| **R primer RSV-A A.D.5.2 Fusion 4 aa truncation** | **TGTGCTGGATATCTGCAGAATTccaTCAAATGTTATTTATCCCAGAGAGC** |
| **F primer RSV-B1 Fusion 4 aa truncation** | **CTTGGTACCGAGCTCGGATCCACTAGCCACCATGGAGCTTCTCATCCATC** |
| **R primer RSV-B1 Fusion 4 aa truncation** | **GCTGGATATCTGCAGAATTCCACCACTAGATATTGTTTATACCACTGAGT** |
| **F primer on RSV-A tr A.D.5.2 (T103A) to create RSV-A tr A.D.5.1** | **CAGCTCCTGATGCAATCAACCCCCGCTGCAAATTCTAGGGCTAGAAGGGAGTTGCCACG** |
| **R primer on RSV-A tr A.D.5.2 (T103A) to create RSV-A tr A.D.5.1** | **CGTGGCAACTCCCTTCTAGCCCTAGAATTTGCAGCGGGGGTTGATTGCATCAGGAGCTG** |
| **F primer on RSV-A tr A.D.5.1 (A122T) to create RSV-A tr A.D.1** | **CACGCTTCATGAATTACACGCTTAATAATACTAAAAATACTAACGTTACTCTTAGTAAG** |
| **R primer on RSV-A tr A.D.5.1 (A122T) to create RSV-A tr A.D.1** | **CTTACTAAGAGTAACGTTAGTATTTTTAGTATTATTAAGCGTGTAATTCATGAAGCGTG** |
| **F primer on RSV-A tr A.D.1 (T12I) to create RSV-A tr A.D.3** | **AACTCCCGATCCTGAAGACTAACGCAATCATAACAATTCTTGCGGCAGTAACACTGTGT** |
| **R primer on RSV-A tr A.D.1 (T12I) to create RSV-A tr A.D.3** | **ACACAGTGTTACTGCCGCAAGAATTGTTATGATTGCGTTAGTCTTCAGGATCGGGAGTT** |
| **F primer on RSV-A tr A.D.5.1 (L15F) to create RSV-A tr A.D.5.3** | **CCTGAAGACTAACGCAATCACAACAATTTTTGCGGCAGTAACACTGTGTTTCGCTTCCT** |
| **R primer on RSV-A tr A.D.5.1 (L15F) to create RSV-A tr A.D.5.3** | **AGGAAGCGAAACACAGTGTTACTGCCGCAAAAATTGTTGTGATTGCGTTAGTCTTCAGG** |
| **F primer on RSV-B tr B.D.E.1 (F12I) to create RSV-B tr B.D.E.1.4 & tr B.D.E.2** | **ATTGCTCATCCACCGCTCTTCTGCTATCATTTTGACCCTCGCGATAAATGCCCTGTATC** |
| **R primer on RSV-B tr B.D.E.1 (F12I) to create RSV-B tr B.D.E.1.4 & tr B.D.E.2** | **GATACAGGGCATTTATCGCGAGGGTCAAAATGATAGCAGAAGAGCGGTGGATGAGCAAT** |
| **F primer on RSV-B tr B.D.E.1 (N190S) to create RSV-B tr B.D.4.1.1** | **CCCTGAGTAATGGAGTCTCCGTACTGACTAGCAGAGTACTTGACCTGAAAAACTATATT** |
| **R primer on RSV-B tr B.D.E.1 (N190S) to create RSV-B tr B.D.4.1.1** | **AATATAGTTTTTCAGGTCAAGTACTCTGCTAGTCAGTACGGAGACTCCATTACTCAGGG** |
| **F primer on RSV-B tr B.D.4.1.1 N190S (N211S) to create RSV-B tr B.D.4.1.1** | **ACCAACTGCTGCCGATGGTCAATCGACAAAGTTGTAGGATATCAAATATAGAAACAGTG** |
| **R primer on RSV-B tr B.D.4.1.1 N190S (N211S) to create RSV-B tr B.D.4.1.1** | **CACTGTTTCTATATTTGATATCCTACAACTTTGTCGATTGACCATCGGCAGCAGTTGGT** |
| **F primer on RSV-B tr B.D.4.1.1 N190S N211S (P389S) to create RSV-B tr B.D.4.1.1** | **TTTCTCTTTGTAATACGGATATTTTCAATTCAAAATACGACTGTAAAATTATGACATCC** |
| **R primer on RSV-B tr B.D.4.1.1 N190S N211S (P389S) to create RSV-B tr B.D.4.1.1** | **GGATGTCATAATTTTACAGTCGTATTTTGAATTGAAAATATCCGTATTACAAAGAGAAA** |
| **F primer on RSV-A tr A.D.5.2 (T8A) to identify rescue mutation** | **tgccaccATGGAACTCCCGATCCTGAAG GCC AACGCAATCACAACAATTCTTGCGGCAG** |
| **R primer on RSV-A tr A.D.5.2 (T8A) to identify rescue mutation** | **CTGCCGCAAGAATTGTTGTGATTGCGTT GGC CTTCAGGATCGGGAGTTCCATGGTGGCA** |
| **F primer on RSV-A tr A.D.5.2 (L20F) to identify rescue mutation** | **AATCACAACAATTCTTGCGGCAGTAACA TTC TGTTTCGCTTCCTCTCAGAACATCACGG** |
| **R primer on RSV-A tr A.D.5.2 (L20F) to identify rescue mutation** | **CCGTGATGTTCTGAGAGGAAGCGAAACA GAA TGTTACTGCCGCAAGAATTGTTGTGATT** |

| | |
|---|---|
| F primer on RSV-A tr A.D.5.2 (T103A) to identify rescue mutation | CCAGCTCCTGATGCAATCAACCCCCGCTgccAATTCTAGGGCTAGAAGGGAGTTGCCAC |
| R primer on RSV-A tr A.D.5.2 (T103A) to identify rescue mutation | GTGGCAACTCCCTTCTAGCCCTAGAATTGGCAGCGGGGGTTGATTGCATCAGGAGCTGG |
| F primer on RSV-A tr A.D.5.2 (S105N) to identify rescue mutation | GATTAAGCAAGAGCTTGATAAGTACAAA AGT GCCGTAACGGAGCTCCAGCTCCTGATGC |
| R primer on RSV-A tr A.D.5.2 (S105N) to identify rescue mutation | GCATCAGGAGCTGGAGCTCCGTTACGGC ACT TTTGTACTTATCAAGCTCTTGCTTAATC |
| F primer on RSV-A tr A.D.5.2 (A122T) to identify rescue mutation | ACGCTTCATGAATTACACGCTTAATAAT ACC AAAAATACTAACGTTACTCTTAGTAAGA |
| R primer on RSV-A tr A.D.5.2 (A122T) to identify rescue mutation | TCTTACTAAGAGTAACGTTAGTATTTTT GGT ATTATTAAGCGTGTAATTCATGAAGCGT |
| F primer on RSV-A tr A.D.5.2 (N124K) to identify rescue mutation | GCTTCATGAATTACACGCTTAATAATGCTAAAAAGACTAACGTTACTCTTAGTAAGAAG |
| R primer on RSV-A tr A.D.5.2 (N124K) to identify rescue mutation | CTTCTTACTAAGAGTAACGTTAGTCTTTTTAGCATTATTAAGCGTGTAATTCATGAAGC |
| F primer on RSV-A tr A.D.5.2 (I152T) to identify rescue mutation | GGGGTAGGAAGCGCAATCGCTAGTGGCaccGCGGTAAGCAAAGTTTTGCATCTCGAAGG |
| R primer on RSV-A tr A.D.5.2 (I152T) to identify rescue mutation | CCTTCGAGATGCAAAACTTTGCTTACCGCGGTGCCACTAGCGATTGCGCTTCCTACCCC |
| F primer on RSV-A tr A.D.5.2 (S213R) to identify rescue mutation | CTGCCCATTGTAAACAAGCAAAGCTGCagaATTAGTAACATTGAGACTGTCATTGAGTT |
| R primer on RSV-A tr A.D.5.2 (S213R) to identify rescue mutation | AACTCAATGACAGTCTCAATGTTACTAATTCTGCAGCTTTGCTTGTTTACAATGGGCAG |
| F primer on RSV-A tr A.D.5.2 (S276N) to identify rescue mutation | TACCAATGATCAAAAGAAGCTCATGAGC AAC AATGTGCAGATAGTACGGCAGCAAAGCT |
| R primer on RSV-A tr A.D.5.2 (S276N) to identify rescue mutation | AGCTTTGCTGCCGTACTATCTGCACA TTG TTGCTCATGAGCTTCTTTTGATCATTGGTA |
| F primer on RSV-A tr A.D.5.2 (I384V) to identify rescue mutation | TTCCTAGCGAAGTAAACCTGTGCAATgtgGATATATTTAACCCCAAATATGATTGCAAA |
| R primer on RSV-A tr A.D.5.2 (I384V) to identify rescue mutation | TTTGCAATCATATTTGGGGTTAAATATATCCACATTGCACAGGTTTACTTCGCTAGGAA |
| F primer on RSV-A tr A.D.5.2 (V442A) to identify rescue mutation | AAAAACCTTTAGCAATGGATGCGATTACgcgTCAAATAAGGGTGTTGATACTGTGTCAG |
| R primer on RSV-A tr A.D.5.2 (V442A) to identify rescue mutation | CTGACACAGTATCAACACCCTTATTTGACGCGTAATCGCATCCATTGCTAAAGGTTTTT |
| F primer on RSV-A tr A.D.5.2 (N515H) to identify rescue mutation | TATCCGAAAATCAGATGAATTGCTTCATcatGTGAATGCAGGCAAATCCACTACTAATA |
| R primer on RSV-A tr A.D.5.2 (N515H) to identify rescue mutation | TATTAGTAGTGGATTTGCCTGCATTCACATGATGAAGCAATTCATCTGATTTTCGGATA |
| F primer on RSV-A tr A.D.5.2 (A540S) to identify rescue mutation | TATCATCGTAATCATTGTAATCCTTTTG AGT CTCATAGCAGTCGGGCTTTTGCTCTACT |

| | |
|---|---|
| R primer on RSV-A tr A.D.5.2 (A540S) to identify rescue mutation | AGTAGAGCAAAAGCCCGACTGCTATGAG ACT CAAAAGGATTACAATGATTACGATGATA |
| F primer on RSV-B tr B.D.E.1 (S8L) to identify rescue mutation | GCCACCATGGAATTGCTCATCCACCGCTTGTCTGCTATCTTTTTGACCCTCGCGATAAA |
| R primer on RSV-B tr B.D.E.1 (S8L) to identify rescue mutation | TTTATCGCGAGGGTCAAAAAGATAGCAGACAAGCGGTGGATGAGCAATTCCATGGTGGC |
| F primer on RSV-B tr B.D.E.1 (L45F) to identify rescue mutation | ACCTGCTCAGCTGTGTCTAGGGGCTACTTCTCTGCTTTGAGAACAGGATGGTACACTTC |
| R primer on RSV-B tr B.D.E.1 (L45F) to identify rescue mutation | GAAGTGTACCATCCTGTTCTCAAAGCAGAGAAGTAGCCCCTAGACACAGCTGAGCAGGT |
| F primer on RSV-B tr B.D.E.1 (V103A) to identify rescue mutation | CAGCTTCTCATGCAGAACACACCGGCGGCAAATAATAGAGCTAGACGAGAGGCCCCGCA |
| R primer on RSV-B tr B.D.E.1 (V103A) to identify rescue mutation | TGCGGGGCCTCTCGTCTAGCTCTATTATTTGCCGCCGGTGTGTTCTGCATGAGAAGCTG |
| F primer on RSV-B tr B.D.E.1 (Q172L + L173S) to identify rescue mutation | AGGTTAACAAGATAAAGAATGCATTGCTATCCACGAATAAGGCGGTGGTTTCCCTGAGT |
| R primer on RSV-B tr B.D.E.1 (Q172L + L173S) to identify rescue mutation | ACTCAGGGAAACCACCGCCTTATTCGTGGATAGCAATGCATTCTTTATCTTGTTAACCT |
| F primer on RSV-B tr B.D.E.1 (N190S and R191K) to identify rescue mutation | TGAGTAATGGAGTCTCCGTACTGACTAGCAAAGTACTTGACCTGAAAAACTATATTAAC |
| R primer on RSV-B tr B.D.E.1 (N190S and R191K) to identify rescue mutation | GTTAATATAGTTTTTCAGGTCAAGTACTTTGCTAGTCAGTACGGAGACTCCATTACTCA |
| F primer on RSV-B tr B.D.E.1 (M206I) to identify rescue mutation | AACTATATTAACAACCAACTGCTGCCGATCGTCAATCGACAAAATTGTAGGATATCAAA |
| R primer on RSV-B tr B.D.E.1 (M206I) to identify rescue mutation | TTTGATATCCTACAATTTTGTCGATTGACGATCGGCAGCAGTTGGTTGTTAATATAGTT |
| F primer on RSV-B tr B.D.E.1 (R209Q) to identify rescue mutation | AACAACCAACTGCTGCCGATGGTCAATCAACAAAATTGTAGGATATCAAATATAGAAAC |
| R primer on RSV-B tr B.D.E.1 (R209Q) to identify rescue mutation | GTTTCTATATTTGATATCCTACAATTTTGTTGATTGACCATCGGCAGCAGTTGGTTGTT |
| F primer on RSV-B tr B.D.E.1 (N211S) to identify rescue mutation | CCAACTGCTGCCGATGGTCAATCGACAAAGTTGTAGGATATCAAATATAGAAACAGTGA |
| R primer on RSV-B tr B.D.E.1 (N211S) to identify rescue mutation | TCACTGTTTCTATATTTGATATCCTACAACTTTGTCGATTGACCATCGGCAGCAGTTGG |
| F primer on RSV-B tr B.D.E.1 (T234N) to identify rescue mutation | AGCAAAAAAAATAGCCGACTGCTCGAAATTAATCGAGAGTTTTCAGTTAATGCCGGTGTC |
| R primer on RSV-B tr B.D.E.1 (T234N) to identify rescue mutation | GACACCGGCATTAACTGAAAACTCTCGATTAATTTCGAGCAGTCGGCTATTTTTTTGCT |
| F primer on RSV-B tr B.D.E.1 (P389S) to identify rescue mutation | TTCTCTTTGTAATACGGATATTTTCAATTCAAAATACGACTGTAAAATTATGACATCCA |
| R primer on RSV-B tr B.D.E.1 (P389S) to identify rescue mutation | TGGATGTCATAATTTTACAGTCGTATTTTGAATTGAAAATATCCGTATTACAAAGAGAA |
| F primer on RSV-B tr B.D.E.1 (A529T) to identify rescue mutation | TAAATCAACGACGAATATCATGATAACCACTATCATTATAGTTATCATAGTTGTATTGC |

| | |
|---|---|
| **R primer on RSV-B tr B.D.E.1 (A529T) to identify rescue mutation** | **GCAATACAACTATGATAACTATAATGATAGTGGTTATCATGATATTCGTCGTTGATTTA** |
| **F primer tr RSV-A2 (clesrovimab) (G446E)** | **CAATGGCTGTGACTATGCGAGCAACAAGGAGGTGGACACAGTGTCTGTGGGCAACACCC** |
| **R primer tr RSV-A2 (clesrovimab) (G446E)** | **GGGTGTTGCCCACAGACACTGTGTCCACCTCCTTGTTGCTCGCATAGTCACAGCCATTG** |
| **F primer tr RSV-A2 (Palivizumab) (N262D)** | **GACCAACTCTGAACTGCTGTCCCTGATAGATGATATGCCAATCACCAATGACCAGAAGA** |
| **R primer tr RSV-A2 (Palivizumab) (N262D)** | **TCTTCTGGTCATTGGTGATTGGCATATCATCTATCAGGGACAGCAGTTCAGAGTTGGTC** |

**Supplementary Table 4 | Primer and oligo design for RSV pseudotyping and Fusion assays.**
